# Cannabinoids accumulate in mouse breast milk and differentially regulate lipid composition and lipid signaling molecules involved in infant development

**DOI:** 10.1101/2021.11.04.467281

**Authors:** Clare T Johnson, Gabriel H Dias de Abreu, Ken Mackie, Hui-Chen Lu, Heather B Bradshaw

## Abstract

Maternal cannabis use during lactation may expose developing infants to cannabinoids (CBs) such as tetrahydrocannabinol (THC) and cannabidiol (CBD). CBs modulate lipid signaling molecules in the central nervous system in age- and cell-dependent ways, but their influence on the lipid composition of breastmilk has yet to be established. This study investigates the effects of THC, CBD, or their combination on milk lipids by analyzing the stomach contents of CD1 mouse pups that have been nursed by dams injected with CBs on postnatal days (PND) 1-10 collected 2 hours after the last injection on PND10. HPLC/MS/MS was used to identify and quantify over 80 endogenous lipid species and cannabinoids in pup stomach contents. We show that CBs differentially accumulate in milk, lead to widespread decreases in free fatty acids, decreases in *N*-acyl methionine species, increases *N*-linoleoyl species, as well as modulate levels of endogenous CBs (eCBs) AEA, 2-AG, and their structural congeners. Our data indicate the passage of CBs to pups through breast milk and that maternal CB exposure alters breast milk lipid compositions.

## 1. Introduction

Nursing moms who report postpartum cannabis use cite physical health and better parenting as reasons for consumption; however, they also report receiving little or conflicting guidance on its perinatal safety [1]. Highlighting a need for better counseling, drugs that act on the cannabinoid (CB) signaling system can have significant effects on infant growth and development. Early THC exposure can influence behavior later in life, as evidenced by significant changes in locomotion at 2 months after a single injection of THC (10mg/kg) on PND10 [2]. The CB1 antagonist SR141716A administered to mice on postnatal days (PND) 2-8 results in starvation via loss of suckling behavior, while THC administration results in significant weight gain [3]. Our labs recently showed that perinatal cannabinoid exposure leads to long-term behavioral changes on repetitive tasks and alters the effect of fluoxetine on immobility time during a forced swim task [4]. Because of the effects of early-life CB exposure and the detection of THC, THC metabolites, and CBD in human breast milk [5-8], it is important to evaluate how CBs given to lactating females transfer into milk and how these levels affect the lipid composition of milk, including endogenous cannabinoids (eCBs) and related signaling lipids.

Independent of exogenous CB content, eCBs are a well-documented constituent of normal breast milk. In breast milk samples from Guatemalan women, arachidonic acid (AA) concentration is the highest of the measured free fatty acids (FFAs) and positively correlated to the eCB 2-arachidonoyl glycerol (2-AG), but not the eCB *N*-arachidonoyl ethanolamine (AEA). Docosahexaenoic acid (DHA) has the second-highest concentration and is positively correlated to 2-docosahexaenoyl glycerol, but not DHEA. Eicosapentaenoic acid (EPA) has the lowest concentration and is not correlated to either corresponding eCB congener [9]. This pattern of FFA concentration remains stable across 2 and 4 weeks postpartum; with the exception that DHA levels are significantly increased in mature milk at 4 weeks postpartum compared to their levels at 2 weeks [10]. These data illustrate that the lipid constituents in breast milk are largely stable in the first postpartum month and have likely evolved to meet the developmental needs of the offspring. However, eCBs and lipids in breast milk are sensitive to the effects of maternal behaviors such as diet; for instance, DHA consumption is positively correlated with its glycerol and ethanolamine derivatives, though changes in AA consumption have no correlation to its eCB derivatives [11]. This suggests that maternal behaviors may alter the lipid composition of their breast milk, potentially in a way that negatively impacts the infant.

Infant growth and development are sensitive to changes in lipid signaling molecules in breast milk. *N*-acyl ethanolamines (NAEs) in milk correlate to indices of infant developmental trajectory, including an increased weight-for-age Z-score when breast milk has low levels of oleoyl ethanolamine (OEA), stearoyl ethanolamine (SEA), and palmitoyl ethanolamine (PEA) [12]. Orally administering AEA to male mice from PND1-21, while they are still nursing, results in significantly larger body weight compared to control animals at PND21 and the weight gain persists even after administration has stopped, being evident on PND150 [13]. The same treatment regime significantly increases hypothalamic CB1 receptors at PND21, but no differences are present at PND150 [14]. Therefore, it appears that the specific lipid composition of breast milk, including lipid signaling molecules, is critical for normal infant development.

Because CBs modulate lipids in the central nervous system (CBD; [15]) including in an age-dependent manner (THC; [16],) we tested the hypothesis that CBs administered to a lactating female would influence the lipid profile of their milk. In specific, this study hypothesizes that chronic treatment of a lactating dam with THC, CBD, or a 1:1 combination of THC and CBD will result in measurable levels of CBs as well as broad changes in eCBs and related lipids in breast milk.

## 2. Methods

### 2.1 Animals and injections

Animal care and procedures were approved by the Indiana University Institutional Care and Use Committee. Trio-housed CD1 dams and bucks were bred. Beginning on PND 1, dams were randomly assigned to receive either 3mg/kg CBD (n=8), 3mg/kg THC (n=10), 3mg/kg CBD+3mg/kg THC (n=9), or vehicle (n=7) (1:1:18 cremophor: ethanol: saline) via i.p. injection once daily for 10 days. Dams and pups were sacrificed via isoflurane followed by rapid cervical dislocation 2hrs after the final injection on PND 10. The stomach contents of pups were then collected, flash frozen in liquid nitrogen, and stored at −80° C until processed for lipid analysis.

### 2.2 Lipid Extraction

On the day of lipid extraction, samples were mixed with 2mL of 100% methanol and an internal standard, 5μL of 1μM deuterium labeled AEA (d8AEA), was added. Samples were incubated on ice in the dark for 30 min before being suspended by sonication and centrifuged at 19,000g at 20°C for 20 min. The resulting lipid-containing supernatants were decanted into 7mL of water to create a ∼25% organic solution. This solution was then partially purified on C18 solid phase extraction columns as previously described [15] and lipids were eluted with 1.5mL of 3 different methanolic concentrations (65, 75, and 100%). Elutions were stored at −80°C until analysis with HPLC/MS/MS.

### 2.3 HPLC/MS/MS

Samples were screened for 86 compounds (see supplemental data for complete list) using an Applied Biosystems API 3000 triple quadrupole mass spectrometer with electrospray ionization (Foster City, CA, USA) using multiple reaction monitoring (MRM) mode as previously described [15, 16]. An injection volume of 20uL from each sample was used.

### 2.4 Data Analysis

Analytes were identified based on retention time and mass/charge ratio when compared to known standards. Analyst software (Applied Biosystems) was used to determine analyte concentrations as previously described [15, 16]. Concentrations were converted to moles/gram for each sample and corrected for extraction efficiency based on each individual sample’s percent recovery of the internal standard d8AEA. Statistical analyses were completed in SPSS Statistics 27 (IBM). One-way ANOVAs followed by Fisher’s Least Significant Difference post-hoc analyses were used to determine statistical differences between each treatment group compared to vehicle across related groups of lipids. Statistical significance for all tests was set at *p* < 0.05, and trending significance at 0.05 < *p* < 0.10.

Descriptive and inferential statistics were used to create heatmaps to visualize changes in lipids compared to vehicle as previously described [15, 16]. Briefly, the direction of changes in the treatment group compared to vehicle are depicted by color, with green representing an increase and orange representing a decrease. Level of significance is shown by color shade, wherein *p* < 0.05 is a dark shade and 0.05 < *p* < 0.1 is a light shade. Direction of the change compared to vehicle is represented by up (increased) or down (decreased) arrows. Effect size is represented by the number of arrows, where 1 arrow corresponds to 1-1.49-fold difference, 2 arrows to a 1.5-1.99-fold difference, 3 arrows to a 2-2.99-fold difference, 4 arrows a 3-9.99-fold difference, and 5 arrows a difference of 10-fold or more. To calculate the effect size of a compound that was significantly higher in the treatment group, the average concentration of the drug group was divided by the average concentration of the vehicle group. To calculate the effect size of a compound that was significantly lower in the treatment group, the same calculation was conducted, except that the inverse of the resulting ratio was used to represent a fold decrease.

## 3. Results

### 3.1 Lipid Detection and Samples

83% of the compounds from the screening library were detected in the milk samples. (*See Supplemental data for full list of compounds detected per treatment group in mole/gram*.) We found that chronic CB exposure substantially changed the lipid composition of the breast milk. Figure 1 illustrates the percentages of those detected compounds that were changed in each treatment group. THC+CBD treatment caused 70% of detected lipids to change relative to vehicle, whereas CBD treatment caused changes in 45.8% and THC caused changes in 61%. This variation was largely the result of differences in the percent of lipids that increased, which was only 10.2% with CBD treatment compared to 28% with THC and 35% with THC+CBD. Across all treatment groups, the percentage of compounds that decreased were more similar, with 35.6% decreased in CBD treatment, 32.2% decreased in THC, and 35% decreased in THC+CBD. Overall, levels of responses to THC alone were in in between levels of responses to CBD alone or THC+CBD.

**Figure 1.**
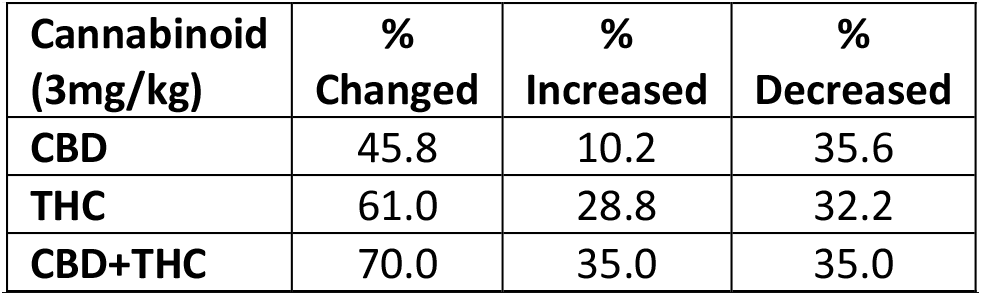
Overall changes in breastmilk lipids recovered from pup stomachs 2 hours after final Cannabinoid injections to dams on P10. Percentages were calculated from the number of individual lipids detected in all samples that were significantly different (either increased or decreased) from vehicle (overall change=% Changed), significantly increased (% Increased), or significantly decreased (% Decreased). Lipids analyzed that were below analytical levels (BAL) or below detectable limits (BDL) were not included in calculations. CBD (Cannabidiol), THC (delta-9 tetrahydrocannabinol)

### 3.2 Cannabinoids

The cannabinoids CBD, THC, and their primary metabolites, 11-OH-THC and 7-OH CBD were detected in each corresponding treatment group. Interestingly, CBD levels were significantly higher in the milk of the THC+CBD group than the CBD alone group (*p*=0.002; Figure 2A). THC levels in the THC+CBD group trended higher than the THC alone group (*p*= 0.104; Figure 2B). CBD levels were significantly higher than THC levels in the THC+CBD group (*p*< 0.001; Figure 2D). Levels of the THC metabolite 11-OH-THC and the CBD metabolite 7-OH-CBD were significantly higher in the THC+CBD group than in either the THC alone or the CBD alone group (*p*< 0.001; Figure 2C-D). THC and CB stock solutions were evaluated by HPLC/MS/MS prior to injection to ensure that they were molecularly equivalent. Furthermore, the leftover injection solutions were also analyzed in conjunction with the milk samples to ensure that any differences were not the result of a difference in injection molarity between the CB injection solutions.

**Figure 2.**
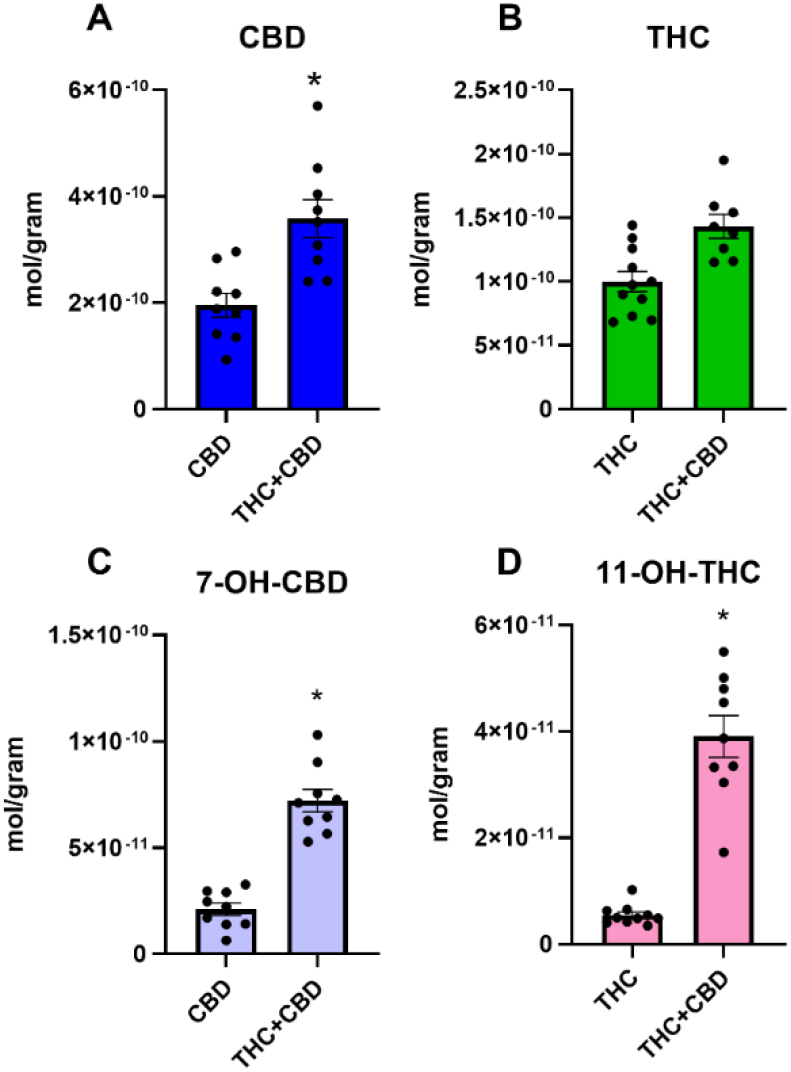
Levels of CBD, THC, and their metabolites in breastmilk recovered from pup stomachs 2 hours after final CB injections to dams on P10. (A) The concentration of CBD was significantly higher in the THC+CBD group than in the CBD alone group (*p=0.002). (B) The concentration of THC trended higher in the THC+CBD group than in the THC alone group (p=0.104). (C) The concentration of CBD metabolite 7-OH-CBD was significantly higher in the THC+CBD group than the CBD alone group (*p<0.001). (D) The concentration of THC metabolite 11-OH-THC was significantly higher in the THC+CBD group than the THC alone group (*p<0.001). Data are presented as mean ±SEM. The individual concentrations of CB measured from each pup are overlaid as scatter. CBD (Cannabidiol), THC (delta-9 tetrahydrocannabinol)

### 3.3 Cannabinoid modulation of endogenous lipids in breast milk more strongly vary by fatty acid chain rather than a specific side-chain conjugate

Figure 3 illustrates the heatmap highlighting significant differences of lipids detected in the milk after chronic CB treatment compared to vehicle control (*see Methods for analytics to generate heatmaps and Supplemental data for moles/gram*). Data are organized by the fatty acid derivative instead of the amine sidechain as in our previous studies [15, 17]. This organization highlights an intriguing finding that the levels of the saturated C18 stearic acid derivatives exclusively decreased after chronic CB treatment (Figure 3B); whereas the levels of the unsaturated (two double bonds) C18 linoleic acid derivatives are almost exclusively increased after chronic CB treatment, though predominately in the THC or THC+CBD groups (Figure 3D). Regarding the oleic acid derivatives (C18, 1 double bond), CB treatments caused changes in both directions (Figure 3C). By contrast, the palmitic acid (saturated C16; 3A) conjugates were highly abundant and, like the OA conjugates, a mosaic of increases and decreases (Figure 3A). Fewer of the AA and DHA conjugates were detected overall as denoted by the “BDL” meaning “below detectable limits”, though there were many that were detected, but not in enough samples to have enough power for statistical analysis and are denoted as “BAL” meaning “below analytical limits” (Figure 3E, F). This overview of the changes in lipid composition in milk after chronic CB exposure in dams suggests CBs dramatically modulate milk lipid metabolism.

**Figure 3.**
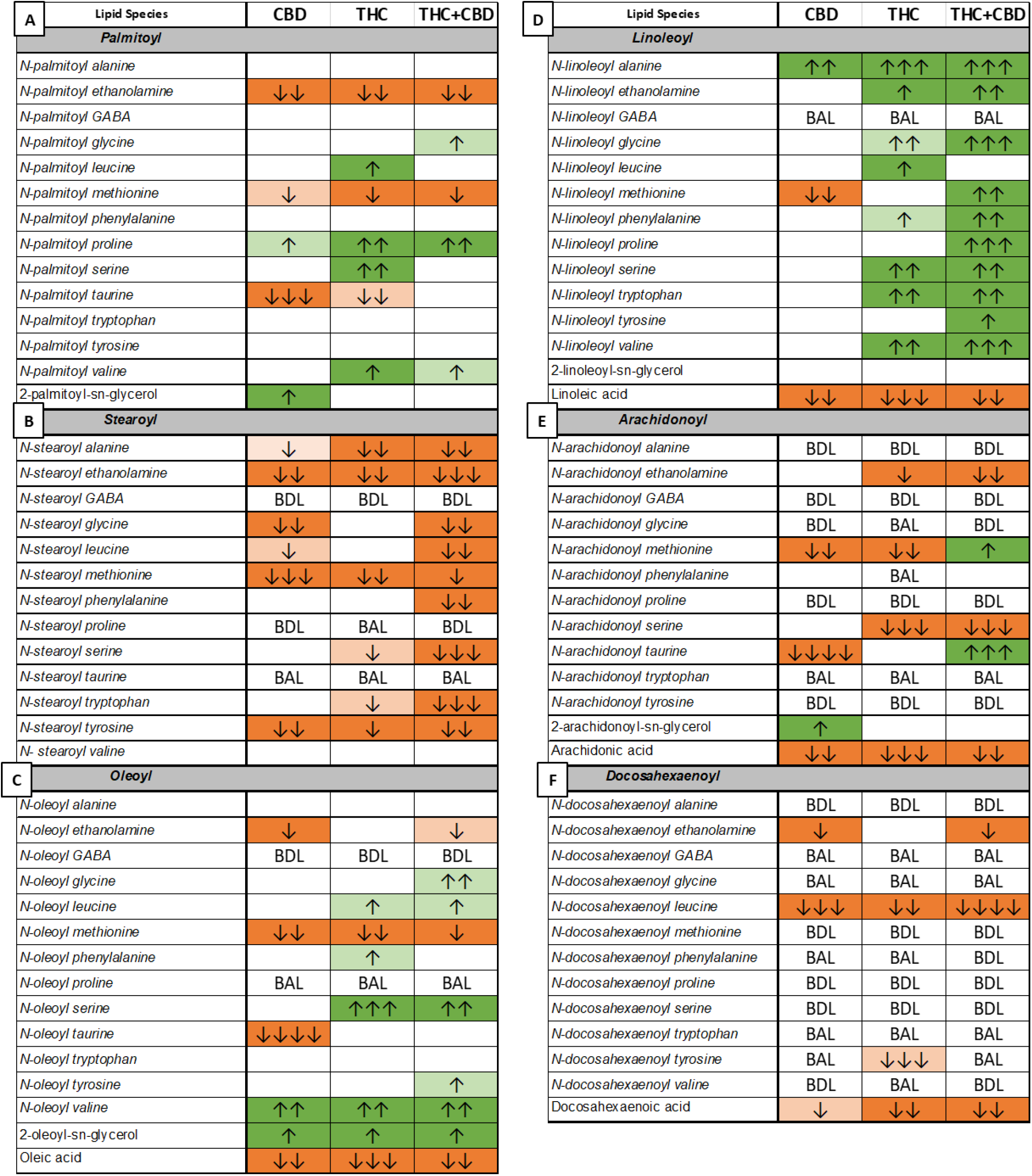
Changes in the milk lipidome compared to vehicle after CBD, THC, or THC+CBD (3mg/kg) administration to lactating dams from P0-P10 collected from pup stomachs 2 hours after final injection to dams on P10. Data are represented as changes with CB treatment compared to the vehicle group and coded as described (*see Methods*). In short, arrows indicate effect size, wherein the more arrows the larger the effect sizer. Green indicates significant increases and orange indicates significant decreases. Lipids that were detected in only a few samples in a treatment group, are labeled “BAL,” below analytical levels. Lipids that were measured in none of the samples are labeled as “BDL” or below detectable limits. (A) Significant and trending differences in the average level of various palmitic acid-derived species relative to vehicle. (B) Significant and trending differences in average concentrations of stearic acid-derived species relative to vehicle. (C) Differences in average concentrations of oleic acid-derived species compared to vehicle. (D) Differences in average concentrations of linoleic acid-derived species compared to vehicle. (E) Differences in average concentrations of arachidonic acid-derived species compared to vehicle. (F) Differences in average concentrations of docosahexaenoic acid-derived species compared to vehicle.

### 3.4 Free Fatty Acids

Figures 4-7 highlight specific lipids from the screen illustrated in Figure 3 as direct quantitative comparisons to provide an additional level of analysis. All CB treatments reduced levels of free fatty acids (FFAs) (Figure 4). The THC treatment group showed the largest effect size, ranging from 1.51 (eicosapentaenoic acid) to 2.2 (arachidonic acid)-fold decreases compared to vehicle. The only FFA that was not significantly decreased was docosahexaenoic acid in the CBD group (*p* = 0.058; Figure 4E). Analytical methods for stearic and palmitic acid are problematic using this shotgun lipidomics approach and were not measured here. It is important to note that the vehicle control animals had a much higher variability across subjects for free fatty acids than those in the CB treatment groups, which are closely clustered together (see scatter plots Figure 4). This may represent a floor effect in the suppression of these lipids with chronic CB treatment.

**Figure 4.**
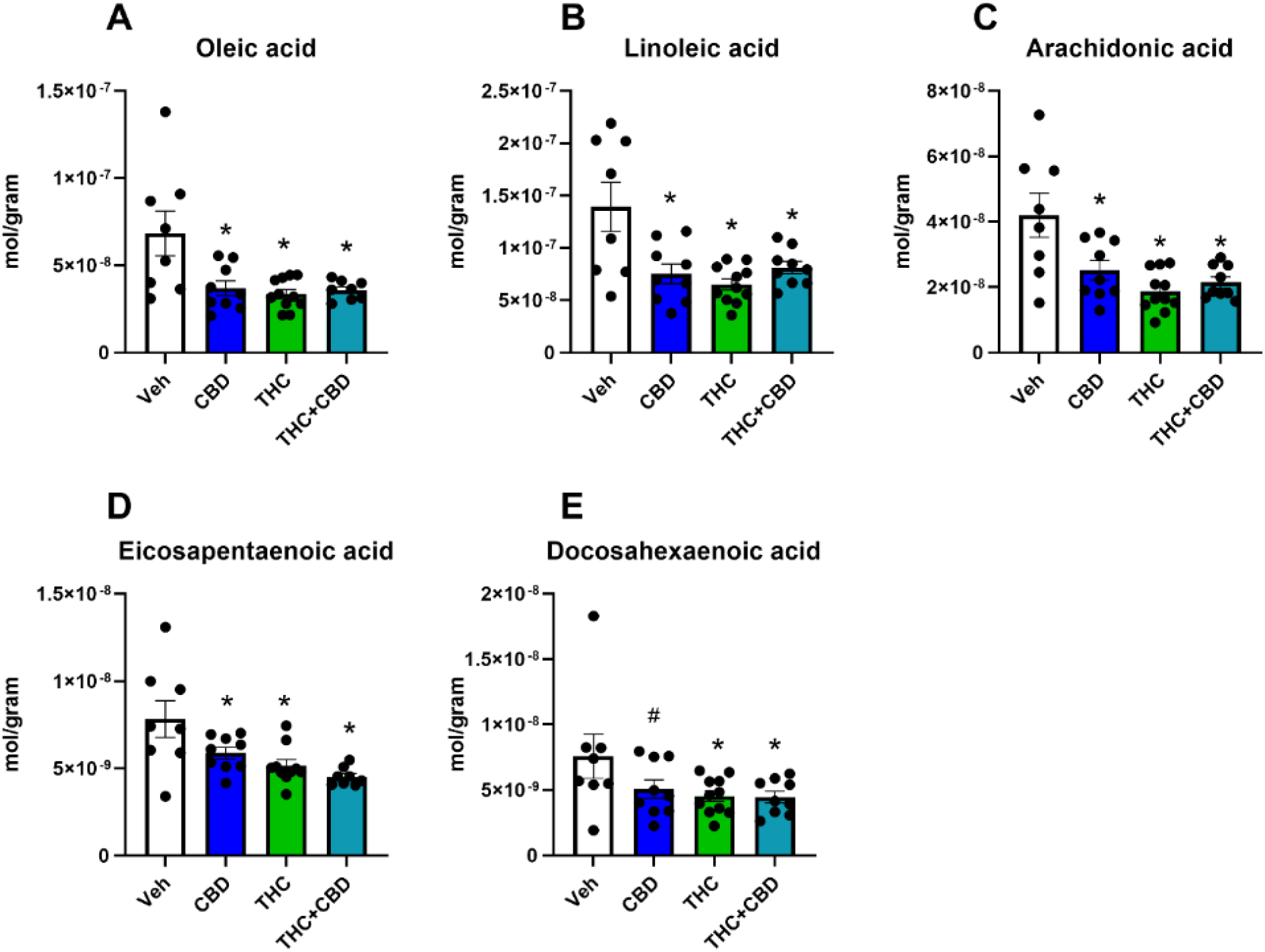
Levels of free fatty acids in milk after CBD, THC, or THC+CBD (3mg/kg) or vehicle administration to lactating dams from P0-P10 collected from pup stomachs 2 hours after final injection to dams on P10. (A) Concentrations of oleic acid in milk. Comparison of vehicle to CBD (p=0.002), THC (p=0.001), or THC+CBD treatment(p=0.002). (B) Concentrations of linoleic acid in milk. Comparison of vehicle to CBD (p=0.001), THC (p=0.001), or THC+CBD treatment(p=0.002). (C) Concentrations of arachidonic acid in milk. Comparison of vehicle to CBD (p=0.003), THC (p=0.001), or THC+CBD treatment(p=0.001). (D) Concentrations of eicosapentaenoic acid in milk. Comparison of vehicle to CBD (p=0.019), THC (p=0.002), or THC+CBD treatment(p=0.001). (E) Concentrations of docosahexaenoic acid in milk. Comparison of vehicle to CBD (p=0.058), THC (p=0.018), or THC+CBD treatment(p=0.020). Data represented as means ±SEM with individual data points from pups plotted as scatter.

### 3. *N*-acyl methionine species

*N*-acyl methionines (Figure 5) are a family of less-studied lipoamines (*aka* lipo amino acids) that are significantly altered by CB treatments. Methionine species were significantly decreased in all treatment groups except for *N*-linoleoyl methionine in the THC group which was unchanged (Figure 5D) and *N*-linoleoyl methionine and *N*-arachidonoyl methionine that were significantly increased in the THC+CBD group (Figure 5D-E).

**Figure 5.**
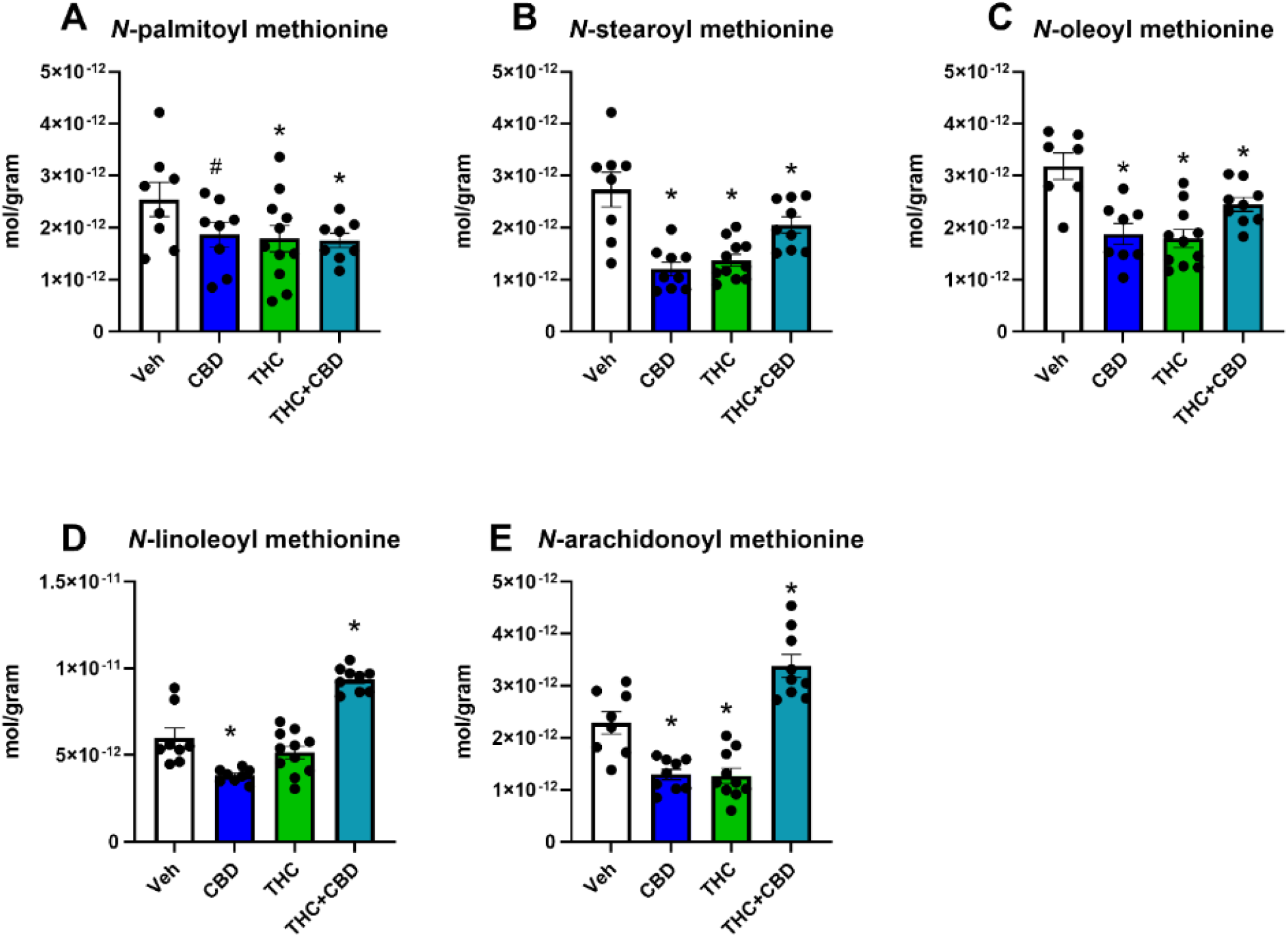
Levels of *N*-acyl methionine in milk after CBD, THC, or THC+CBD (3mg/kg) or vehicle administration to lactating dams from P0-P10 collected from pup stomachs 2 hours after final injection to dams on P10. (A) Concentrations of *N*-palmitoyl methionine in milk. Comparison of vehicle to CBD (p=0.083), THC (p=0.038), or THC+CBD treatment(p=0.044). (B) Concentrations of *N*-stearoyl methionine in milk. Comparison of vehicle to CBD (p=0.001), THC (p=0.001), or THC+CBD treatment(p=0.001). (C) Concentrations of *N*-oleoyl methionine in milk. Comparison of vehicle to CBD (p=0.001), THC (p=0.001), or THC+CBD treatment(p=0.018). (D) Concentrations of *N*-linoleoyl methionine in milk. Comparison of vehicle to CBD (p=0.001), THC (*ns*), or THC+CBD treatment(p=0.001). (E) Concentrations of *N*-arachidonoyl methionine in milk. Comparison of vehicle to CBD (p=0.001), THC (p=0.001), or THC+CBD treatment(p=0.001). Data represented as means ±SEM with individual data points from pups plotted as scatter. *ns*=no significance

### 3.6. 2-acyl-sn-glycerol species including 2-AG

Of the significant differences among 2-AG congeners, all were increases (Figure 6). 2-oleoyl glycerol (2-OG) was increased in all treatment groups compared to vehicle (Figure 6C), whereas 2-AG and 2-palmitoyl glycerol (2-PG) were increased only in the CBD group (Figure 6A, 6B). 2-linoleoyl glycerol (2-LG) was not affected by any CB treatment (Figure 6D).

**Figure 6.**
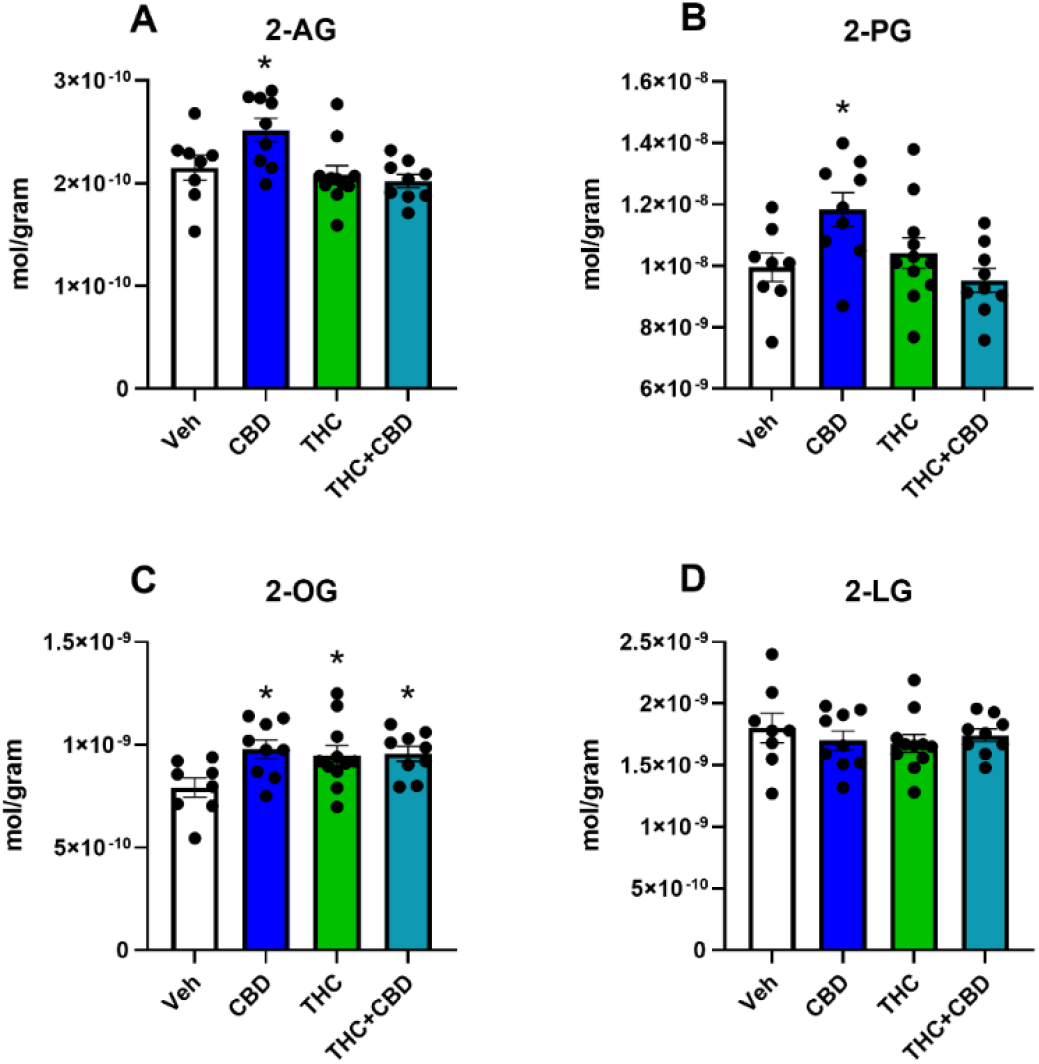
Levels of 2-acyl glycerol in milk after CBD, THC, or THC+CBD (3mg/kg) or vehicle administration to lactating dams from P0-P10 collected from pup stomachs 2 hours after final injection to dams on P10. (A) Concentrations of 2-arachidonoyl-glycerol (2-AG) in milk. Comparison of vehicle to CBD (p=0.018), THC (*ns*), or THC+CBD treatment(*ns*). (B) Concentrations of 2-palmitoyl-glycerol (2-PG) in milk. Comparison of vehicle to CBD (p=0.014), THC (*ns*), or THC+CBD treatment(*ns*). (C) Concentrations of 2-oleoyl-glycerol (2-OG) in milk. Comparison of vehicle to CBD (p=0.009), THC (p=0.020), or THC+CBD treatment(p=0.019). (D) Concentrations of 2-linoleoyl-glycerol (2-LG) in milk. Comparison of vehicle to CBD (*ns*), THC (*ns*), or THC+CBD treatment(*ns*). Data represented as means ±SEM with individual data points from pups plotted as scatter. *ns*=no significance

### 3.5 *N*-acyl ethanolamine species including the endocannabinoid, anandamide

Levels of anandamide (AEA) were significantly lower relative to vehicle in the THC and THD+CBD group only (Figure 7A); whereas the eCB, 2-AG, increased in CBD group only (Figure 6A). Levels of the AEA congeners (*N*-acyl ethanolamines; NAEs), *N*-palmitoyl ethanolamine (PEA) and *N*-stearoyl ethanolamine (SEA) were the only NAEs that were universally affected by all CB treatments, with both being decreased by CBs in all groups (Figure 7B, C). *N*-oleoyl ethanolamine (OEA) and *N*-docosahexaenoyl ethanolamine (DHEA) were significantly lower in CBD and THC+CBD groups only (Figure 7D, F). The only increase in any NAE was LEA that was significantly higher in the THC and THC+CBD groups (7E).

**Figure 7.**
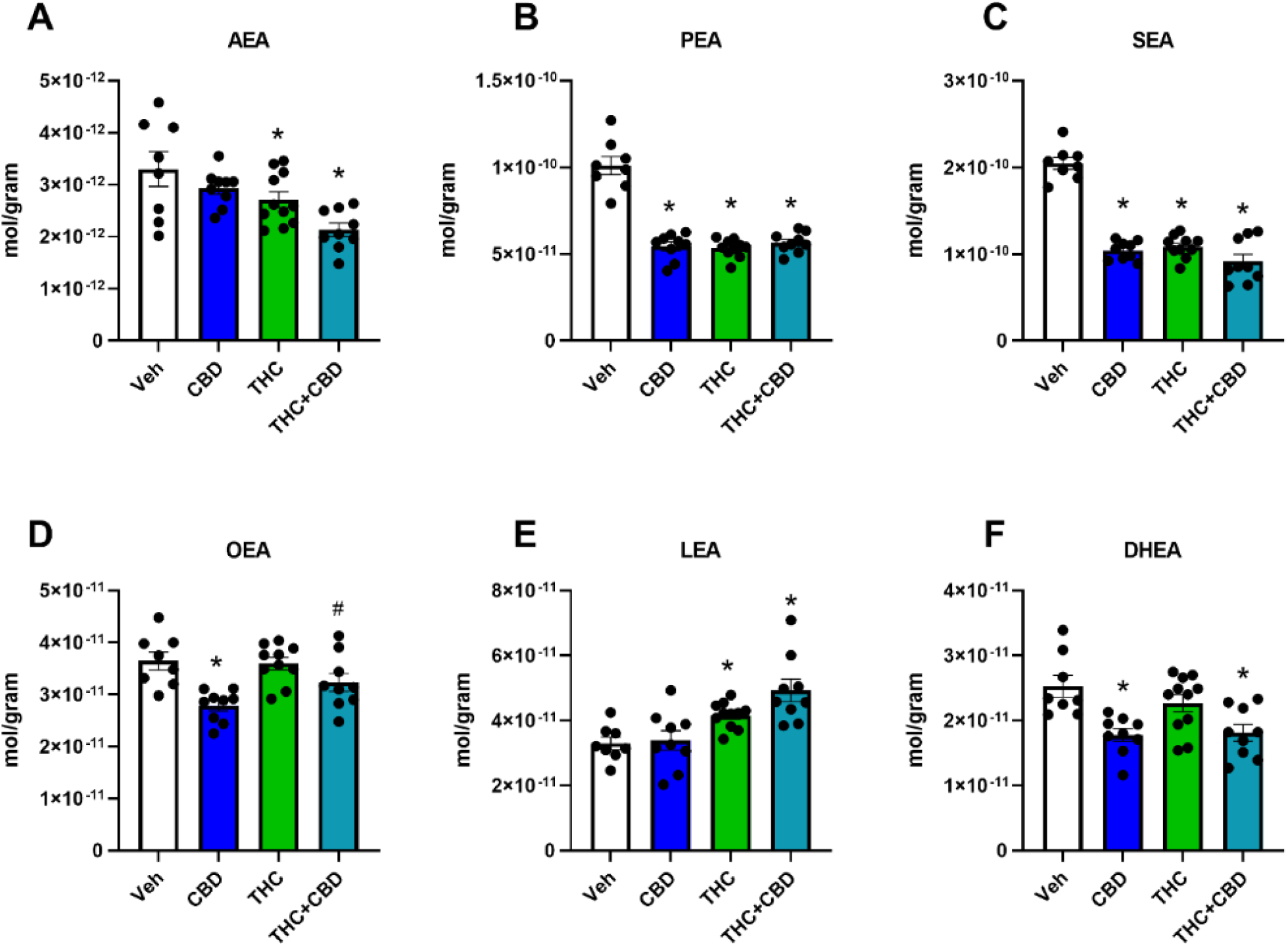
Levels of *N*-acyl ethanolamine in milk after CBD, THC, or THC+CBD (3mg/kg) or vehicle administration to lactating dams from P0-P10 collected from pup stomachs 2 hours after final injection to dams on P10. (A) Concentrations of *N*-arachidonoyl ethanolamine (AEA) in milk. Comparison of vehicle to CBD (*ns*), THC (p=0.035), or THC+CBD treatment(p=0.001). (B) Concentrations of *N*-palmitoyl ethanolamine (PEA) in milk. Comparison of vehicle to CBD (p=0.001), THC (p=0.001), or THC+CBD treatment(p=0.001). (C) Concentrations of *N*-stearoyl ethanolamine (SEA) in milk. Comparison of vehicle to CBD (p=0.001), THC (p=0.001), or THC+CBD treatment(p=0.001). (D) Concentrations of *N*-oleoyl ethanolamine (OEA) in milk. Comparison of vehicle to CBD (p=0.001), THC (*ns*), or THC+CBD treatment(p=0.054). (E) Concentrations of *N*-linoleoyl ethanolamine (LEA) in milk. Comparison of vehicle to CBD (*ns*), THC (p=0.018), or THC+CBD treatment(p=0.001). (F) Concentrations of *N*-docosahexaenoyl ethanolamine (DHEA) in milk. Comparison of vehicle to CBD (0.001), THC (*ns*), or THC+CBD treatment(p=0.001). Data represented as means ±SEM with individual data points from pups plotted as scatter. *ns*=no significance

## 4. Discussion

Here we characterized the lipid profile of milk taken from the stomach of pups nursed by dams that were chronically injected with either CBD, THC, or a 1:1 combination of THC+CBD when their pups were PND1-PND10. By comparing the lipidomic profiles across treatment groups, the data provide insight into how CBs are differentially present in breast milk and how chronic treatment with CBs modulates the lipid and lipid signaling molecule compositions.

### 4.1 Maternally administered cannabinoids are differentially present in breast milk

As found in human studies, CBD, THC, and THC metabolites are present in breast milk after maternal consumption [5-8]. Here, we add to this knowledge by showing that CBD metabolites are present in the milk as well. Across groups, concentrations of CBD in milk are significantly greater than concentrations of THC. This could be a result of slower CBD metabolism, greater transport/accumulation in milk, a combination of both, or some as-yet-unrecognized factor. Consistent with these findings, CBD was measured at a greater concentration and in a greater proportion of breast milk samples than plasma samples taken from women who reported using cannabis [8]. Our data also suggest that combining CBD with THC influences their rate of metabolism or accumulation in milk, in that we measured higher average concentrations of both CBs in the THC+CBD treatment group compared to either drug alone. This replicates previous findings that combination THC+CBD results in higher plasma levels of THC than when THC is administered alone [18]. Higher concentrations of 11-OH THC and 7-OH CBD in the combination group compared to either CB alone suggests that either the CBs are metabolized faster when combined, or that clearance of their metabolites occurs more slowly. Because the differences in concentrations can be caused by an increase in clearance or a decrease in accumulation, pharmacokinetic studies are needed to determine which is occurring.

### 4.2 Combination THC+CBD creates a third phenotype compared to changes in lipids seen in treatment with either CB alone

While CBD has been shown to prevent some changes in behavior seen after chronic THC exposure [18], the combination of CBD and THC in a 1:1 ratio does not result in an intermediate lipidomic profile in milk. Instead, the combination of both CBD and THC has a broad effect on lipids that is not always additive or predictable based on changes in response to one CB alone. 16% of the detected endogenous lipids in the THC+CBD group are altered in a way different from CBD alone or THC alone treatment. When changes occurred in the same direction as either drug alone, they tended to be in the same direction as changes seen in THC. This suggests that, despite having lower concentrations than CBD in milk, THC might have a stronger impact on the production of lipid signaling molecules in milk that is not modified by coadministration of CBD.

### 4.3 Milk free fatty acids are sensitive to CBs

During the limited period when milk must be produced and secreted, a host of morphological and transcriptional changes occur that result in increased transcription for proteins involved in fatty acid synthesis and decreased transcription for proteins involved in fatty acid degradation [19, 20]. Lipids are secreted from mammary tissue through cytoplasmic lipid droplets (CLD), and CLDs contain lipids either derived from maternal stores and circulation or synthesized *de novo* in mammary epithelial cells (reviewed; [21]). Long chain fatty acids secreted in milk are primarily derived from diet or other tissue stores, suggesting that these are the source of the FFAs measured here (reviewed; [22]). To confirm the origin of lipids secreted in milk, further investigation is needed to determine the mechanism through which systematic CB administration to a lactating dam decreases the free fatty acids AA, LA, OA, EPA, and DHA. In specific, analysis of dam plasma and liver tissue would provide insight into when circulating lipids are altered. The changes in FFAs in brain tissue and neural cell lines after CBD and THC treatment are distinct from the changes that we report in milk with acute CB exposure [15, 16], so it is possible that mammary epithelial cells respond uniquely to CBs and decrease synthesis of FFAs. However, this could also be an effect of chronic use, especially given that both CBD and THC had similar effects in overall suppression of FFAs. Practically, dietary supplementation with FFAs might not be able to restore lipid concentrations of all FFAs to baseline levels. In studies investigating the effects of dietary FFAs on milk content, DHA supplementation was able to increase DHA in deficient milk samples [23], but no correlation was found between dietary AA and levels in milk [11]. Additionally, there are likely many consequences to increasing dietary fatty acids during lactation, given that supplementation with DHA affects tissue and serum levels of amino acids in piglets, including increases in methionine in plasma, taurine and glycine in the liver, and decreases in methionine, glutamate, and leucine in the brain [24]. One caveat of the current study is that the milk is collected from the stomach of the pups rather than the mammary gland of the dam. This introduces the possibility that lipids in the milk had undergone gut processing by the time they were extracted and analyzed, which may be particularly relevant to FFA absorption. While early investigation of the fat content in stomach-derived milk shows that it contains a greater amount of fat than mammary-expressed milk [25], experimental manipulations that alter the fat content in expressed milk alter fat content from the stomach in a similar fashion [26, 27]. Therefore, the correlation between changes in breast milk and stomach milk supports our method to investigate stomach milk as an indicator of the lipid content of milk, though adding chronic CB exposure to these pups may change that dynamic.

### 4.5 *N*-acyl methionine species are decreased in milk after CB treatment

The decreases in *N*-acyl methionine species across all conditions are notable in part because of the relationship between methionine and fatty acids in milk. Methionine is an essential amino acid whose metabolic product is used in methyltransferase reactions (reviewed; [28]) that are essential for protein synthesis. Administering methionine to cow mammary epithelial cells leads to increases in triglyceride secretion [29], however, removing dietary methionine leads to significantly higher methionine levels in goat milk despite no changes in its mammary uptake [30]. Supplementation of DHA to developing pigs results in significantly higher methionine levels in plasma, brain tissue, and skeletal muscle; again, demonstrating an important link between levels of fatty acids in breast milk and methionine regulation [24]. Importantly, methionine also influences infant growth in that the addition of methionine to a protein-deficient diet during lactation increases weight gain in pups [31]. Therefore, decreases in *N-*acyl methionine species may simply indicate broad changes in milk composition; however, it is also possible that decreases in the FFA content of milk alter the subsequent availability of methionine for lipoamine biosynthesis. Taken together, the data on the importance of methionine during development are a valuable consideration when evaluating the dramatic changes in the methionine lipoamine species after chronic CB exposure.

### 4.8 Changes in lipids could impact feeding behavior and long-term development of infants

Breastmilk contains as-yet-unknown important bioactive compounds that are beneficial to infant development. This is evidenced by data showing that breastmilk-based diets have consistently better developmental outcomes for very underweight preterm infants than formula [32]. The benefits of breastmilk on infant growth and development are due in part to its lipid content. Adding AA to a dam’s diet during lactation increases litter birth weights, an effect that persists to PND28 [33]. Offspring are more sensitive to maternal dietary FA content than dams, and a diet high in linoleic acid results in litter weights that are significantly higher than those of a dam on a low-fat diet [34]. The dangers of a suboptimal fat content in milk are also well established. Administering a mixture of trans-10 and cis-10 conjugated linoleic acid (CLA), which decreases the fat content of breastmilk, leads to small litter weights. The effect can be rescued by discontinuing CLA treatment [26]. A maternal diet deficient in omega-3 fatty acids during pregnancy results in decreased levels of DHA and increased levels of AA in the brain, as well as decreased performance on hippocampal learning and memory tasks at 6 months that cannot be rescued by switching to an adequate diet at 2 or 4 months old [35]. Additionally, not all FAs appear to be equally beneficial. A diet containing fish oil, high in EPA and DHA, results in birthweights that are significantly smaller and pup sizes that are significantly shorter than those of a diet containing olive oil, rich in OA [36].

Outside of their presence in milk, a growing literature is showing that many of the lipids in our screening library are bioactive signaling molecules. *N*-linoleoyl tyrosine blunts behavioral deficits and neuronal damage caused by ischemic injury [37]; *N*-palmitoyl serine improves changes in neurological severity scores as animals recover from traumatic brain injury [38]; administering *N-*arachidonoyl or *N*-oleoyl taurine increases insulin secretion by rat islet cells, in part through TRPV1 channels [39]; and *N*-oleoyl taurine improves insulin secretion, in part through GPR119 activation [40]. The ability of these compounds to alter cell morphology, recovery outcomes after neurological injury, and bodily homeostasis suggest that changes in their availability to infants could have far-reaching effects. Here, we show that each of these lipids is affected by at least one of the CB treatment groups, which illustrates how unknown constituents of breast milk can have effects on typical development.

### 4.7 *N-*acyl ethanolamines predominately decrease after CB treatment

Mice administered either THC or CBD have distinct brain lipidomes, but both are marked by changes in NAEs. NAEs are largely decreased following THC treatment in the adult mouse brain [16] and increased after CBD treatment [15]. NAEs in milk are mostly decreased after CB treatments. The only increases are in *N*-linoleoyl ethanolamine in both the THC and THC+CBD condition, which follows the pattern of other lipoamines in the screen where linoleoyl conjugates increase after CB treatment. The maintenance of this pattern despite widespread decreases supports the importance of CB effects on *N-*acyl chains.

Maternal dietary intake of lipids are not always related to the endocannabinoid compounds measured in breastmilk [11]. However, NAEs in milk are correlated to infant growth outcomes. For instance, lower levels of OEA, SEA, and PEA are correlated to a higher weight-for-age z-score [12]. In animal models, early life repeated exposure to AEA (20μg/kg) leads to significantly more epididymal fat, larger adipocytes, and increased body weight that continued to be elevated once treatment stops [13]. Therefore, decreases in breastmilk NAEs caused by maternal CB consumption may translate into higher weight-for-age-z-score and impact the early life development of infants.

### 4.9 Many 2-sn-acyl glycerols are increased after CB treatment

Changes in 2-sn-acyl glycerol species, as opposed to NAEs, were all increases. CBD treatment alone resulted in particularly broad differences, with 2-PG, 2-OG, and 2-AG all significantly increasing. This appears to be similar to increases in 2-acyl-glycerol species seen after CBD treatment in the brains of wildtype mice [15]. Similar to NAEs, 2-sn-acyl glycerol exposure in early life can impact infant development. A cocktail of 2-AG, 2-LG, and 2-PG was capable of decreasing mortality rates caused by a reduction in feeding in mouse pups given an acute injection of CB1 antagonist SR141716A [3], suggesting a strong relationship to these lipids and normal feeding behavior.

## Conclusion

Lipidomic analysis of milk produced by lactating dams with chronic CBD, THC, or THC+CBD exposure reveals widespread changes bioactive lipids and CB presence in breastmilk. That the amount of CBD and THC in the milk are differentially regulated regardless of maternal exposure suggests metabolic and transport processes that are specific for each CB. All chronic CB treatments result in decreased levels of FFAs, which could be indicative of more extensive effects on milk composition than just changes in lipid signaling molecules. The marked differences in modulation of lipids dependent on fatty acid chain length and saturation also suggest dysregulation of finely tuned production of breastmilk lipids that likely affect more lipids derived from these FFAs than those measured here. The concept in both some aspects of the literature and widely accepted in culture that CBD “protects” against the effects of THC was not supported here in that the largest changes in lipid composition of breastmilk was in the THC+CBD combination group, though what constitutes “protection” is likely context dependent. Taken together, these data provide compelling evidence that maternal exposure to CBs during lactation have significant outcomes on breastmilk composition and by extension infant health.

## References

1. Barbosa-Leiker, C., et al., Daily Cannabis Use During Pregnancy and Postpartum in a State With Legalized Recreational Cannabis. J Addict Med, 2020. 14(6): p. 467–474.

2. Philippot, G., et al., Short-term exposure and long-term consequences of neonatal exposure to Δ(9)-tetrahydrocannabinol (THC) and ibuprofen in mice. Behav Brain Res, 2016. 307: p. 137–44.

3. Fride, E., et al., Critical role of the endogenous cannabinoid system in mouse pup suckling and growth. Eur J Pharmacol, 2001. 419(2-3): p. 207–14.

4. Maciel, I.S., et al., Perinatal CBD or THC Exposure Results in Lasting Resistance to Fluoxetine in the Forced Swim Test: Reversal by Fatty Acid Amide Hydrolase Inhibition. Cannabis Cannabinoid Res, 2021.

5. Wymore, E.M., et al., Persistence of Δ-9-Tetrahydrocannabinol in Human Breast Milk. JAMA Pediatrics, 2021.

6. Baker, T., et al., Transfer of Inhaled Cannabis Into Human Breast Milk. Obstet Gynecol, 2018. 131(5): p. 783–788.

7. Bertrand, K.A., et al., Marijuana Use by Breastfeeding Mothers and Cannabinoid Concentrations in Breast Milk. Pediatrics, 2018. 142(3).

8. Moss, M.J., et al., Cannabis use and measurement of cannabinoids in plasma and breast milk of breastfeeding mothers. Pediatr Res, 2021.

9. Gaitán, A.V., et al., Endocannabinoid Metabolome Characterization of Milk from Guatemalan Women Living in the Western Highlands. Curr Dev Nutr, 2019. 3(6): p. nzz018.

10. Gaitán, A.V., et al., Endocannabinoid Metabolome Characterization of Transitional and Mature Human Milk. Nutrients, 2018. 10(9).

11. Gaitán, A.V., et al., Maternal Dietary Fatty Acids and Their Relationship to Derived Endocannabinoids in Human Milk. J Hum Lact, 2021: p. 890334421993468.

12. Bruun, S., et al., Satiety Factors Oleoylethanolamide, Stearoylethanolamide, and Palmitoylethanolamide in Mother’s Milk Are Strongly Associated with Infant Weight at Four Months of Age-Data from the Odense Child Cohort. Nutrients, 2018. 10(11).

13. Aguirre, C.A., V.A. Castillo, and M.N. Llanos, The endocannabinoid anandamide during lactation increases body fat content and CB1 receptor levels in mice adipose tissue. Nutr Diabetes, 2015. 5(6): p. e167.

14. Aguirre, C., V. Castillo, and M. Llanos, Oral Administration of the Endocannabinoid Anandamide during Lactation: Effects on Hypothalamic Cannabinoid Type 1 Receptor and Food Intake in Adult Mice. J Nutr Metab, 2017. 2017: p. 2945010.

15. Leishman, E., et al., Cannabidiol’s Upregulation of N-acyl Ethanolamines in the Central Nervous System Requires N-acyl Phosphatidyl Ethanolamine-Specific Phospholipase D. Cannabis Cannabinoid Res, 2018. 3(1): p. 228–241.

16. Leishman, E., et al., Δ(9)-Tetrahydrocannabinol changes the brain lipidome and transcriptome differentially in the adolescent and the adult. Biochim Biophys Acta Mol Cell Biol Lipids, 2018. 1863(5): p. 479–492.

17. Leishman, E., et al., Lipidomics profile of a NAPE-PLD KO mouse provides evidence of a broader role of this enzyme in lipid metabolism in the brain. Biochim Biophys Acta, 2016. 1861(6): p. 491–500.

18. Murphy, M., et al., Chronic Adolescent Δ(9)-Tetrahydrocannabinol Treatment of Male Mice Leads to Long-Term Cognitive and Behavioral Dysfunction, Which Are Prevented by Concurrent Cannabidiol Treatment. Cannabis Cannabinoid Res, 2017. 2(1): p. 235–246.

19. Rudolph, M.C., et al., Functional development of the mammary gland: use of expression profiling and trajectory clustering to reveal changes in gene expression during pregnancy, lactation, and involution. J Mammary Gland Biol Neoplasia, 2003. 8(3): p. 287–307.

20. Gopalakrishnan, K., et al., Histology and Transcriptome Profiles of the Mammary Gland across Critical Windows of Development in Sprague Dawley Rats. J Mammary Gland Biol Neoplasia, 2018. 23(3): p. 149–163.

21. Rudolph, M.C., M.C. Neville, and S.M. Anderson, Lipid synthesis in lactation: diet and the fatty acid switch. J Mammary Gland Biol Neoplasia, 2007. 12(4): p. 269–81.

22. Neville, M.C. and M.F. Picciano, REGULATION OF MILK LIPID SECRETION AND COMPOSITION. Annual Review of Nutrition, 1997. 17(1): p. 159–184.

23. Jackson, K.H., et al., Baseline red blood cell and breast milk DHA levels affect responses to standard dose of DHA in lactating women on a controlled feeding diet. Prostaglandins, Leukotrienes and Essential Fatty Acids, 2021. 166.

24. Li, P., et al., Dietary supplementation with cholesterol and docosahexaenoic acid affects concentrations of amino acids in tissues of young pigs. Amino Acids, 2009. 37(4): p. 709–16.

25. Naismith, D.J., A. Mittwoch, and B.S. Platt, Changes in composition of rat’s milk in the stomach of the suckling. Br J Nutr, 1969. 23(3): p. 683–93.

26. Harvatine, K.J., et al., Trans-10, cis-12 CLA dose-dependently inhibits milk fat synthesis without disruption of lactation in C57BL/6J mice. J Nutr, 2014. 144(12): p. 1928–34.

27. Richard, C., et al., The content of docosahexaenoic acid in the maternal diet differentially affects the immune response in lactating dams and suckled offspring. Eur J Nutr, 2016. 55(7): p. 2255–64.

28. Brosnan, J.T. and M.E. Brosnan, The sulfur-containing amino acids: an overview. J Nutr, 2006. 136(6 Suppl): p. 1636s–1640s.

29. Li, P., et al., CRTC2 Is a Key Mediator of Amino Acid-Induced Milk Fat Synthesis in Mammary Epithelial Cells. J Agric Food Chem, 2019. 67(37): p. 10513–10520.

30. Liu, W., et al., Short-term lactation and mammary metabolism responses in lactating goats to graded removal of methionine from an intravenously infused complete amino acid mixture. J Dairy Sci, 2019. 102(5): p. 4094–4104.

31. Liu, G.M., et al., Methionine, leucine, isoleucine, or threonine effects on mammary cell signaling and pup growth in lactating mice. J Dairy Sci, 2017. 100(5): p. 4038–4050.

32. Ottolini, K.M., et al., Improved brain growth and microstructural development in breast milk-fed very low birth weight premature infants. Acta Paediatr, 2020. 109(8): p. 1580–1587.

33. Hadley, K.B., et al., Supplementing dams with both arachidonic and docosahexaenoic acid has beneficial effects on growth and immune development. Prostaglandins Leukot Essent Fatty Acids, 2017. 126: p. 55–63.

34. Walker, R.E., et al., Dietary SFAs and ω-6 Fatty Acids Alter Incorporation of ω-3 Fatty Acids into Milk Fat of Lactating CD-1 Mice and Tissues of Offspring. J Nutr, 2021.

35. Lozada, L.E., et al., Perinatal Brain Docosahexaenoic Acid Concentration Has a Lasting Impact on Cognition in Mice. J Nutr, 2017. 147(9): p. 1624–1630.

36. Jiménez, M.J., et al., Fish oil diet in pregnancy and lactation reduces pup weight and modifies newborn hepatic metabolic adaptations in rats. Eur J Nutr, 2017. 56(1): p. 409–420.

37. Cheng, L., et al., N-Linoleyltyrosine Protects against Transient Cerebral Ischemia in Gerbil via CB2 Receptor Involvement in PI3K/Akt Signaling Pathway. Biol Pharm Bull, 2019. 42(11): p. 1867–1876.

38. Mann, A., et al., Palmitoyl Serine: An Endogenous Neuroprotective Endocannabinoid-Like Entity After Traumatic Brain Injury. J Neuroimmune Pharmacol, 2015. 10(2): p. 356–63.

39. Waluk, D.P., et al., N-Acyl taurines trigger insulin secretion by increasing calcium flux in pancreatic β-cells. Biochem Biophys Res Commun, 2013. 430(1): p. 54–9.

40. Grevengoed, T.J., et al., <em>N</em>-acyl taurines are endogenous lipid messengers that improve glucose homeostasis. Proceedings of the National Academy of Sciences, 2019. 116(49): p. 24770–24778.

