## Supplemental Figures for "Cannabinoids accumulate in mouse breast milk and differentially regulate lipid composition and lipid signaling molecules involved in infant development"

Supplemental Figure 1. Lipids in HPLC/MS/MS screening library

Supplemental Figure 2. Lipid levels (moles/gram) in milk collected form the stomachs of pups nursing a dam injected from postnatal day (PND) 0 to PND 10 with vehicle, CBD (3mg/kg), THC (3mg/kg), or THC+CBD (3mg/kg) and descriptive statistics.

Supplemental Figure 3. ANOVA for the effect of CBD, THC, or THC+CBD injections to a dam on *N-*acyl alanines measured in milk collected from nursing pups’ stomachs

Supplemental Figure 4. Post-hoc Fisher’s LSD results for *N-*acyl alanines

Supplemental Figure 5. ANOVA for the effect of CBD, THC, or THC+CBD injections to a dam on *N-*acyl ethanolamines measured in milk collected from nursing pups’ stomachs

Supplemental Figure 6. Post-hoc Fisher’s LSD results for *N-*acyl ethanolamines

Supplemental Figure 7. ANOVA for the effect of CBD, THC, or THC+CBD injections to a dam on *N-*acyl GABAs measured in milk collected from nursing pups’ stomachs

Supplemental Figure 8. Post-hoc Fisher’s LSD results for *N-*acyl GABAs

Supplemental Figure 9. ANOVA for the effect of CBD, THC, or THC+CBD injections to a dam on *N-*acyl glycines measured in milk collected from nursing pups’ stomachs

Supplemental Figure 10. Post-hoc Fisher’s LSD results for *N-*acyl glycines

Supplemental Figure 11. ANOVA for the effect of CBD, THC, or THC+CBD injections to a dam on *N-*acyl leucines measured in milk collected from nursing pups’ stomachs

Supplemental Figure 12. Post-hoc Fisher’s LSD results for *N-*acyl leucines

Supplemental Figure 13. ANOVA for the effect of CBD, THC, or THC+CBD injections to a dam on *N-*acyl methionines measured in milk collected from nursing pups’ stomachs

Supplemental Figure 14. Post-hoc Fisher’s LSD results for *N-*acyl methionines

Supplemental Figure 15. ANOVA for the effect of CBD, THC, or THC+CBD injections to a dam on *N-*acyl phenylalanines measured in milk collected from nursing pups’ stomachs

Supplemental Figure 16. Post-hoc Fisher’s LSD results for *N-*acyl phenylalanines

Supplemental Figure 17. ANOVA for the effect of CBD, THC, or THC+CBD injections to a dam on *N-*acyl prolines measured in milk collected from nursing pups’ stomachs

Supplemental Figure 18. Post-hoc Fisher’s LSD results for *N-*acyl prolines

Supplemental Figure 19. ANOVA for the effect of CBD, THC, or THC+CBD injections to a dam on *N-*acyl serines measured in milk collected from nursing pups’ stomachs

Supplemental Figure 20. Post-hoc Fisher’s LSD results for *N-*acyl serines

Supplemental Figure 21. ANOVA for the effect of CBD, THC, or THC+CBD injections to a dam on *N-*acyl taurines measured in milk collected from nursing pups’ stomachs

Supplemental Figure 22. Post-hoc Fisher’s LSD results for *N-*acyl taurines

Supplemental Figure 23. ANOVA for the effect of CBD, THC, or THC+CBD injections to a dam on *N-*acyl tryptophans measured in milk collected from nursing pups’ stomachs

Supplemental Figure 24. Post-hoc Fisher’s LSD results for *N-*acyl tryptophans

Supplemental Figure 25. ANOVA for the effect of CBD, THC, or THC+CBD injections to a dam on *N-*acyl tyrosines measured in milk collected from nursing pups’ stomachs

Supplemental Figure 26. Post-hoc Fisher’s LSD results for *N-*acyl tyrosines

Supplemental Figure 27. ANOVA for the effect of CBD, THC, or THC+CBD injections to a dam N-acyl valines measured in milk collected from nursing pups’ stomachs

Supplemental Figure 28. Post-hoc Fisher’s LSD results for *N-*acyl valines

Supplemental Figure 29. ANOVA for the effect of CBD, THC, or THC+CBD injections to a dam on 2-acyl-glycerols measured in milk collected from nursing pups’ stomachs

Supplemental Figure 30. Post-hoc Fisher’s LSD results for 2-acyl-glycerols

Supplemental 31. ANOVA for the effect of CBD, THC, or THC+CBD injections to a dam on free fatty acids measured in milk collected from nursing pups’ stomachs

Supplemental 32. Post-hoc Fisher’s LSD results for free fatty acids

Supplemental Figure 33. Descriptive statistics for CBD and THC measured in milk taken from stomachs of pups nursing dams injected with 3mg/kg CBD, THC, or THC+CBD

Supplemental Figure 34. ANOVA for the effect of CBD, THC, or THC+CBD injections administered to a dam on cannabinoids measured in milk collected from nursing pups’ stomachs

Supplemental Figure 35. Post-hoc Fisher’s LSD results for cannabinoids

Supplemental Figure 36: Descriptive statistics for 11-OH-THC and 7-OH-CBD measured in milk from the stomachs of pups nursing a dam injected with (3mg/kg) CBD, THC, or THC+CBD

Supplemental Figure 37: ANOVA for the effect of CBD, THC, or THC+CBD injections to a dam on 11-OH-THC measured in milk collected from nursing pups’ stomachs

Supplemental Figure 38: ANOVA for the effect of CBD, THC, or THC+CBD injections to a nursing dam on 7-OH-CBD measured in milk collected from pups’ stomachs

Supplemental Figure 39: Example chromatograms identifying 7-OH-CBD in standard and milk samples from each treatment group

### Supplemental Figure 1. Lipids in HPLC/MS/MS screening library

| *N-*acyl alanine | [M–H]^-^ | Fragment |  | *N-*acyl proline | [M–H]^-^ | Fragment |
| --- | --- | --- | --- | --- | --- | --- |
| *N-*palmitoyl alanine | 326.5 | 88.09 |  | *N-*palmitoyl proline | 352.53 | 114.12 |
| *N-*stearoyl alanine | 354.55 | 88.09 |  | *N-*stearoyl proline | 380.59 | 114.12 |
| *N-*oleoyl alanine | 352.53 | 88.09 |  | *N-*oleoyl proline | 378.31 | 114.12 |
| *N-*linoleoyl alanine | 350.52 | 88.09 |  | *N-*linoleoyl proline | 376.56 | 114.12 |
| *N-*arachidonoyl alanine | 374.5 | 88.09 |  | *N-*arachidonoyl proline | 400.58 | 114.12 |
| *N-*docosahexaenoyl alanine | 398.56 | 88.09 |  | *N-*docosahexaenoyl proline | 424.6 | 114.12 |
| *N-*acyl ethanolamine | [M+H]^+^ | Fragment |  | *N-*acyl serine | [M–H]^-^ | Fragment |
| *N-*palmitoyl ethanolamine | 300.29 | 62.1 |  | *N-*palmitoyl serine | 342.3 | 74 |
| *N-*stearoyl ethanolamine | 328.3 | 62.1 |  | *N-*stearoyl serine | 370.3 | 74 |
| *N-*oleoyl ethanolamine | 326.3 | 62.1 |  | *N-*oleoyl serine | 368.3 | 74 |
| *N-*linoleoyl ethanolamine | 324.3 | 62.1 |  | *N-*linoleoyl serine | 366.27 | 74 |
| *N-*arachidonoyl ethanolamine | 348.29 | 62.1 |  | *N-*arachidonoyl serine | 390.3 | 74 |
| *N-*docosahexaenoyl ethanolamine | 372.6 | 62.1 |  | *N-*docosahexaenoyl serine | 414.3 | 74 |
| *N-*acyl GABA | [M–H]^-^ | Fragment |  | *N-*acyl taurine | [M–H]^-^ | Fragment |
| *N-*palmitoyl GABA | 340.54 | 102.1 |  | *N-*palmitoyl taurine | 362.6 | 124 |
| *N-*stearoyl GABA | 368.58 | 102.1 |  | *N-*stearoyl taurine | 390.6 | 124 |
| *N-*oleoyl GABA | 366.57 | 102.1 |  | *N-*oleoyl taurine | 388.6 | 124 |
| *N-*linoleoyl GABA | 364.54 | 102.1 |  | *N-*arachidonoyl taurine | 410.6 | 124 |
| *N-*arachidonoyl GABA | 388.57 | 102.1 |  | *N-*acyl tryptophan | [M–H]^-^ | Fragment |
| *N-*docosahexaenoyl GABA | 412.59 | 102.1 |  | *N-*palmitoyl tryptophan | 441.63 | 203.1 |
| *N-*acyl glycine | [M–H]^-^ | Fragment |  | *N-*stearoyl tryptophan | 469.68 | 203.1 |
| *N-*palmitoyl glycine | 312.26 | 74.2 |  | *N-*oleoyl tryptophan | 467.67 | 203.1 |
| *N-*stearoyl glycine | 340.3 | 74.2 |  | *N-*linoleoyl tryptophan | 465.65 | 203.1 |
| *N-*oleoyl glycine | 338.3 | 74.2 |  | *N-*arachidonoyl tryptophan | 489.67 | 203.1 |
| *N-*linoleoyl glycine | 336.3 | 74.2 |  | *N-*docosahexaenoyl tryptophan | 513.69 | 203.1 |
| *N-*arachidonoyl glycine | 360.3 | 74.2 |  | *N-*acyl tyrosine | [M–H]^-^ | Fragment |
| *N-*docosahexaenoyl glycine | 384.3 | 74.2 |  | *N-*palmitoyl tyrosine | 418.59 | 180.18 |
| *N-*acyl leucine | [M–H]^-^ | Fragment |  | *N-*stearoyl tyrosine | 446.65 | 180.18 |
| *N-*palmitoyl leucine | 368.58 | 130.1 |  | *N-*oleoyl tyrosine | 444.63 | 180.18 |
| *N-*stearoyl leucine | 396.63 | 130.1 |  | *N-*linoleoyl tyrosine | 442.61 | 180.18 |
| *N-*oleoyl leucine | 394.61 | 130.1 |  | *N-*arachidonoyl tyrosine | 466 | 180.18 |
| *N-*linoleoyl leucine | 392.6 | 130.1 |  | *N-*docosahexaenoyl tyrosine | 490.66 | 180.18 |
| *N-*docosahexaenoyl leucine | 440.64 | 130.1 |  | *N-*acyl valine | [M–H]^-^ | Fragment |
| *N-*acyl methionine | [M–H]^-^ | Fragment |  | *N-*palmitoyl valine | 354.31 | 116.31 |
| *N-*palmitoyl methionine | 386.62 | 148.2 |  | *N-*stearoyl valine | 382.6 | 116.14 |
| *N-*stearoyl methionine | 414.64 | 148.2 |  | *N-*oleoyl valine | 380.59 | 116.14 |
| *N-*oleoyl methionine | 412.65 | 148.2 |  | *N-*linoleoyl valine | 378.58 | 116.14 |
| *N-*linoleoyl methionine | 410.64 | 148.2 |  | *N-*docosahexaenoyl valine | 426.62 | 116.14 |
| *N-*arachidonoyl methionine | 434.66 | 148.2 |  | Free Fatty Acids | [M–H]^-^ | Fragment |
| *N-*docosahexaenoyl methionine | 458.68 | 148.2 |  | Oleic acid | 281.5 | 263 |
| *N-*acyl phenylalanine | [M–H]^-^ | Fragment |  | Linoleic acid | 279.5 | 261 |
| *N-*palmitoyl phenylalanine | 402.59 | 164.1 |  | Arachidonic acid | 303.5 | 285 |
| *N-*stearoyl phenylalanine | 430.65 | 164.1 |  | Docosahexaenoic acid | 327.5 | 283 |
| *N-*oleoyl phenylalanine | 428.63 | 164.1 |  | Eicosapentaenoic acid | 301.5 | 257 |
| *N-*linoleoyl phenylalanine | 426.61 | 164.1 |  | 2-acyl glycerol | [M+H]^+^ | Fragment |
| *N-*arachidonoyl phenylalanine | 450.64 | 164.1 |  | 2-arachidonoyl glycerol | 379.3 | 287.5 |
| *N-*docosahexaenoyl phenylalanine | 474.66 | 164.1 |  | 2-linoleoyl glycerol | 355.5 | 245 |
| Phytocannabinoids | [M+H]^+^ | Fragment |  | 2-oleoyl glycerol | 357.5 | 265.2 |
| CBD | 315.2 | 191 |  | 2-palmitoyl glycerol | 331.5 | 239.5 |
| THC | 315.2 | 123.2 |  | Prostaglandins | [M–H]^-^ | Fragment |
| CB Metabolites | [M–H]^-^ | Fragment |  | PGE_2_ | 351.2 | 315 |
| 7-OH-CBD | 329.5 | 173 |  | PGF_2α_ | 353.3 | 309.2 |
| 11-OH-THC | 329.5 | 173 |  | 6-ketoPGF_1α_ | 369.3 | 206.9 |

### Supplemental Figure 2. Lipid levels (moles/gram) in milk collected from the stomachs of pups nursing a dam injected from postnatal day (PND) 0 to PND 10 with vehicle, CBD (3mg/kg), THC (3mg/kg), or THC+CBD (3mg/kg) and descriptive statistics.

Compounds present in too few samples for statistical analyses are denoted by “BAL” or below analytical levels. Compounds detected in no samples are denoted by “BDL” or below detectable levels.

|  |  | N | Mean | Std. Deviation | Std. Error | 95% Confidence Interval for Mean | | Minimum | Maximum |
| --- | --- | --- | --- | --- | --- | --- | --- | --- | --- |
|  |  |  |  |  |  | Lower Bound | Upper Bound |  |  |
| *N-*acyl alanines |  |  |  |  |  |  |  |  |  |
| *N-*palmitoyl alanine | Vehicle | 8 | 4.77E-12 | 1.61E-12 | 5.70E-13 | 3.42E-12 | 6.12E-12 | 2.42E-12 | 7.84E-12 |
|  | CBD | 9 | 4.99E-12 | 1.76E-12 | 5.88E-13 | 3.63E-12 | 6.34E-12 | 2.29E-12 | 7.73E-12 |
|  | THC | 11 | 4.52E-12 | 1.76E-12 | 5.32E-13 | 3.34E-12 | 5.70E-12 | 2.59E-12 | 7.78E-12 |
|  | THC+CBD | 8 | 4.84E-12 | 1.55E-12 | 5.48E-13 | 3.55E-12 | 6.14E-12 | 3.47E-12 | 7.96E-12 |
| *N-*stearoyl alanine | Vehicle | 8 | 2.85E-12 | 1.65E-12 | 5.83E-13 | 1.48E-12 | 4.23E-12 | 1.18E-12 | 6.03E-12 |
|  | CBD | 8 | 2.05E-12 | 7.42E-13 | 2.62E-13 | 1.43E-12 | 2.67E-12 | 1.29E-12 | 3.55E-12 |
|  | THC | 10 | 1.65E-12 | 3.54E-13 | 1.12E-13 | 1.40E-12 | 1.90E-12 | 1.05E-12 | 2.11E-12 |
|  | THC+CBD | 8 | 1.83E-12 | 3.96E-13 | 1.40E-13 | 1.50E-12 | 2.17E-12 | 1.32E-12 | 2.29E-12 |
| *N-*oleoyl alanine | Vehicle | 7 | 1.10E-12 | 4.35E-13 | 1.64E-13 | 7.02E-13 | 1.51E-12 | 5.29E-13 | 1.89E-12 |
|  | CBD | 9 | 1.21E-12 | 8.60E-13 | 2.87E-13 | 5.48E-13 | 1.87E-12 | 5.07E-13 | 2.83E-12 |
|  | THC | 11 | 9.45E-13 | 2.80E-13 | 8.45E-14 | 7.56E-13 | 1.13E-12 | 5.40E-13 | 1.35E-12 |
|  | THC+CBD | 8 | 1.09E-12 | 1.27E-13 | 4.48E-14 | 9.81E-13 | 1.19E-12 | 9.37E-13 | 1.24E-12 |
| *N-*linoleoyl alanine | Vehicle | 8 | 1.63E-12 | 6.49E-13 | 2.30E-13 | 1.09E-12 | 2.17E-12 | 6.39E-13 | 2.60E-12 |
|  | CBD | 8 | 2.89E-12 | 8.48E-13 | 3.00E-13 | 2.18E-12 | 3.60E-12 | 1.32E-12 | 3.93E-12 |
|  | THC | 10 | 3.76E-12 | 3.00E-13 | 9.47E-14 | 3.55E-12 | 3.97E-12 | 3.21E-12 | 4.16E-12 |
|  | THC+CBD | 9 | 4.47E-12 | 9.41E-13 | 3.14E-13 | 3.75E-12 | 5.19E-12 | 3.24E-12 | 5.77E-12 |
| *N-*arachidonoyl alanine | Vehicle | BDL |  |  |  |  |  |  |  |
|  | CBD | BDL |  |  |  |  |  |  |  |
|  | THC | BDL |  |  |  |  |  |  |  |
|  | THC+CBD | BDL |  |  |  |  |  |  |  |
| *N-*docosahexaenoyl alanine | Vehicle | BDL |  |  |  |  |  |  |  |
|  | CBD | BDL |  |  |  |  |  |  |  |
|  | THC | BDL |  |  |  |  |  |  |  |
|  | THC+CBD | BDL |  |  |  |  |  |  |  |
| *N-*acyl ethanolamines |  |  |  |  |  |  |  |  |  |
| *N-*palmitoyl ethanolamine | Vehicle | 8 | 1.01E-10 | 1.47E-11 | 5.21E-12 | 8.87E-11 | 1.13E-10 | 7.92E-11 | 1.27E-10 |
|  | CBD | 9 | 5.44E-11 | 7.53E-12 | 2.51E-12 | 4.86E-11 | 6.02E-11 | 4.03E-11 | 6.26E-11 |
|  | THC | 10 | 5.33E-11 | 5.23E-12 | 1.66E-12 | 4.96E-11 | 5.70E-11 | 4.21E-11 | 5.97E-11 |
|  | THC+CBD | 9 | 5.66E-11 | 5.73E-12 | 1.91E-12 | 5.22E-11 | 6.10E-11 | 4.69E-11 | 6.48E-11 |
| *N-*stearoyl ethanolamine | Vehicle | 8 | 2.05E-10 | 1.92E-11 | 6.80E-12 | 1.89E-10 | 2.21E-10 | 1.77E-10 | 2.41E-10 |
|  | CBD | 9 | 1.04E-10 | 1.06E-11 | 3.52E-12 | 9.59E-11 | 1.12E-10 | 8.89E-11 | 1.18E-10 |
|  | THC | 10 | 1.09E-10 | 1.30E-11 | 4.11E-12 | 9.94E-11 | 1.18E-10 | 8.35E-11 | 1.27E-10 |
|  | THC+CBD | 9 | 9.13E-11 | 2.49E-11 | 8.29E-12 | 7.22E-11 | 1.10E-10 | 6.31E-11 | 1.26E-10 |
| *N-*oleoyl ethanolamine | Vehicle | 8 | 3.65E-11 | 4.94E-12 | 1.75E-12 | 3.24E-11 | 4.06E-11 | 2.98E-11 | 4.48E-11 |
|  | CBD | 9 | 2.79E-11 | 3.04E-12 | 1.01E-12 | 2.55E-11 | 3.02E-11 | 2.25E-11 | 3.12E-11 |
|  | THC | 10 | 3.60E-11 | 3.74E-12 | 1.18E-12 | 3.33E-11 | 3.86E-11 | 2.92E-11 | 4.04E-11 |
|  | THC+CBD | 9 | 3.23E-11 | 5.22E-12 | 1.74E-12 | 2.83E-11 | 3.63E-11 | 2.48E-11 | 4.13E-11 |
| *N-*linoleoyl ethanolamine | Vehicle | 8 | 3.30E-11 | 5.41E-12 | 1.91E-12 | 2.84E-11 | 3.75E-11 | 2.46E-11 | 4.25E-11 |
|  | CBD | 9 | 3.39E-11 | 8.95E-12 | 2.98E-12 | 2.70E-11 | 4.08E-11 | 2.03E-11 | 4.93E-11 |
|  | THC | 11 | 4.16E-11 | 3.92E-12 | 1.18E-12 | 3.90E-11 | 4.43E-11 | 3.43E-11 | 4.79E-11 |
|  | THC+CBD | 9 | 4.93E-11 | 1.04E-11 | 3.48E-12 | 4.13E-11 | 5.73E-11 | 3.84E-11 | 7.09E-11 |
| *N-*arachidonoyl ethanolamine | Vehicle | 8 | 3.31E-12 | 9.49E-13 | 3.36E-13 | 2.51E-12 | 4.10E-12 | 2.02E-12 | 4.58E-12 |
|  | CBD | 9 | 2.94E-12 | 3.51E-13 | 1.17E-13 | 2.67E-12 | 3.21E-12 | 2.36E-12 | 3.55E-12 |
|  | THC | 11 | 2.72E-12 | 4.95E-13 | 1.49E-13 | 2.38E-12 | 3.05E-12 | 2.12E-12 | 3.46E-12 |
|  | THC+CBD | 9 | 2.13E-12 | 3.85E-13 | 1.28E-13 | 1.84E-12 | 2.43E-12 | 1.48E-12 | 2.64E-12 |
| *N-*docosahexaenoyl ethanolamine | Vehicle | 8 | 2.53E-11 | 4.81E-12 | 1.70E-12 | 2.13E-11 | 2.93E-11 | 2.09E-11 | 3.39E-11 |
|  | CBD | 9 | 1.78E-11 | 2.95E-12 | 9.82E-13 | 1.55E-11 | 2.01E-11 | 1.16E-11 | 2.13E-11 |
|  | THC | 11 | 2.27E-11 | 4.33E-12 | 1.31E-12 | 1.98E-11 | 2.56E-11 | 1.54E-11 | 2.75E-11 |
|  | THC+CBD | 9 | 1.81E-11 | 3.89E-12 | 1.30E-12 | 1.51E-11 | 2.11E-11 | 1.27E-11 | 2.33E-11 |
| *N-*acyl GABAs |  |  |  |  |  |  |  |  |  |
| *N-*palmitoyl GABA | Vehicle | 3 | 1.13E-12 | 4.47E-13 | 2.58E-13 | 1.79E-14 | 2.24E-12 | 8.30E-13 | 1.64E-12 |
|  | CBD | 7 | 9.85E-13 | 3.60E-13 | 1.36E-13 | 6.53E-13 | 1.32E-12 | 6.05E-13 | 1.54E-12 |
|  | THC | 6 | 9.28E-13 | 9.78E-14 | 3.99E-14 | 8.26E-13 | 1.03E-12 | 7.76E-13 | 1.07E-12 |
|  | THC+CBD | 8 | 1.16E-12 | 2.56E-13 | 9.06E-14 | 9.46E-13 | 1.37E-12 | 8.66E-13 | 1.50E-12 |
| *N-*stearoyl GABA | Vehicle | BDL |  |  |  |  |  |  |  |
|  | CBD | BDL |  |  |  |  |  |  |  |
|  | THC | BDL |  |  |  |  |  |  |  |
|  | THC+CBD | BDL |  |  |  |  |  |  |  |
| *N-*oleoyl GABA | Vehicle | BDL |  |  |  |  |  |  |  |
|  | CBD | BDL |  |  |  |  |  |  |  |
|  | THC | BDL |  |  |  |  |  |  |  |
|  | THC+CBD | BDL |  |  |  |  |  |  |  |
| *N-*linoleoyl GABA | Vehicle | BAL |  |  |  |  |  |  |  |
|  | CBD | BAL |  |  |  |  |  |  |  |
|  | THC | 7 | 7.25E-13 | 1.04E-13 | 3.92E-14 | 6.29E-13 | 8.21E-13 | 5.76E-13 | 8.43E-13 |
|  | THC+CBD | 8 | 8.55E-13 | 2.85E-13 | 1.01E-13 | 6.17E-13 | 1.09E-12 | 5.22E-13 | 1.21E-12 |
| *N-*arachidonoyl GABA | Vehicle | BDL |  |  |  |  |  |  |  |
|  | CBD | BDL |  |  |  |  |  |  |  |
|  | THC | BDL |  |  |  |  |  |  |  |
|  | THC+CBD | BDL |  |  |  |  |  |  |  |
| *N-*docosahexaenoyl GABA | Vehicle | BDL |  |  |  |  |  |  |  |
|  | CBD | BAL |  |  |  |  |  |  |  |
|  | THC | 9 | 2.18E-13 | 6.36E-14 | 2.12E-14 | 1.69E-13 | 2.67E-13 | 1.08E-13 | 3.37E-13 |
|  | THC+CBD | 4 | 1.32E-13 | 4.65E-14 | 2.33E-14 | 5.84E-14 | 2.06E-13 | 9.68E-14 | 1.95E-13 |
| *N-*acyl glycines |  |  |  |  |  |  |  |  |  |
| *N-*palmitoyl glycine | Vehicle | 8 | 5.56E-12 | 1.72E-12 | 6.07E-13 | 4.12E-12 | 7.00E-12 | 3.92E-12 | 8.93E-12 |
|  | CBD | 8 | 5.45E-12 | 1.03E-12 | 3.64E-13 | 4.59E-12 | 6.31E-12 | 3.52E-12 | 6.95E-12 |
|  | THC | 10 | 6.62E-12 | 1.32E-12 | 4.17E-13 | 5.68E-12 | 7.56E-12 | 5.36E-12 | 9.16E-12 |
|  | THC+CBD | 9 | 6.95E-12 | 1.50E-12 | 5.00E-13 | 5.80E-12 | 8.10E-12 | 4.86E-12 | 9.30E-12 |
| *N-*stearoyl glycine | Vehicle | 7 | 8.68E-13 | 4.78E-13 | 1.81E-13 | 4.27E-13 | 1.31E-12 | 3.42E-13 | 1.69E-12 |
|  | CBD | 8 | 5.13E-13 | 1.83E-13 | 6.48E-14 | 3.60E-13 | 6.66E-13 | 1.62E-13 | 7.56E-13 |
|  | THC | 10 | 6.55E-13 | 1.91E-13 | 6.04E-14 | 5.19E-13 | 7.92E-13 | 4.32E-13 | 1.05E-12 |
|  | THC+CBD | 9 | 5.53E-13 | 1.78E-13 | 5.94E-14 | 4.16E-13 | 6.90E-13 | 3.63E-13 | 8.35E-13 |
| *N-*oleoyl glycine | Vehicle | 8 | 6.03E-13 | 4.43E-13 | 1.56E-13 | 2.33E-13 | 9.73E-13 | 1.67E-13 | 1.44E-12 |
|  | CBD | 9 | 7.34E-13 | 2.45E-13 | 8.16E-14 | 5.46E-13 | 9.22E-13 | 2.98E-13 | 1.08E-12 |
|  | THC | 10 | 8.09E-13 | 3.33E-13 | 1.05E-13 | 5.71E-13 | 1.05E-12 | 3.80E-13 | 1.32E-12 |
|  | THC+CBD | 9 | 9.68E-13 | 4.63E-13 | 1.54E-13 | 6.12E-13 | 1.32E-12 | 5.20E-13 | 1.77E-12 |
| *N-*linoleoyl glycine | Vehicle | 7 | 1.08E-12 | 7.73E-13 | 2.92E-13 | 3.62E-13 | 1.79E-12 | 4.20E-13 | 2.51E-12 |
|  | CBD | 9 | 1.03E-12 | 2.94E-13 | 9.81E-14 | 8.01E-13 | 1.25E-12 | 6.35E-13 | 1.51E-12 |
|  | THC | 10 | 1.65E-12 | 6.86E-13 | 2.17E-13 | 1.16E-12 | 2.15E-12 | 6.82E-13 | 2.88E-12 |
|  | THC+CBD | 9 | 2.25E-12 | 6.60E-13 | 2.20E-13 | 1.74E-12 | 2.76E-12 | 1.44E-12 | 3.35E-12 |
| *N-*arachidonoyl glycine | Vehicle | BAL |  |  |  |  |  |  |  |
|  | CBD | BDL |  |  |  |  |  |  |  |
|  | THC | BAL |  |  |  |  |  |  |  |
|  | THC+CBD | BDL |  |  |  |  |  |  |  |
| *N-*docosahexaenoyl glycine | Vehicle | BAL |  |  |  |  |  |  |  |
|  | CBD | BAL |  |  |  |  |  |  |  |
|  | THC | BAL |  |  |  |  |  |  |  |
|  | THC+CBD | BAL |  |  |  |  |  |  |  |
| *N-*acyl leucines |  |  |  |  |  |  |  |  |  |
| *N-*palmitoyl leucine | Vehicle | 8 | 1.70E-12 | 5.55E-13 | 1.96E-13 | 1.23E-12 | 2.16E-12 | 8.87E-13 | 2.58E-12 |
|  | CBD | 9 | 2.01E-12 | 5.97E-13 | 1.99E-13 | 1.55E-12 | 2.47E-12 | 1.22E-12 | 2.81E-12 |
|  | THC | 11 | 2.33E-12 | 6.18E-13 | 1.86E-13 | 1.91E-12 | 2.74E-12 | 1.52E-12 | 3.52E-12 |
|  | THC+CBD | 9 | 1.95E-12 | 6.41E-13 | 2.14E-13 | 1.46E-12 | 2.44E-12 | 1.27E-12 | 3.00E-12 |
| *N-*stearoyl leucine | Vehicle | 8 | 6.24E-13 | 2.41E-13 | 8.52E-14 | 4.22E-13 | 8.25E-13 | 3.86E-13 | 1.09E-12 |
|  | CBD | 9 | 4.53E-13 | 1.88E-13 | 6.28E-14 | 3.08E-13 | 5.97E-13 | 2.08E-13 | 7.13E-13 |
|  | THC | 11 | 4.76E-13 | 1.73E-13 | 5.22E-14 | 3.60E-13 | 5.93E-13 | 2.23E-13 | 7.38E-13 |
|  | THC+CBD | 9 | 4.12E-13 | 2.15E-13 | 7.17E-14 | 2.46E-13 | 5.77E-13 | 2.04E-13 | 7.44E-13 |
| *N-*linoleoyl leucine | Vehicle | 8 | 3.20E-12 | 1.86E-12 | 6.57E-13 | 1.65E-12 | 4.76E-12 | 9.24E-13 | 6.80E-12 |
|  | CBD | 9 | 3.32E-12 | 1.22E-12 | 4.07E-13 | 2.38E-12 | 4.26E-12 | 1.90E-12 | 5.22E-12 |
|  | THC | 10 | 4.67E-12 | 1.34E-12 | 4.24E-13 | 3.71E-12 | 5.63E-12 | 2.79E-12 | 7.37E-12 |
|  | THC+CBD | 8 | 4.26E-12 | 1.22E-12 | 4.30E-13 | 3.24E-12 | 5.28E-12 | 3.31E-12 | 6.83E-12 |
| *N-*oleoyl leucine | Vehicle | 8 | 1.14E-12 | 5.92E-13 | 2.09E-13 | 6.47E-13 | 1.64E-12 | 2.68E-13 | 2.30E-12 |
|  | CBD | 9 | 1.35E-12 | 4.65E-13 | 1.55E-13 | 9.90E-13 | 1.70E-12 | 7.29E-13 | 2.13E-12 |
|  | THC | 11 | 1.61E-12 | 4.17E-13 | 1.26E-13 | 1.33E-12 | 1.89E-12 | 9.65E-13 | 2.13E-12 |
|  | THC+CBD | 9 | 1.60E-12 | 5.96E-13 | 1.99E-13 | 1.15E-12 | 2.06E-12 | 1.12E-12 | 2.65E-12 |
| *N-*docosahexaenoyl leucine | Vehicle | 8 | 1.28E-12 | 8.45E-13 | 2.99E-13 | 5.71E-13 | 1.98E-12 | 5.85E-13 | 3.26E-12 |
|  | CBD | 8 | 6.20E-13 | 9.28E-14 | 3.28E-14 | 5.42E-13 | 6.97E-13 | 4.83E-13 | 7.37E-13 |
|  | THC | 11 | 7.91E-13 | 3.41E-13 | 1.03E-13 | 5.62E-13 | 1.02E-12 | 4.48E-13 | 1.44E-12 |
|  | THC+CBD | 8 | 2.86E-13 | 1.54E-13 | 5.44E-14 | 1.57E-13 | 4.14E-13 | 1.81E-13 | 6.28E-13 |
| *N-*acyl methionines |  |  |  |  |  |  |  |  |  |
| *N-*palmitoyl methionine | Vehicle | 8 | 2.54E-12 | 9.32E-13 | 3.29E-13 | 1.76E-12 | 3.32E-12 | 1.40E-12 | 4.22E-12 |
|  | CBD | 8 | 1.87E-12 | 6.69E-13 | 2.36E-13 | 1.31E-12 | 2.43E-12 | 8.50E-13 | 2.67E-12 |
|  | THC | 11 | 1.79E-12 | 8.48E-13 | 2.56E-13 | 1.22E-12 | 2.36E-12 | 5.87E-13 | 3.36E-12 |
|  | THC+CBD | 8 | 1.75E-12 | 3.88E-13 | 1.37E-13 | 1.43E-12 | 2.08E-12 | 1.17E-12 | 2.36E-12 |
| *N-*stearoyl methionine | Vehicle | 8 | 2.74E-12 | 9.43E-13 | 3.33E-13 | 1.95E-12 | 3.52E-12 | 1.32E-12 | 4.22E-12 |
|  | CBD | 9 | 1.21E-12 | 4.04E-13 | 1.35E-13 | 8.96E-13 | 1.52E-12 | 7.81E-13 | 1.97E-12 |
|  | THC | 11 | 1.37E-12 | 3.76E-13 | 1.13E-13 | 1.12E-12 | 1.62E-12 | 9.01E-13 | 2.02E-12 |
|  | THC+CBD | 9 | 2.05E-12 | 4.70E-13 | 1.57E-13 | 1.69E-12 | 2.41E-12 | 1.51E-12 | 2.62E-12 |
| *N-*oleoyl methionine | Vehicle | 7 | 3.18E-12 | 6.78E-13 | 2.56E-13 | 2.56E-12 | 3.81E-12 | 2.00E-12 | 3.85E-12 |
|  | CBD | 8 | 1.88E-12 | 5.75E-13 | 2.03E-13 | 1.40E-12 | 2.36E-12 | 1.04E-12 | 2.75E-12 |
|  | THC | 11 | 1.79E-12 | 5.75E-13 | 1.73E-13 | 1.40E-12 | 2.18E-12 | 1.17E-12 | 2.86E-12 |
|  | THC+CBD | 9 | 2.45E-12 | 3.97E-13 | 1.32E-13 | 2.15E-12 | 2.76E-12 | 1.83E-12 | 3.03E-12 |
| *N-*linoleoyl methionine | Vehicle | 8 | 6.00E-12 | 1.63E-12 | 5.77E-13 | 4.63E-12 | 7.36E-12 | 4.48E-12 | 8.88E-12 |
|  | CBD | 8 | 3.82E-12 | 3.92E-13 | 1.39E-13 | 3.49E-12 | 4.14E-12 | 3.19E-12 | 4.38E-12 |
|  | THC | 11 | 5.15E-12 | 1.22E-12 | 3.68E-13 | 4.33E-12 | 5.97E-12 | 3.05E-12 | 6.93E-12 |
|  | THC+CBD | 9 | 9.38E-12 | 7.00E-13 | 2.33E-13 | 8.84E-12 | 9.91E-12 | 8.39E-12 | 1.05E-11 |
| *N-*arachidonoyl methionine | Vehicle | 8 | 2.29E-12 | 6.17E-13 | 2.18E-13 | 1.77E-12 | 2.80E-12 | 1.38E-12 | 3.08E-12 |
|  | CBD | 9 | 1.29E-12 | 3.01E-13 | 1.00E-13 | 1.06E-12 | 1.53E-12 | 8.46E-13 | 1.66E-12 |
|  | THC | 10 | 1.27E-12 | 4.56E-13 | 1.44E-13 | 9.44E-13 | 1.60E-12 | 6.00E-13 | 2.04E-12 |
|  | THC+CBD | 9 | 3.38E-12 | 6.55E-13 | 2.18E-13 | 2.88E-12 | 3.89E-12 | 2.73E-12 | 4.54E-12 |
| *N-*docosahexaenoyl methionine | Vehicle | BDL |  |  |  |  |  |  |  |
|  | CBD | BDL |  |  |  |  |  |  |  |
|  | THC | BDL |  |  |  |  |  |  |  |
|  | THC+CBD | BDL |  |  |  |  |  |  |  |
| *N-*acyl phenylalanines |  |  |  |  |  |  |  |  |  |
| *N-*palmitoyl phenylalanine | Vehicle | 7 | 9.45E-13 | 2.31E-13 | 8.73E-14 | 7.32E-13 | 1.16E-12 | 6.50E-13 | 1.21E-12 |
|  | CBD | 8 | 9.81E-13 | 2.82E-13 | 9.99E-14 | 7.45E-13 | 1.22E-12 | 6.77E-13 | 1.53E-12 |
|  | THC | 10 | 1.08E-12 | 1.83E-13 | 5.80E-14 | 9.49E-13 | 1.21E-12 | 8.03E-13 | 1.47E-12 |
|  | THC+CBD | 9 | 9.03E-13 | 2.12E-13 | 7.07E-14 | 7.40E-13 | 1.07E-12 | 5.79E-13 | 1.28E-12 |
| *N-*stearoyl phenylalanine | Vehicle | 8 | 2.97E-13 | 8.34E-14 | 2.95E-14 | 2.27E-13 | 3.66E-13 | 1.55E-13 | 4.29E-13 |
|  | CBD | 9 | 2.72E-13 | 4.53E-14 | 1.51E-14 | 2.37E-13 | 3.07E-13 | 1.98E-13 | 3.35E-13 |
|  | THC | 11 | 3.51E-13 | 7.92E-14 | 2.39E-14 | 2.98E-13 | 4.04E-13 | 2.51E-13 | 5.09E-13 |
|  | THC+CBD | 8 | 1.82E-13 | 6.85E-14 | 2.42E-14 | 1.25E-13 | 2.39E-13 | 1.28E-13 | 2.85E-13 |
| *N-*oleoyl phenylalanine | Vehicle | 8 | 6.09E-13 | 2.77E-13 | 9.81E-14 | 3.77E-13 | 8.41E-13 | 2.65E-13 | 1.04E-12 |
|  | CBD | 8 | 6.93E-13 | 1.30E-13 | 4.60E-14 | 5.84E-13 | 8.02E-13 | 5.30E-13 | 9.23E-13 |
|  | THC | 11 | 7.84E-13 | 1.92E-13 | 5.80E-14 | 6.55E-13 | 9.14E-13 | 5.01E-13 | 1.11E-12 |
|  | THC+CBD | 9 | 7.72E-13 | 1.98E-13 | 6.59E-14 | 6.20E-13 | 9.24E-13 | 5.38E-13 | 1.07E-12 |
| *N-*linoleoyl phenylalanine | Vehicle | 8 | 9.18E-13 | 3.51E-13 | 1.24E-13 | 6.24E-13 | 1.21E-12 | 4.98E-13 | 1.47E-12 |
|  | CBD | 9 | 1.20E-12 | 4.71E-13 | 1.57E-13 | 8.34E-13 | 1.56E-12 | 6.61E-13 | 1.85E-12 |
|  | THC | 10 | 1.31E-12 | 4.44E-13 | 1.40E-13 | 9.89E-13 | 1.62E-12 | 6.96E-13 | 2.02E-12 |
|  | THC+CBD | 9 | 1.54E-12 | 5.66E-13 | 1.89E-13 | 1.10E-12 | 1.97E-12 | 9.35E-13 | 2.57E-12 |
| *N-*arachidonoyl phenylalanine | Vehicle | 5 | 1.37E-13 | 8.38E-14 | 3.75E-14 | 3.30E-14 | 2.41E-13 | 4.90E-14 | 2.47E-13 |
|  | CBD | 6 | 1.09E-13 | 4.85E-14 | 1.98E-14 | 5.77E-14 | 1.59E-13 | 4.28E-14 | 1.59E-13 |
|  | THC | BAL |  |  |  |  |  |  |  |
|  | THC+CBD | 4 | 1.06E-13 | 3.44E-14 | 1.72E-14 | 5.11E-14 | 1.60E-13 | 7.35E-14 | 1.47E-13 |
| *N-*docosahexaenoyl phenylalanine | Vehicle | BAL |  |  |  |  |  |  |  |
|  | CBD | BAL |  |  |  |  |  |  |  |
|  | THC | BAL |  |  |  |  |  |  |  |
|  | THC+CBD | BDL |  |  |  |  |  |  |  |
| *N-*acyl prolines |  |  |  |  |  |  |  |  |  |
| *N-*palmitoyl proline | Vehicle | 8 | 6.97E-14 | 3.60E-14 | 1.27E-14 | 3.96E-14 | 9.98E-14 | 3.72E-14 | 1.46E-13 |
|  | CBD | 9 | 1.03E-13 | 4.03E-14 | 1.34E-14 | 7.16E-14 | 1.34E-13 | 3.41E-14 | 1.70E-13 |
|  | THC | 11 | 1.10E-13 | 3.59E-14 | 1.08E-14 | 8.62E-14 | 1.34E-13 | 3.86E-14 | 1.49E-13 |
|  | THC+CBD | 8 | 1.32E-13 | 3.71E-14 | 1.31E-14 | 1.01E-13 | 1.63E-13 | 8.91E-14 | 1.98E-13 |
| *N-*stearoyl proline | Vehicle | BAL |  |  |  |  |  |  |  |
|  | CBD | BDL |  |  |  |  |  |  |  |
|  | THC | BAL |  |  |  |  |  |  |  |
|  | THC+CBD | BDL |  |  |  |  |  |  |  |
| *N-*oleoyl proline | Vehicle | BAL |  |  |  |  |  |  |  |
|  | CBD | 3 | 7.32E-14 | 2.99E-14 | 1.72E-14 | -9.73E-16 | 1.47E-13 | 3.89E-14 | 9.32E-14 |
|  | THC | 4 | 7.14E-14 | 1.32E-14 | 6.62E-15 | 5.03E-14 | 9.25E-14 | 5.73E-14 | 8.50E-14 |
|  | THC+CBD | 5 | 8.00E-14 | 3.33E-14 | 1.49E-14 | 3.86E-14 | 1.21E-13 | 4.06E-14 | 1.15E-13 |
| *N-*linoleoyl proline | Vehicle | 4 | 2.08E-14 | 6.50E-15 | 3.25E-15 | 1.05E-14 | 3.11E-14 | 1.33E-14 | 2.89E-14 |
|  | CBD | 7 | 2.76E-14 | 8.99E-15 | 3.40E-15 | 1.93E-14 | 3.59E-14 | 1.43E-14 | 3.71E-14 |
|  | THC | 11 | 3.27E-14 | 8.74E-15 | 2.64E-15 | 2.68E-14 | 3.85E-14 | 2.48E-14 | 4.81E-14 |
|  | THC+CBD | 9 | 5.24E-14 | 2.06E-14 | 6.87E-15 | 3.65E-14 | 6.82E-14 | 3.10E-14 | 8.43E-14 |
| *N-*arachidonoyl proline | Vehicle | BDL |  |  |  |  |  |  |  |
|  | CBD | BDL |  |  |  |  |  |  |  |
|  | THC | BDL |  |  |  |  |  |  |  |
|  | THC+CBD | BDL |  |  |  |  |  |  |  |
| *N-*docosahexaenoyl proline | Vehicle | BDL |  |  |  |  |  |  |  |
|  | CBD | BDL |  |  |  |  |  |  |  |
|  | THC | BDL |  |  |  |  |  |  |  |
|  | THC+CBD | BDL |  |  |  |  |  |  |  |
| *N-*acyl serines |  |  |  |  |  |  |  |  |  |
| *N-*palmitoyl serine | Vehicle | 8 | 3.60E-12 | 1.68E-12 | 5.95E-13 | 2.20E-12 | 5.01E-12 | 2.10E-12 | 6.17E-12 |
|  | CBD | 9 | 3.64E-12 | 7.42E-13 | 2.47E-13 | 3.07E-12 | 4.21E-12 | 2.66E-12 | 4.56E-12 |
|  | THC | 11 | 6.00E-12 | 1.80E-12 | 5.43E-13 | 4.79E-12 | 7.21E-12 | 2.90E-12 | 8.48E-12 |
|  | THC+CBD | 9 | 3.97E-12 | 8.47E-13 | 2.82E-13 | 3.32E-12 | 4.63E-12 | 2.83E-12 | 5.18E-12 |
| *N-*stearoyl serine | Vehicle | 7 | 6.99E-13 | 2.53E-13 | 9.55E-14 | 4.65E-13 | 9.32E-13 | 2.66E-13 | 1.08E-12 |
|  | CBD | 9 | 5.56E-13 | 3.07E-13 | 1.02E-13 | 3.20E-13 | 7.91E-13 | 2.18E-13 | 1.03E-12 |
|  | THC | 10 | 5.09E-13 | 1.22E-13 | 3.85E-14 | 4.22E-13 | 5.96E-13 | 3.35E-13 | 6.85E-13 |
|  | THC+CBD | 8 | 2.64E-13 | 8.90E-14 | 3.15E-14 | 1.90E-13 | 3.39E-13 | 1.76E-13 | 3.98E-13 |
| *N-*oleoyl serine | Vehicle | 8 | 8.54E-13 | 4.18E-13 | 1.48E-13 | 5.04E-13 | 1.20E-12 | 3.74E-13 | 1.43E-12 |
|  | CBD | 8 | 1.01E-12 | 8.24E-14 | 2.91E-14 | 9.37E-13 | 1.08E-12 | 8.88E-13 | 1.12E-12 |
|  | THC | 11 | 1.73E-12 | 4.37E-13 | 1.32E-13 | 1.44E-12 | 2.02E-12 | 9.55E-13 | 2.28E-12 |
|  | THC+CBD | 9 | 1.60E-12 | 5.99E-13 | 2.00E-13 | 1.13E-12 | 2.06E-12 | 8.19E-13 | 2.48E-12 |
| *N-*linoleoyl serine | Vehicle | 8 | 2.86E-12 | 6.25E-13 | 2.21E-13 | 2.34E-12 | 3.39E-12 | 2.03E-12 | 3.82E-12 |
|  | CBD | 8 | 2.98E-12 | 3.57E-13 | 1.26E-13 | 2.69E-12 | 3.28E-12 | 2.34E-12 | 3.45E-12 |
|  | THC | 11 | 4.70E-12 | 1.32E-12 | 3.97E-13 | 3.81E-12 | 5.58E-12 | 3.00E-12 | 6.63E-12 |
|  | THC+CBD | 9 | 4.46E-12 | 1.33E-12 | 4.44E-13 | 3.44E-12 | 5.49E-12 | 2.62E-12 | 6.76E-12 |
| *N-*arachidonoyl serine | Vehicle | 3 | 1.34E-13 | 5.51E-14 | 3.18E-14 | -2.53E-15 | 2.71E-13 | 7.09E-14 | 1.71E-13 |
|  | CBD | 3 | 9.02E-14 | 3.39E-14 | 1.96E-14 | 5.90E-15 | 1.75E-13 | 5.40E-14 | 1.21E-13 |
|  | THC | 4 | 5.07E-14 | 2.31E-14 | 1.15E-14 | 1.39E-14 | 8.74E-14 | 3.26E-14 | 8.44E-14 |
|  | THC+CBD | 4 | 6.07E-14 | 2.45E-14 | 1.22E-14 | 2.17E-14 | 9.96E-14 | 4.42E-14 | 9.70E-14 |
| *N-*docosahexaenoyl serine | Vehicle | BDL |  |  |  |  |  |  |  |
|  | CBD | BDL |  |  |  |  |  |  |  |
|  | THC | BDL |  |  |  |  |  |  |  |
|  | THC+CBD | BDL |  |  |  |  |  |  |  |
| *N-*acyl taurines |  |  |  |  |  |  |  |  |  |
| *N-*palmitoyl taurine | Vehicle | 7 | 5.78E-11 | 5.20E-11 | 1.97E-11 | 9.72E-12 | 1.06E-10 | 2.05E-11 | 1.56E-10 |
|  | CBD | 7 | 2.17E-11 | 7.21E-12 | 2.73E-12 | 1.50E-11 | 2.83E-11 | 1.02E-11 | 3.26E-11 |
|  | THC | 10 | 3.34E-11 | 1.42E-11 | 4.49E-12 | 2.32E-11 | 4.35E-11 | 1.75E-11 | 5.89E-11 |
|  | THC+CBD | 9 | 4.86E-11 | 9.84E-12 | 3.28E-12 | 4.10E-11 | 5.61E-11 | 3.63E-11 | 6.28E-11 |
| *N-*stearoyl taurine | Vehicle | BAL |  |  |  |  |  |  |  |
|  | CBD | BAL |  |  |  |  |  |  |  |
|  | THC | 3 | 1.03E-11 | 4.15E-12 | 2.40E-12 | 3.69E-14 | 2.06E-11 | 7.49E-12 | 1.51E-11 |
|  | THC+CBD | BAL |  |  |  |  |  |  |  |
| *N-*oleoyl taurine | Vehicle | 8 | 2.79E-11 | 2.24E-11 | 7.93E-12 | 9.17E-12 | 4.67E-11 | 8.34E-12 | 7.17E-11 |
|  | CBD | 8 | 8.44E-12 | 1.44E-12 | 5.07E-13 | 7.24E-12 | 9.64E-12 | 6.70E-12 | 1.16E-11 |
|  | THC | 10 | 1.99E-11 | 6.40E-12 | 2.02E-12 | 1.54E-11 | 2.45E-11 | 9.47E-12 | 3.06E-11 |
|  | THC+CBD | 8 | 2.52E-11 | 2.90E-12 | 1.03E-12 | 2.28E-11 | 2.76E-11 | 2.15E-11 | 2.97E-11 |
| *N-*arachidonoyl taurine | Vehicle | 8 | 4.69E-11 | 2.99E-11 | 1.06E-11 | 2.19E-11 | 7.19E-11 | 2.07E-11 | 1.01E-10 |
|  | CBD | 7 | 1.46E-11 | 4.40E-12 | 1.66E-12 | 1.05E-11 | 1.87E-11 | 9.95E-12 | 2.06E-11 |
|  | THC | 10 | 3.61E-11 | 1.33E-11 | 4.22E-12 | 2.66E-11 | 4.56E-11 | 2.08E-11 | 6.25E-11 |
|  | THC+CBD | 9 | 9.93E-11 | 3.09E-11 | 1.03E-11 | 7.55E-11 | 1.23E-10 | 4.68E-11 | 1.36E-10 |
| *N-*acyl tryptophans |  |  |  |  |  |  |  |  |  |
| *N-*oleoyl tryptophan | Vehicle | 5 | 2.19E-12 | 1.07E-12 | 4.81E-13 | 8.54E-13 | 3.52E-12 | 8.57E-13 | 3.61E-12 |
|  | CBD | 7 | 2.05E-12 | 9.46E-13 | 3.57E-13 | 1.18E-12 | 2.92E-12 | 9.52E-13 | 3.41E-12 |
|  | THC | 9 | 1.57E-12 | 3.42E-13 | 1.14E-13 | 1.30E-12 | 1.83E-12 | 1.17E-12 | 2.08E-12 |
|  | THC+CBD | 9 | 1.67E-12 | 5.60E-13 | 1.87E-13 | 1.24E-12 | 2.10E-12 | 1.06E-12 | 2.60E-12 |
| *N-*palmitoyl tryptophan | Vehicle | 4 | 1.03E-12 | 4.51E-13 | 2.25E-13 | 3.11E-13 | 1.74E-12 | 5.31E-13 | 1.60E-12 |
|  | CBD | 8 | 1.23E-12 | 5.58E-13 | 1.97E-13 | 7.61E-13 | 1.69E-12 | 3.98E-13 | 1.90E-12 |
|  | THC | 10 | 1.03E-12 | 4.60E-13 | 1.45E-13 | 7.00E-13 | 1.36E-12 | 6.39E-13 | 2.02E-12 |
|  | THC+CBD | 8 | 9.11E-13 | 4.44E-13 | 1.57E-13 | 5.40E-13 | 1.28E-12 | 4.75E-13 | 1.63E-12 |
| *N-*arachidonoyl tryptophan | Vehicle | BDL |  |  |  |  |  |  |  |
|  | CBD | 3 | 8.40E-13 | 2.91E-13 | 1.68E-13 | 1.17E-13 | 1.56E-12 | 5.28E-13 | 1.10E-12 |
|  | THC | BAL |  |  |  |  |  |  |  |
|  | THC+CBD | BAL |  |  |  |  |  |  |  |
| *N-*stearoyl tryptophan | Vehicle | 7 | 2.16E-12 | 9.23E-13 | 3.49E-13 | 1.31E-12 | 3.01E-12 | 9.41E-13 | 3.69E-12 |
|  | CBD | 9 | 1.63E-12 | 3.94E-13 | 1.31E-13 | 1.33E-12 | 1.93E-12 | 9.52E-13 | 2.13E-12 |
|  | THC | 8 | 1.55E-12 | 7.27E-13 | 2.57E-13 | 9.43E-13 | 2.16E-12 | 5.87E-13 | 2.86E-12 |
|  | THC+CBD | 8 | 9.69E-13 | 2.97E-13 | 1.05E-13 | 7.21E-13 | 1.22E-12 | 5.98E-13 | 1.46E-12 |
| *N-*linoleoyl tryptophan | Vehicle | 8 | 3.05E-12 | 1.31E-12 | 4.63E-13 | 1.96E-12 | 4.15E-12 | 1.75E-12 | 5.55E-12 |
|  | CBD | 9 | 4.19E-12 | 1.15E-12 | 3.84E-13 | 3.31E-12 | 5.08E-12 | 2.46E-12 | 6.29E-12 |
|  | THC | 10 | 4.84E-12 | 9.95E-13 | 3.15E-13 | 4.12E-12 | 5.55E-12 | 2.81E-12 | 5.99E-12 |
|  | THC+CBD | 9 | 6.01E-12 | 1.94E-12 | 6.47E-13 | 4.51E-12 | 7.50E-12 | 2.97E-12 | 9.03E-12 |
| *N-*docosahexaenoyl tryptophan | Vehicle | 3 | 6.15E-13 | 6.83E-14 | 3.94E-14 | 4.45E-13 | 7.84E-13 | 5.45E-13 | 6.81E-13 |
|  | CBD | 7 | 4.92E-13 | 1.99E-13 | 7.54E-14 | 3.07E-13 | 6.76E-13 | 2.54E-13 | 8.39E-13 |
|  | THC | 4 | 4.21E-13 | 1.31E-13 | 6.55E-14 | 2.12E-13 | 6.29E-13 | 2.90E-13 | 5.77E-13 |
|  | THC+CBD | 4 | 4.06E-13 | 1.79E-13 | 8.97E-14 | 1.21E-13 | 6.92E-13 | 1.63E-13 | 5.59E-13 |
| *N-*acyl tyrosines |  |  |  |  |  |  |  |  |  |
| *N-*palmitoyl tyrosine | Vehicle | 7 | 9.26E-13 | 2.62E-13 | 9.91E-14 | 6.83E-13 | 1.17E-12 | 6.01E-13 | 1.23E-12 |
|  | CBD | 8 | 9.11E-13 | 1.33E-13 | 4.70E-14 | 8.00E-13 | 1.02E-12 | 6.38E-13 | 1.04E-12 |
|  | THC | 11 | 1.01E-12 | 3.57E-13 | 1.08E-13 | 7.65E-13 | 1.25E-12 | 6.18E-13 | 1.66E-12 |
|  | THC+CBD | 9 | 1.10E-12 | 1.35E-13 | 4.49E-14 | 9.97E-13 | 1.20E-12 | 8.44E-13 | 1.32E-12 |
| *N-*stearoyl tyrosine | Vehicle | 6 | 1.82E-13 | 6.83E-14 | 2.79E-14 | 1.11E-13 | 2.54E-13 | 1.05E-13 | 2.57E-13 |
|  | CBD | 8 | 1.21E-13 | 4.98E-14 | 1.76E-14 | 7.98E-14 | 1.63E-13 | 3.60E-14 | 2.10E-13 |
|  | THC | 10 | 1.27E-13 | 3.96E-14 | 1.25E-14 | 9.83E-14 | 1.55E-13 | 7.70E-14 | 2.01E-13 |
|  | THC+CBD | 7 | 1.06E-13 | 5.20E-14 | 1.97E-14 | 5.77E-14 | 1.54E-13 | 5.48E-14 | 1.98E-13 |
| *N-*oleoyl tyrosine | Vehicle | 8 | 8.42E-13 | 4.58E-13 | 1.62E-13 | 4.59E-13 | 1.22E-12 | 2.66E-13 | 1.62E-12 |
|  | CBD | 8 | 8.88E-13 | 2.94E-13 | 1.04E-13 | 6.42E-13 | 1.13E-12 | 5.21E-13 | 1.30E-12 |
|  | THC | 11 | 1.11E-12 | 5.34E-13 | 1.61E-13 | 7.49E-13 | 1.47E-12 | 5.60E-13 | 2.07E-12 |
|  | THC+CBD | 9 | 1.23E-12 | 2.33E-13 | 7.77E-14 | 1.05E-12 | 1.41E-12 | 9.21E-13 | 1.67E-12 |
| *N-*linoleoyl tyrosine | Vehicle | 8 | 1.22E-12 | 4.14E-13 | 1.46E-13 | 8.70E-13 | 1.56E-12 | 8.88E-13 | 2.06E-12 |
|  | CBD | 8 | 1.10E-12 | 1.33E-13 | 4.70E-14 | 9.91E-13 | 1.21E-12 | 9.69E-13 | 1.40E-12 |
|  | THC | 11 | 1.42E-12 | 5.09E-13 | 1.53E-13 | 1.08E-12 | 1.76E-12 | 7.77E-13 | 2.10E-12 |
|  | THC+CBD | 9 | 1.80E-12 | 3.00E-13 | 9.99E-14 | 1.57E-12 | 2.03E-12 | 1.44E-12 | 2.25E-12 |
| *N-*arachidonoyl tyrosine | Vehicle | BDL |  |  |  |  |  |  |  |
|  | CBD | BDL |  |  |  |  |  |  |  |
|  | THC | BDL |  |  |  |  |  |  |  |
|  | THC+CBD | BDL |  |  |  |  |  |  |  |
| *N-*docosahexaenoyl tyrosine | Vehicle | 3 | 1.43E-12 | 9.14E-13 | 5.28E-13 | -8.39E-13 | 3.70E-12 | 8.63E-13 | 2.49E-12 |
|  | CBD | BAL |  |  |  |  |  |  |  |
|  | THC | 4 | 6.78E-13 | 2.01E-13 | 1.00E-13 | 3.59E-13 | 9.98E-13 | 5.24E-13 | 9.73E-13 |
|  | THC+CBD | BAL |  |  |  |  |  |  |  |
| *N-*acyl valines |  |  |  |  |  |  |  |  |  |
| *N-*palmitoyl valine | Vehicle | 8 | 2.54E-12 | 9.31E-13 | 3.29E-13 | 1.77E-12 | 3.32E-12 | 1.35E-12 | 3.81E-12 |
|  | CBD | 8 | 2.95E-12 | 7.46E-13 | 2.64E-13 | 2.33E-12 | 3.57E-12 | 2.00E-12 | 4.26E-12 |
|  | THC | 11 | 3.47E-12 | 1.05E-12 | 3.18E-13 | 2.76E-12 | 4.18E-12 | 2.08E-12 | 5.44E-12 |
|  | THC+CBD | 9 | 3.30E-12 | 7.40E-13 | 2.47E-13 | 2.73E-12 | 3.87E-12 | 2.30E-12 | 4.44E-12 |
| *N-*linoleoyl valine | Vehicle | 8 | 3.99E-12 | 2.02E-12 | 7.13E-13 | 2.30E-12 | 5.68E-12 | 1.03E-12 | 6.12E-12 |
|  | CBD | 9 | 5.17E-12 | 2.11E-12 | 7.05E-13 | 3.54E-12 | 6.79E-12 | 2.24E-12 | 9.17E-12 |
|  | THC | 10 | 6.34E-12 | 4.87E-13 | 1.54E-13 | 5.99E-12 | 6.69E-12 | 5.76E-12 | 7.36E-12 |
|  | THC+CBD | 9 | 8.37E-12 | 2.64E-12 | 8.80E-13 | 6.34E-12 | 1.04E-11 | 6.23E-12 | 1.30E-11 |
| *N-*stearoyl valine | Vehicle | 8 | 6.23E-13 | 1.87E-13 | 6.62E-14 | 4.67E-13 | 7.80E-13 | 4.25E-13 | 9.45E-13 |
|  | CBD | 9 | 6.93E-13 | 2.28E-13 | 7.59E-14 | 5.18E-13 | 8.68E-13 | 2.71E-13 | 9.35E-13 |
|  | THC | 11 | 7.72E-13 | 1.88E-13 | 5.66E-14 | 6.46E-13 | 8.99E-13 | 4.91E-13 | 1.06E-12 |
|  | THC+CBD | 9 | 6.58E-13 | 3.73E-13 | 1.24E-13 | 3.72E-13 | 9.44E-13 | 3.37E-13 | 1.20E-12 |
| *N-*oleoyl valine | Vehicle | 8 | 9.60E-13 | 5.58E-13 | 1.97E-13 | 4.93E-13 | 1.43E-12 | 2.87E-13 | 1.65E-12 |
|  | CBD | 9 | 1.51E-12 | 4.73E-13 | 1.58E-13 | 1.15E-12 | 1.88E-12 | 8.44E-13 | 2.31E-12 |
|  | THC | 10 | 1.61E-12 | 3.17E-13 | 1.00E-13 | 1.38E-12 | 1.84E-12 | 1.17E-12 | 2.17E-12 |
|  | THC+CBD | 8 | 1.82E-12 | 4.27E-13 | 1.51E-13 | 1.47E-12 | 2.18E-12 | 1.36E-12 | 2.63E-12 |
| *N-*docosahexaenoyl valine | Vehicle | BAL |  |  |  |  |  |  |  |
|  | CBD | BDL |  |  |  |  |  |  |  |
|  | THC | BAL |  |  |  |  |  |  |  |
|  | THC+CBD | BDL |  |  |  |  |  |  |  |
| Free Fatty Acids |  |  |  |  |  |  |  |  |  |
| Oleic acid | Vehicle | 8 | 6.84E-08 | 3.62E-08 | 1.28E-08 | 3.82E-08 | 9.86E-08 | 3.11E-08 | 1.38E-07 |
|  | CBD | 9 | 3.69E-08 | 1.27E-08 | 4.24E-09 | 2.71E-08 | 4.67E-08 | 2.11E-08 | 5.56E-08 |
|  | THC | 11 | 3.37E-08 | 8.62E-09 | 2.60E-09 | 2.79E-08 | 3.94E-08 | 2.15E-08 | 4.45E-08 |
|  | T plus C | 8 | 3.59E-08 | 5.52E-09 | 1.95E-09 | 3.12E-08 | 4.05E-08 | 2.80E-08 | 4.33E-08 |
| Linoleoic acid | Vehicle | 8 | 1.39E-07 | 6.67E-08 | 2.36E-08 | 8.36E-08 | 1.95E-07 | 5.39E-08 | 2.19E-07 |
|  | CBD | 9 | 7.53E-08 | 2.78E-08 | 9.27E-09 | 5.39E-08 | 9.67E-08 | 3.76E-08 | 1.16E-07 |
|  | THC | 11 | 6.52E-08 | 1.78E-08 | 5.36E-09 | 5.33E-08 | 7.72E-08 | 3.59E-08 | 8.92E-08 |
|  | T plus C | 9 | 8.15E-08 | 1.77E-08 | 5.91E-09 | 6.78E-08 | 9.51E-08 | 5.66E-08 | 1.10E-07 |
| Arachidonic acid | Vehicle | 8 | 4.20E-08 | 1.90E-08 | 6.71E-09 | 2.62E-08 | 5.79E-08 | 1.52E-08 | 7.27E-08 |
|  | CBD | 9 | 2.52E-08 | 8.86E-09 | 2.95E-09 | 1.84E-08 | 3.20E-08 | 1.29E-08 | 3.67E-08 |
|  | THC | 11 | 1.88E-08 | 6.22E-09 | 1.88E-09 | 1.46E-08 | 2.30E-08 | 9.20E-09 | 2.74E-08 |
|  | T plus C | 9 | 2.15E-08 | 5.08E-09 | 1.69E-09 | 1.76E-08 | 2.54E-08 | 1.57E-08 | 2.91E-08 |
| Eicosapentaenoic acid | Vehicle | 8 | 7.83E-09 | 2.99E-09 | 1.06E-09 | 5.34E-09 | 1.03E-08 | 3.40E-09 | 1.31E-08 |
|  | CBD | 9 | 5.87E-09 | 9.87E-10 | 3.29E-10 | 5.11E-09 | 6.63E-09 | 4.16E-09 | 7.02E-09 |
|  | THC | 10 | 5.16E-09 | 1.10E-09 | 3.48E-10 | 4.37E-09 | 5.95E-09 | 3.52E-09 | 7.44E-09 |
|  | T plus C | 8 | 4.52E-09 | 5.17E-10 | 1.83E-10 | 4.09E-09 | 4.96E-09 | 4.03E-09 | 5.48E-09 |
| Docosahexaenoic acid | Vehicle | 8 | 7.60E-09 | 4.80E-09 | 1.70E-09 | 3.58E-09 | 1.16E-08 | 1.93E-09 | 1.83E-08 |
|  | CBD | 9 | 5.07E-09 | 2.11E-09 | 7.05E-10 | 3.45E-09 | 6.70E-09 | 2.26E-09 | 7.96E-09 |
|  | THC | 11 | 4.55E-09 | 1.38E-09 | 4.17E-10 | 3.61E-09 | 5.48E-09 | 2.26E-09 | 6.47E-09 |
|  | T plus C | 9 | 4.46E-09 | 1.33E-09 | 4.44E-10 | 3.44E-09 | 5.48E-09 | 2.62E-09 | 6.24E-09 |
| *N-*acyl-s*N-*glycerols |  |  |  |  |  |  |  |  |  |
| *N-*arachidonoyl-s*N-*glycerol | Vehicle | 8 | 2.15E-10 | 3.42E-11 | 1.21E-11 | 1.87E-10 | 2.44E-10 | 1.53E-10 | 2.68E-10 |
|  | CBD | 9 | 2.52E-10 | 3.42E-11 | 1.14E-11 | 2.26E-10 | 2.78E-10 | 1.99E-10 | 2.90E-10 |
|  | THC | 11 | 2.08E-10 | 3.07E-11 | 9.27E-12 | 1.87E-10 | 2.29E-10 | 1.59E-10 | 2.77E-10 |
|  | THC+CBD | 9 | 2.02E-10 | 1.94E-11 | 6.48E-12 | 1.87E-10 | 2.17E-10 | 1.71E-10 | 2.32E-10 |
| *N-*palmitoyl-s*N-*glycerol | Vehicle | 8 | 9.95E-09 | 1.33E-09 | 4.70E-10 | 8.84E-09 | 1.11E-08 | 7.51E-09 | 1.19E-08 |
|  | CBD | 9 | 1.18E-08 | 1.67E-09 | 5.56E-10 | 1.05E-08 | 1.31E-08 | 8.70E-09 | 1.40E-08 |
|  | THC | 11 | 1.04E-08 | 1.66E-09 | 5.00E-10 | 9.28E-09 | 1.15E-08 | 7.68E-09 | 1.38E-08 |
|  | THC+CBD | 9 | 9.53E-09 | 1.17E-09 | 3.90E-10 | 8.63E-09 | 1.04E-08 | 7.58E-09 | 1.14E-08 |
| *N-*oleoyl-s*N-*glycerol | Vehicle | 8 | 7.91E-10 | 1.32E-10 | 4.66E-11 | 6.81E-10 | 9.02E-10 | 5.45E-10 | 9.38E-10 |
|  | CBD | 9 | 9.76E-10 | 1.35E-10 | 4.49E-11 | 8.73E-10 | 1.08E-09 | 7.51E-10 | 1.14E-09 |
|  | THC | 11 | 9.47E-10 | 1.60E-10 | 4.82E-11 | 8.39E-10 | 1.05E-09 | 6.97E-10 | 1.25E-09 |
|  | THC+CBD | 9 | 9.56E-10 | 1.10E-10 | 3.67E-11 | 8.71E-10 | 1.04E-09 | 7.94E-10 | 1.10E-09 |
| *N-*linoleoyl-s*N-*glycerol | Vehicle | 8 | 1.80E-09 | 3.40E-10 | 1.20E-10 | 1.52E-09 | 2.09E-09 | 1.27E-09 | 2.40E-09 |
|  | CBD | 9 | 1.70E-09 | 2.35E-10 | 7.83E-11 | 1.52E-09 | 1.88E-09 | 1.32E-09 | 1.98E-09 |
|  | THC | 11 | 1.68E-09 | 2.39E-10 | 7.20E-11 | 1.52E-09 | 1.84E-09 | 1.28E-09 | 2.19E-09 |
|  | THC+CBD | 9 | 1.74E-09 | 1.56E-10 | 5.22E-11 | 1.62E-09 | 1.86E-09 | 1.48E-09 | 1.96E-09 |
| Prostaglandins |  |  |  |  |  |  |  |  |  |
| PGE_2_ | Vehicle | BDL |  |  |  |  |  |  |  |
|  | CBD | BDL |  |  |  |  |  |  |  |
|  | THC | BDL |  |  |  |  |  |  |  |
|  | THC+CBD | BDL |  |  |  |  |  |  |  |
| PGF_2α_ | Vehicle | BDL |  |  |  |  |  |  |  |
|  | CBD | BDL |  |  |  |  |  |  |  |
|  | THC | BDL |  |  |  |  |  |  |  |
|  | THC+CBD | BDL |  |  |  |  |  |  |  |
| 6-keto-PGF_1α_ | Vehicle | BDL |  |  |  |  |  |  |  |
|  | CBD | BDL |  |  |  |  |  |  |  |
|  | THC | BDL |  |  |  |  |  |  |  |
|  | THC+CBD | BDL |  |  |  |  |  |  |  |

### Supplemental Figure 3. ANOVA for the effect of CBD, THC, or THC+CBD (3mg/kg) injections administered to a dam on *N-*acyl alanines measured in milk collected from nursing pups’ stomachs

|  | | Sum of Squares | df | Mean Square | F | Sig. |
| --- | --- | --- | --- | --- | --- | --- |
| *N-*palmitoyl alanine | Between Groups | 0.000 | 3 | 0.000 | 0.135 | 0.938 |
|  | Within Groups | 0.000 | 32 | 0.000 |  |  |
|  | Total | 0.000 | 35 |  |  |  |
| *N-*stearoyl alanine | Between Groups | 0.000 | 3 | 0.000 | 2.844 | 0.054 |
|  | Within Groups | 0.000 | 30 | 0.000 |  |  |
|  | Total | 0.000 | 33 |  |  |  |
| *N-*oleoyl alanine | Between Groups | 0.000 | 3 | 0.000 | 0.462 | 0.711 |
|  | Within Groups | 0.000 | 31 | 0.000 |  |  |
|  | Total | 0.000 | 34 |  |  |  |
| *N-*linoleoyl alanine | Between Groups | 0.000 | 3 | 0.000 | 24.762 | 0.000 |
|  | Within Groups | 0.000 | 31 | 0.000 |  |  |
|  | Total | 0.000 | 34 |  |  |  |

### Supplemental Figure 4. Post-hoc Fisher’s LSD results for *N-*acyl alanines

| Dependent Variable | | | Mean Difference (I-J) | Std. Error | Sig. | 95% Confidence Interval | |  |
| --- | --- | --- | --- | --- | --- | --- | --- | --- |
|  |  |  |  |  |  | Lower Bound | Upper Bound | Magnitude (drug/vehicle) |
| *N-*palmitoyl alanine | Vehicle | CBD | -2.20E-13 | 8.19E-13 | 0.790 | -1.89E-12 | 1.45E-12 |  |
|  |  | THC | 2.47E-13 | 7.83E-13 | 0.754 | -1.35E-12 | 1.84E-12 |  |
|  |  | THC+CBD | -7.65E-14 | 8.43E-13 | 0.928 | -1.79E-12 | 1.64E-12 |  |
|  | CBD | Vehicle | 2.20E-13 | 8.19E-13 | 0.790 | -1.45E-12 | 1.89E-12 |  |
|  |  | THC | 4.67E-13 | 7.58E-13 | 0.542 | -1.08E-12 | 2.01E-12 |  |
|  |  | THC+CBD | 1.43E-13 | 8.19E-13 | 0.862 | -1.53E-12 | 1.81E-12 |  |
|  | THC | Vehicle | -2.47E-13 | 7.83E-13 | 0.754 | -1.84E-12 | 1.35E-12 |  |
|  |  | CBD | -4.67E-13 | 7.58E-13 | 0.542 | -2.01E-12 | 1.08E-12 |  |
|  |  | THC+CBD | -3.24E-13 | 7.83E-13 | 0.682 | -1.92E-12 | 1.27E-12 |  |
|  | THC+CBD | Vehicle | 7.65E-14 | 8.43E-13 | 0.928 | -1.64E-12 | 1.79E-12 |  |
|  |  | CBD | -1.43E-13 | 8.19E-13 | 0.862 | -1.81E-12 | 1.53E-12 |  |
|  |  | THC | 3.24E-13 | 7.83E-13 | 0.682 | -1.27E-12 | 1.92E-12 |  |
| *N-*stearoyl alanine | Vehicle | CBD | 8.00E-13 | 4.58E-13 | 0.090 | -1.34E-13 | 1.74E-12 | 0.72 |
|  |  | THC | 1.20593E-012^*^ | 4.34E-13 | 0.009 | 3.19E-13 | 2.09E-12 | 0.58 |
|  |  | THC+CBD | 1.02024E-012^*^ | 4.58E-13 | 0.033 | 8.57E-14 | 1.95E-12 | 0.64 |
|  | CBD | Vehicle | -8.00E-13 | 4.58E-13 | 0.090 | -1.74E-12 | 1.34E-13 |  |
|  |  | THC | 4.05E-13 | 4.34E-13 | 0.358 | -4.81E-13 | 1.29E-12 |  |
|  |  | THC+CBD | 2.20E-13 | 4.58E-13 | 0.635 | -7.15E-13 | 1.15E-12 |  |
|  | THC | Vehicle | -1.20593E-012^*^ | 4.34E-13 | 0.009 | -2.09E-12 | -3.19E-13 |  |
|  |  | CBD | -4.05E-13 | 4.34E-13 | 0.358 | -1.29E-12 | 4.81E-13 |  |
|  |  | THC+CBD | -1.86E-13 | 4.34E-13 | 0.672 | -1.07E-12 | 7.01E-13 |  |
|  | THC+CBD | Vehicle | -1.02024E-012^*^ | 4.58E-13 | 0.033 | -1.95E-12 | -8.57E-14 |  |
|  |  | CBD | -2.20E-13 | 4.58E-13 | 0.635 | -1.15E-12 | 7.15E-13 |  |
|  |  | THC | 1.86E-13 | 4.34E-13 | 0.672 | -7.01E-13 | 1.07E-12 |  |
| *N-*oleoyl alanine | Vehicle | CBD | -1.04E-13 | 2.55E-13 | 0.685 | -6.25E-13 | 4.16E-13 |  |
|  |  | THC | 1.60E-13 | 2.45E-13 | 0.519 | -3.40E-13 | 6.59E-13 |  |
|  |  | THC+CBD | 1.76E-14 | 2.62E-13 | 0.947 | -5.17E-13 | 5.52E-13 |  |
|  | CBD | Vehicle | 1.04E-13 | 2.55E-13 | 0.685 | -4.16E-13 | 6.25E-13 |  |
|  |  | THC | 2.64E-13 | 2.28E-13 | 0.255 | -2.00E-13 | 7.29E-13 |  |
|  |  | THC+CBD | 1.22E-13 | 2.46E-13 | 0.623 | -3.80E-13 | 6.24E-13 |  |
|  | THC | Vehicle | -1.60E-13 | 2.45E-13 | 0.519 | -6.59E-13 | 3.40E-13 |  |
|  |  | CBD | -2.64E-13 | 2.28E-13 | 0.255 | -7.29E-13 | 2.00E-13 |  |
|  |  | THC+CBD | -1.42E-13 | 2.35E-13 | 0.550 | -6.22E-13 | 3.38E-13 |  |
|  | THC+CBD | Vehicle | -1.76E-14 | 2.62E-13 | 0.947 | -5.52E-13 | 5.17E-13 |  |
|  |  | CBD | -1.22E-13 | 2.46E-13 | 0.623 | -6.24E-13 | 3.80E-13 |  |
|  |  | THC | 1.42E-13 | 2.35E-13 | 0.550 | -3.38E-13 | 6.22E-13 |  |
| *N-*linoleoyl alanine | Vehicle | CBD | -1.26400E-012^*^ | 3.58E-13 | 0.001 | -1.99E-12 | -5.34E-13 | 1.78 |
|  |  | THC | -2.13111E-012^*^ | 3.39E-13 | 0.000 | -2.82E-12 | -1.44E-12 | 2.31 |
|  |  | THC+CBD | -2.84018E-012^*^ | 3.48E-13 | 0.000 | -3.55E-12 | -2.13E-12 | 2.74 |
|  | CBD | Vehicle | 1.26400E-012^*^ | 3.58E-13 | 0.001 | 5.34E-13 | 1.99E-12 |  |
|  |  | THC | -8.67108E-013^*^ | 3.39E-13 | 0.016 | -1.56E-12 | -1.75E-13 |  |
|  |  | THC+CBD | -1.57617E-012^*^ | 3.48E-13 | 0.000 | -2.29E-12 | -8.67E-13 |  |
|  | THC | Vehicle | 2.13111E-012^*^ | 3.39E-13 | 0.000 | 1.44E-12 | 2.82E-12 |  |
|  |  | CBD | 8.67108E-013^*^ | 3.39E-13 | 0.016 | 1.75E-13 | 1.56E-12 |  |
|  |  | THC+CBD | -7.09067E-013^*^ | 3.29E-13 | 0.039 | -1.38E-12 | -3.87E-14 |  |
|  | THC+CBD | Vehicle | 2.84018E-012^*^ | 3.48E-13 | 0.000 | 2.13E-12 | 3.55E-12 |  |
|  |  | CBD | 1.57617E-012^*^ | 3.48E-13 | 0.000 | 8.67E-13 | 2.29E-12 |  |
|  |  | THC | 7.09067E-013^*^ | 3.29E-13 | 0.039 | 3.87E-14 | 1.38E-12 |  |

### Supplemental Figure 5. ANOVA for the effect of CBD, THC, or THC+CBD (3mg/kg) injections administered to a dam on levels of *N-*acyl ethanolamines measured in milk collected from nursing pups’ stomachs

|  | | Sum of Squares | df | Mean Square | F | Sig. |
| --- | --- | --- | --- | --- | --- | --- |
| *N-*palmitoyl ethanolamine | Between Groups | 0.000 | 3 | 0.000 | 57.615 | 0.000 |
|  | Within Groups | 0.000 | 32 | 0.000 |  |  |
|  | Total | 0.000 | 35 |  |  |  |
| *N-*stearoyl ethanolamine | Between Groups | 0.000 | 3 | 0.000 | 72.377 | 0.000 |
|  | Within Groups | 0.000 | 32 | 0.000 |  |  |
|  | Total | 0.000 | 35 |  |  |  |
| *N-*oleoyl ethanolamine | Between Groups | 0.000 | 3 | 0.000 | 7.742 | 0.001 |
|  | Within Groups | 0.000 | 32 | 0.000 |  |  |
|  | Total | 0.000 | 35 |  |  |  |
| *N-*linoleoyl ethanolamine | Between Groups | 0.000 | 3 | 0.000 | 9.018 | 0.000 |
|  | Within Groups | 0.000 | 33 | 0.000 |  |  |
|  | Total | 0.000 | 36 |  |  |  |
| *N-*arachidonoyl ethanolamine | Between Groups | 0.000 | 3 | 0.000 | 6.246 | 0.002 |
|  | Within Groups | 0.000 | 33 | 0.000 |  |  |
|  | Total | 0.000 | 36 |  |  |  |
| *N-*docosahexaenoyl ethanolamine | Between Groups | 0.000 | 3 | 0.000 | 7.105 | 0.001 |
|  | Within Groups | 0.000 | 33 | 0.000 |  |  |
|  | Total | 0.000 | 36 |  |  |  |

### Supplemental Figure 6. Post-hoc Fisher’s LSD results for *N-*acyl ethanolamines

| Dependent Variable | | | Mean Difference (I-J) | Std. Error | Sig. | 95% Confidence Interval | |  |
| --- | --- | --- | --- | --- | --- | --- | --- | --- |
|  |  |  |  |  |  | Lower Bound | Upper Bound | Magnitude (drug/vehicle) |
| *N-*palmitoyl ethanolamine | Vehicle | CBD | 4.66157E-011^*^ | 4.28E-12 | 0.000 | 3.79E-11 | 5.53E-11 | 0.54 |
|  |  | THC | 4.77378E-011^*^ | 4.18E-12 | 0.000 | 3.92E-11 | 5.62E-11 | 0.53 |
|  |  | THC+CBD | 4.44477E-011^*^ | 4.28E-12 | 0.000 | 3.57E-11 | 5.32E-11 | 0.56 |
|  | CBD | Vehicle | -4.66157E-011^*^ | 4.28E-12 | 0.000 | -5.53E-11 | -3.79E-11 |  |
|  |  | THC | 1.12E-12 | 4.05E-12 | 0.783 | -7.12E-12 | 9.36E-12 |  |
|  |  | THC+CBD | -2.17E-12 | 4.15E-12 | 0.605 | -1.06E-11 | 6.29E-12 |  |
|  | THC | Vehicle | -4.77378E-011^*^ | 4.18E-12 | 0.000 | -5.62E-11 | -3.92E-11 |  |
|  |  | CBD | -1.12E-12 | 4.05E-12 | 0.783 | -9.36E-12 | 7.12E-12 |  |
|  |  | THC+CBD | -3.29E-12 | 4.05E-12 | 0.422 | -1.15E-11 | 4.95E-12 |  |
|  | THC+CBD | Vehicle | -4.44477E-011^*^ | 4.28E-12 | 0.000 | -5.32E-11 | -3.57E-11 |  |
|  |  | CBD | 2.17E-12 | 4.15E-12 | 0.605 | -6.29E-12 | 1.06E-11 |  |
|  |  | THC | 3.29E-12 | 4.05E-12 | 0.422 | -4.95E-12 | 1.15E-11 |  |
| *N-*stearoyl ethanolamine | Vehicle | CBD | 1.00623E-010^*^ | 8.57E-12 | 0.000 | 8.32E-11 | 1.18E-10 | 0.51 |
|  |  | THC | 9.59090E-011^*^ | 8.37E-12 | 0.000 | 7.89E-11 | 1.13E-10 | 0.53 |
|  |  | THC+CBD | 1.13296E-010^*^ | 8.57E-12 | 0.000 | 9.58E-11 | 1.31E-10 | 0.45 |
|  | CBD | Vehicle | -1.00623E-010^*^ | 8.57E-12 | 0.000 | -1.18E-10 | -8.32E-11 |  |
|  |  | THC | -4.71E-12 | 8.10E-12 | 0.565 | -2.12E-11 | 1.18E-11 |  |
|  |  | THC+CBD | 1.27E-11 | 8.31E-12 | 0.137 | -4.26E-12 | 2.96E-11 |  |
|  | THC | Vehicle | -9.59090E-011^*^ | 8.37E-12 | 0.000 | -1.13E-10 | -7.89E-11 |  |
|  |  | CBD | 4.71E-12 | 8.10E-12 | 0.565 | -1.18E-11 | 2.12E-11 |  |
|  |  | THC+CBD | 1.73865E-011^*^ | 8.10E-12 | 0.040 | 8.81E-13 | 3.39E-11 |  |
|  | THC+CBD | Vehicle | -1.13296E-010^*^ | 8.57E-12 | 0.000 | -1.31E-10 | -9.58E-11 |  |
|  |  | CBD | -1.27E-11 | 8.31E-12 | 0.137 | -2.96E-11 | 4.26E-12 |  |
|  |  | THC | -1.73865E-011^*^ | 8.10E-12 | 0.040 | -3.39E-11 | -8.81E-13 |  |
| *N-*oleoyl ethanolamine | Vehicle | CBD | 8.63193E-012^*^ | 2.08E-12 | 0.000 | 4.39E-12 | 1.29E-11 | 0.76 |
|  |  | THC | 5.18E-13 | 2.03E-12 | 0.801 | -3.63E-12 | 4.66E-12 |  |
|  |  | THC+CBD | 4.16E-12 | 2.08E-12 | 0.054 | -8.24E-14 | 8.41E-12 | 0.89 |
|  | CBD | Vehicle | -8.63193E-012^*^ | 2.08E-12 | 0.000 | -1.29E-11 | -4.39E-12 |  |
|  |  | THC | -8.11443E-012^*^ | 1.97E-12 | 0.000 | -1.21E-11 | -4.10E-12 |  |
|  |  | THC+CBD | -4.47024E-012^*^ | 2.02E-12 | 0.034 | -8.59E-12 | -3.53E-13 |  |
|  | THC | Vehicle | -5.18E-13 | 2.03E-12 | 0.801 | -4.66E-12 | 3.63E-12 |  |
|  |  | CBD | 8.11443E-012^*^ | 1.97E-12 | 0.000 | 4.10E-12 | 1.21E-11 |  |
|  |  | THC+CBD | 3.64E-12 | 1.97E-12 | 0.074 | -3.69E-13 | 7.66E-12 |  |
|  | THC+CBD | Vehicle | -4.16E-12 | 2.08E-12 | 0.054 | -8.41E-12 | 8.24E-14 |  |
|  |  | CBD | 4.47024E-012^*^ | 2.02E-12 | 0.034 | 3.53E-13 | 8.59E-12 |  |
|  |  | THC | -3.64E-12 | 1.97E-12 | 0.074 | -7.66E-12 | 3.69E-13 |  |
| *N-*linoleoyl ethanolamine | Vehicle | CBD | -9.54E-13 | 3.66E-12 | 0.796 | -8.40E-12 | 6.49E-12 |  |
|  |  | THC | -8.68428E-012^*^ | 3.50E-12 | 0.018 | -1.58E-11 | -1.57E-12 | 1.26 |
|  |  | THC+CBD | -1.63280E-011^*^ | 3.66E-12 | 0.000 | -2.38E-11 | -8.89E-12 | 1.50 |
|  | CBD | Vehicle | 9.54E-13 | 3.66E-12 | 0.796 | -6.49E-12 | 8.40E-12 |  |
|  |  | THC | -7.72994E-012^*^ | 3.38E-12 | 0.029 | -1.46E-11 | -8.45E-13 |  |
|  |  | THC+CBD | -1.53737E-011^*^ | 3.55E-12 | 0.000 | -2.26E-11 | -8.15E-12 |  |
|  | THC | Vehicle | 8.68428E-012^*^ | 3.50E-12 | 0.018 | 1.57E-12 | 1.58E-11 |  |
|  |  | CBD | 7.72994E-012^*^ | 3.38E-12 | 0.029 | 8.45E-13 | 1.46E-11 |  |
|  |  | THC+CBD | -7.64372E-012^*^ | 3.38E-12 | 0.031 | -1.45E-11 | -7.59E-13 |  |
|  | THC+CBD | Vehicle | 1.63280E-011^*^ | 3.66E-12 | 0.000 | 8.89E-12 | 2.38E-11 |  |
|  |  | CBD | 1.53737E-011^*^ | 3.55E-12 | 0.000 | 8.15E-12 | 2.26E-11 |  |
|  |  | THC | 7.64372E-012^*^ | 3.38E-12 | 0.031 | 7.59E-13 | 1.45E-11 |  |
| *N-*arachidonoyl ethanolamine | Vehicle | CBD | 3.71E-13 | 2.80E-13 | 0.194 | -1.98E-13 | 9.40E-13 |  |
|  |  | THC | 5.89442E-013^*^ | 2.67E-13 | 0.035 | 4.54E-14 | 1.13E-12 | 0.82 |
|  |  | THC+CBD | 1.17179E-012^*^ | 2.80E-13 | 0.000 | 6.03E-13 | 1.74E-12 | 0.65 |
|  | CBD | Vehicle | -3.71E-13 | 2.80E-13 | 0.194 | -9.40E-13 | 1.98E-13 |  |
|  |  | THC | 2.19E-13 | 2.59E-13 | 0.404 | -3.08E-13 | 7.45E-13 |  |
|  |  | THC+CBD | 8.00922E-013^*^ | 2.71E-13 | 0.006 | 2.49E-13 | 1.35E-12 |  |
|  | THC | Vehicle | -5.89442E-013^*^ | 2.67E-13 | 0.035 | -1.13E-12 | -4.54E-14 |  |
|  |  | CBD | -2.19E-13 | 2.59E-13 | 0.404 | -7.45E-13 | 3.08E-13 |  |
|  |  | THC+CBD | 5.82346E-013^*^ | 2.59E-13 | 0.031 | 5.61E-14 | 1.11E-12 |  |
|  | THC+CBD | Vehicle | -1.17179E-012^*^ | 2.80E-13 | 0.000 | -1.74E-12 | -6.03E-13 |  |
|  |  | CBD | -8.00922E-013^*^ | 2.71E-13 | 0.006 | -1.35E-12 | -2.49E-13 |  |
|  |  | THC | -5.82346E-013^*^ | 2.59E-13 | 0.031 | -1.11E-12 | -5.61E-14 |  |
| *N-*docosahexaenoyl ethanolamine | Vehicle | CBD | 7.51323E-012^*^ | 1.97E-12 | 0.001 | 3.51E-12 | 1.15E-11 | 0.70 |
|  |  | THC | 2.59E-12 | 1.88E-12 | 0.177 | -1.23E-12 | 6.42E-12 |  |
|  |  | THC+CBD | 7.20729E-012^*^ | 1.97E-12 | 0.001 | 3.21E-12 | 1.12E-11 | 0.72 |
|  | CBD | Vehicle | -7.51323E-012^*^ | 1.97E-12 | 0.001 | -1.15E-11 | -3.51E-12 |  |
|  |  | THC | -4.91975E-012^*^ | 1.82E-12 | 0.011 | -8.62E-12 | -1.22E-12 |  |
|  |  | THC+CBD | -3.06E-13 | 1.91E-12 | 0.874 | -4.19E-12 | 3.57E-12 |  |
|  | THC | Vehicle | -2.59E-12 | 1.88E-12 | 0.177 | -6.42E-12 | 1.23E-12 |  |
|  |  | CBD | 4.91975E-012^*^ | 1.82E-12 | 0.011 | 1.22E-12 | 8.62E-12 |  |
|  |  | THC+CBD | 4.61381E-012^*^ | 1.82E-12 | 0.016 | 9.14E-13 | 8.31E-12 |  |
|  | THC+CBD | Vehicle | -7.20729E-012^*^ | 1.97E-12 | 0.001 | -1.12E-11 | -3.21E-12 |  |
|  |  | CBD | 3.06E-13 | 1.91E-12 | 0.874 | -3.57E-12 | 4.19E-12 |  |
|  |  | THC | -4.61381E-012^*^ | 1.82E-12 | 0.016 | -8.31E-12 | -9.14E-13 |  |

### Supplemental Figure 7. ANOVA for the effect of CBD, THC, or THC+CBD (3mg/kg) injections administered to a dam on *N-*acyl GABAs measured in milk collected from nursing pups’ stomachs

|  | | Sum of Squares | df | Mean Square | F | Sig. |
| --- | --- | --- | --- | --- | --- | --- |
| *N-*palmitoyl GABA | Between Groups | 0.000 | 3 | 0.000 | 0.929 | 0.445 |
|  | Within Groups | 0.000 | 20 | 0.000 |  |  |
|  | Total | 0.000 | 23 |  |  |  |

### Supplemental Figure 8. Post-hoc Fisher’s LSD results for *N-*acyl GABAs

| Dependent Variable | | | Mean Difference (I-J) | Std. Error | Sig. | 95% Confidence Interval | |  |
| --- | --- | --- | --- | --- | --- | --- | --- | --- |
|  |  |  |  |  |  | Lower Bound | Upper Bound | Magnitude (drug/vehicle) |
| *N-*palmitoyl GABA | Vehicle | CBD | 1.44E-13 | 2.00E-13 | 0.481 | -2.74E-13 | 5.62E-13 |  |
|  |  | THC | 2.01E-13 | 2.05E-13 | 0.339 | -2.27E-13 | 6.29E-13 |  |
|  |  | THC+CBD | -3.15E-14 | 1.96E-13 | 0.874 | -4.41E-13 | 3.78E-13 |  |
|  | CBD | Vehicle | -1.44E-13 | 2.00E-13 | 0.481 | -5.62E-13 | 2.74E-13 |  |
|  |  | THC | 5.71E-14 | 1.61E-13 | 0.727 | -2.80E-13 | 3.94E-13 |  |
|  |  | THC+CBD | -1.75E-13 | 1.50E-13 | 0.257 | -4.89E-13 | 1.38E-13 |  |
|  | THC | Vehicle | -2.01E-13 | 2.05E-13 | 0.339 | -6.29E-13 | 2.27E-13 |  |
|  |  | CBD | -5.71E-14 | 1.61E-13 | 0.727 | -3.94E-13 | 2.80E-13 |  |
|  |  | THC+CBD | -2.32E-13 | 1.57E-13 | 0.154 | -5.59E-13 | 9.45E-14 |  |
|  | THC+CBD | Vehicle | 3.15E-14 | 1.96E-13 | 0.874 | -3.78E-13 | 4.41E-13 |  |
|  |  | CBD | 1.75E-13 | 1.50E-13 | 0.257 | -1.38E-13 | 4.89E-13 |  |
|  |  | THC | 2.32E-13 | 1.57E-13 | 0.154 | -9.45E-14 | 5.59E-13 |  |

### Supplemental Figure 9. ANOVA for the effect of CBD, THC, or THC+CBD (3mg/kg) injections administered to a dam on *N-*acyl glycines measured in milk collected from nursing pups’ stomachs

|  | | Sum of Squares | df | Mean Square | F | Sig. |
| --- | --- | --- | --- | --- | --- | --- |
| *N-*palmitoyl glycine | Between Groups | 0.000 | 3 | 0.000 | 2.435 | 0.083 |
|  | Within Groups | 0.000 | 31 | 0.000 |  |  |
|  | Total | 0.000 | 34 |  |  |  |
| *N-*linoleoyl glycine | Between Groups | 0.000 | 3 | 0.000 | 7.331 | 0.001 |
|  | Within Groups | 0.000 | 31 | 0.000 |  |  |
|  | Total | 0.000 | 34 |  |  |  |
| *N-*stearoyl glycine | Between Groups | 0.000 | 3 | 0.000 | 2.582 | 0.072 |
|  | Within Groups | 0.000 | 30 | 0.000 |  |  |
|  | Total | 0.000 | 33 |  |  |  |
| *N-*oleoyl glycine | Between Groups | 0.000 | 3 | 0.000 | 1.394 | 0.263 |
|  | Within Groups | 0.000 | 32 | 0.000 |  |  |
|  | Total | 0.000 | 35 |  |  |  |

### Supplemental Figure 10. Post-hoc Fisher’s LSD results for *N-*acyl glycines

| Dependent Variable | | | Mean Difference (I-J) | Std. Error | Sig. | 95% Confidence Interval | |  |
| --- | --- | --- | --- | --- | --- | --- | --- | --- |
|  |  |  |  |  |  | Lower Bound | Upper Bound | Magnitude (drug/vehicle) |
| *N-*palmitoyl glycine | Vehicle | CBD | 1.07E-13 | 7.05E-13 | 0.880 | -1.33E-12 | 1.55E-12 |  |
|  |  | THC | -1.06E-12 | 6.69E-13 | 0.124 | -2.42E-12 | 3.06E-13 |  |
|  |  | THC+CBD | -1.39E-12 | 6.86E-13 | 0.051 | -2.79E-12 | 9.40E-15 | 1.25 |
|  | CBD | Vehicle | -1.07E-13 | 7.05E-13 | 0.880 | -1.55E-12 | 1.33E-12 |  |
|  |  | THC | -1.17E-12 | 6.69E-13 | 0.091 | -2.53E-12 | 1.99E-13 |  |
|  |  | THC+CBD | -1.49600E-012^*^ | 6.86E-13 | 0.037 | -2.89E-12 | -9.78E-14 |  |
|  | THC | Vehicle | 1.06E-12 | 6.69E-13 | 0.124 | -3.06E-13 | 2.42E-12 |  |
|  |  | CBD | 1.17E-12 | 6.69E-13 | 0.091 | -1.99E-13 | 2.53E-12 |  |
|  |  | THC+CBD | -3.30E-13 | 6.48E-13 | 0.614 | -1.65E-12 | 9.92E-13 |  |
|  | THC+CBD | Vehicle | 1.39E-12 | 6.86E-13 | 0.051 | -9.40E-15 | 2.79E-12 |  |
|  |  | CBD | 1.49600E-012^*^ | 6.86E-13 | 0.037 | 9.78E-14 | 2.89E-12 |  |
|  |  | THC | 3.30E-13 | 6.48E-13 | 0.614 | -9.92E-13 | 1.65E-12 |  |
| *N-*linoleoyl glycine | Vehicle | CBD | 4.94E-14 | 3.14E-13 | 0.876 | -5.90E-13 | 6.89E-13 |  |
|  |  | THC | -5.78E-13 | 3.07E-13 | 0.069 | -1.20E-12 | 4.71E-14 | 1.54 |
|  |  | THC+CBD | -1.17149E-012^*^ | 3.14E-13 | 0.001 | -1.81E-12 | -5.32E-13 | 2.09 |
|  | CBD | Vehicle | -4.94E-14 | 3.14E-13 | 0.876 | -6.89E-13 | 5.90E-13 |  |
|  |  | THC | -6.27592E-013^*^ | 2.86E-13 | 0.036 | -1.21E-12 | -4.46E-14 |  |
|  |  | THC+CBD | -1.22089E-012^*^ | 2.93E-13 | 0.000 | -1.82E-12 | -6.23E-13 |  |
|  | THC | Vehicle | 5.78E-13 | 3.07E-13 | 0.069 | -4.71E-14 | 1.20E-12 |  |
|  |  | CBD | 6.27592E-013^*^ | 2.86E-13 | 0.036 | 4.46E-14 | 1.21E-12 |  |
|  |  | THC+CBD | -5.93296E-013^*^ | 2.86E-13 | 0.046 | -1.18E-12 | -1.03E-14 |  |
|  | THC+CBD | Vehicle | 1.17149E-012^*^ | 3.14E-13 | 0.001 | 5.32E-13 | 1.81E-12 |  |
|  |  | CBD | 1.22089E-012^*^ | 2.93E-13 | 0.000 | 6.23E-13 | 1.82E-12 |  |
|  |  | THC | 5.93296E-013^*^ | 2.86E-13 | 0.046 | 1.03E-14 | 1.18E-12 |  |
| *N-*stearoyl glycine | Vehicle | CBD | 3.55268E-013^*^ | 1.40E-13 | 0.016 | 6.99E-14 | 6.41E-13 | 0.59 |
|  |  | THC | 2.13E-13 | 1.33E-13 | 0.120 | -5.87E-14 | 4.85E-13 |  |
|  |  | THC+CBD | 3.15510E-013^*^ | 1.36E-13 | 0.027 | 3.77E-14 | 5.93E-13 | 0.64 |
|  | CBD | Vehicle | -3.55268E-013^*^ | 1.40E-13 | 0.016 | -6.41E-13 | -6.99E-14 |  |
|  |  | THC | -1.42E-13 | 1.28E-13 | 0.275 | -4.04E-13 | 1.19E-13 |  |
|  |  | THC+CBD | -3.98E-14 | 1.31E-13 | 0.764 | -3.08E-13 | 2.28E-13 |  |
|  | THC | Vehicle | -2.13E-13 | 1.33E-13 | 0.120 | -4.85E-13 | 5.87E-14 |  |
|  |  | CBD | 1.42E-13 | 1.28E-13 | 0.275 | -1.19E-13 | 4.04E-13 |  |
|  |  | THC+CBD | 1.02E-13 | 1.24E-13 | 0.415 | -1.51E-13 | 3.56E-13 |  |
|  | THC+CBD | Vehicle | -3.15510E-013^*^ | 1.36E-13 | 0.027 | -5.93E-13 | -3.77E-14 |  |
|  |  | CBD | 3.98E-14 | 1.31E-13 | 0.764 | -2.28E-13 | 3.08E-13 |  |
|  |  | THC | -1.02E-13 | 1.24E-13 | 0.415 | -3.56E-13 | 1.51E-13 |  |
| *N-*oleoyl glycine | Vehicle | CBD | -1.31E-13 | 1.83E-13 | 0.480 | -5.05E-13 | 2.42E-13 |  |
|  |  | THC | -2.06E-13 | 1.79E-13 | 0.258 | -5.71E-13 | 1.58E-13 |  |
|  |  | THC+CBD | -3.65E-13 | 1.83E-13 | 0.055 | -7.39E-13 | 8.55E-15 | 1.61 |
|  | CBD | Vehicle | 1.31E-13 | 1.83E-13 | 0.480 | -2.42E-13 | 5.05E-13 |  |
|  |  | THC | -7.52E-14 | 1.73E-13 | 0.668 | -4.29E-13 | 2.78E-13 |  |
|  |  | THC+CBD | -2.34E-13 | 1.78E-13 | 0.198 | -5.96E-13 | 1.29E-13 |  |
|  | THC | Vehicle | 2.06E-13 | 1.79E-13 | 0.258 | -1.58E-13 | 5.71E-13 |  |
|  |  | CBD | 7.52E-14 | 1.73E-13 | 0.668 | -2.78E-13 | 4.29E-13 |  |
|  |  | THC+CBD | -1.59E-13 | 1.73E-13 | 0.367 | -5.12E-13 | 1.95E-13 |  |
|  | THC+CBD | Vehicle | 3.65E-13 | 1.83E-13 | 0.055 | -8.55E-15 | 7.39E-13 |  |
|  |  | CBD | 2.34E-13 | 1.78E-13 | 0.198 | -1.29E-13 | 5.96E-13 |  |
|  |  | THC | 1.59E-13 | 1.73E-13 | 0.367 | -1.95E-13 | 5.12E-13 |  |

### Supplemental Figure 11. ANOVA for the effect of CBD, THC, or THC+CBD (3mg/kg) injections administered to a dam on *N-*acyl leucines measured in milk collected from nursing pups’ stomachs

|  | | Sum of Squares | df | Mean Square | F | Sig. |
| --- | --- | --- | --- | --- | --- | --- |
| *N-*palmitoyl leucine | Between Groups | 0.000 | 3 | 0.000 | 1.741 | 0.178 |
|  | Within Groups | 0.000 | 33 | 0.000 |  |  |
|  | Total | 0.000 | 36 |  |  |  |
| *N-*stearoyl leucine | Between Groups | 0.000 | 3 | 0.000 | 1.717 | 0.183 |
|  | Within Groups | 0.000 | 33 | 0.000 |  |  |
|  | Total | 0.000 | 36 |  |  |  |
| *N-*linoleoyl leucine | Between Groups | 0.000 | 3 | 0.000 | 2.287 | 0.098 |
|  | Within Groups | 0.000 | 31 | 0.000 |  |  |
|  | Total | 0.000 | 34 |  |  |  |
| *N-*oleoyl leucine | Between Groups | 0.000 | 3 | 0.000 | 1.682 | 0.190 |
|  | Within Groups | 0.000 | 33 | 0.000 |  |  |
|  | Total | 0.000 | 36 |  |  |  |
| *N-*docosahexaenoyl leucine | Between Groups | 0.000 | 3 | 0.000 | 6.643 | 0.001 |
|  | Within Groups | 0.000 | 31 | 0.000 |  |  |
|  | Total | 0.000 | 34 |  |  |  |

### Supplemental Figure 12. Post-hoc Fisher’s LSD results for *N-*acyl leucines

| Dependent Variable | | | Mean Difference (I-J) | Std. Error | Sig. | 95% Confidence Interval | |  |
| --- | --- | --- | --- | --- | --- | --- | --- | --- |
|  |  |  |  |  |  | Lower Bound | Upper Bound | Magnitude (drug/vehicle) |
| *N-*palmitoyl leucine | Vehicle | CBD | -3.14E-13 | 2.94E-13 | 0.294 | -9.13E-13 | 2.85E-13 |  |
|  |  | THC | -6.29443E-013^*^ | 2.81E-13 | 0.032 | -1.20E-12 | -5.69E-14 | 1.37 |
|  |  | THC+CBD | -2.50E-13 | 2.94E-13 | 0.402 | -8.49E-13 | 3.49E-13 |  |
|  | CBD | Vehicle | 3.14E-13 | 2.94E-13 | 0.294 | -2.85E-13 | 9.13E-13 |  |
|  |  | THC | -3.15E-13 | 2.72E-13 | 0.255 | -8.69E-13 | 2.38E-13 |  |
|  |  | THC+CBD | 6.40E-14 | 2.86E-13 | 0.824 | -5.17E-13 | 6.45E-13 |  |
|  | THC | Vehicle | 6.29443E-013^*^ | 2.81E-13 | 0.032 | 5.69E-14 | 1.20E-12 |  |
|  |  | CBD | 3.15E-13 | 2.72E-13 | 0.255 | -2.38E-13 | 8.69E-13 |  |
|  |  | THC+CBD | 3.79E-13 | 2.72E-13 | 0.173 | -1.74E-13 | 9.33E-13 |  |
|  | THC+CBD | Vehicle | 2.50E-13 | 2.94E-13 | 0.402 | -3.49E-13 | 8.49E-13 |  |
|  |  | CBD | -6.40E-14 | 2.86E-13 | 0.824 | -6.45E-13 | 5.17E-13 |  |
|  |  | THC | -3.79E-13 | 2.72E-13 | 0.173 | -9.33E-13 | 1.74E-13 |  |
| *N-*stearoyl leucine | Vehicle | CBD | 1.71E-13 | 9.86E-14 | 0.092 | -2.97E-14 | 3.72E-13 | 0.73 |
|  |  | THC | 1.47E-13 | 9.43E-14 | 0.128 | -4.47E-14 | 3.39E-13 |  |
|  |  | THC+CBD | 2.11928E-013^*^ | 9.86E-14 | 0.039 | 1.13E-14 | 4.13E-13 | 0.66 |
|  | CBD | Vehicle | -1.71E-13 | 9.86E-14 | 0.092 | -3.72E-13 | 2.97E-14 |  |
|  |  | THC | -2.38E-14 | 9.12E-14 | 0.796 | -2.09E-13 | 1.62E-13 |  |
|  |  | THC+CBD | 4.09E-14 | 9.57E-14 | 0.672 | -1.54E-13 | 2.36E-13 |  |
|  | THC | Vehicle | -1.47E-13 | 9.43E-14 | 0.128 | -3.39E-13 | 4.47E-14 |  |
|  |  | CBD | 2.38E-14 | 9.12E-14 | 0.796 | -1.62E-13 | 2.09E-13 |  |
|  |  | THC+CBD | 6.48E-14 | 9.12E-14 | 0.483 | -1.21E-13 | 2.50E-13 |  |
|  | THC+CBD | Vehicle | -2.11928E-013^*^ | 9.86E-14 | 0.039 | -4.13E-13 | -1.13E-14 |  |
|  |  | CBD | -4.09E-14 | 9.57E-14 | 0.672 | -2.36E-13 | 1.54E-13 |  |
|  |  | THC | -6.48E-14 | 9.12E-14 | 0.483 | -2.50E-13 | 1.21E-13 |  |
| *N-*linoleoyl leucine | Vehicle | CBD | -1.13E-13 | 6.91E-13 | 0.871 | -1.52E-12 | 1.30E-12 |  |
|  |  | THC | -1.46371E-012^*^ | 6.74E-13 | 0.038 | -2.84E-12 | -8.87E-14 | 1.46 |
|  |  | THC+CBD | -1.06E-12 | 7.11E-13 | 0.148 | -2.50E-12 | 3.94E-13 |  |
|  | CBD | Vehicle | 1.13E-13 | 6.91E-13 | 0.871 | -1.30E-12 | 1.52E-12 |  |
|  |  | THC | -1.35084E-012^*^ | 6.53E-13 | 0.047 | -2.68E-12 | -1.90E-14 |  |
|  |  | THC+CBD | -9.42E-13 | 6.91E-13 | 0.182 | -2.35E-12 | 4.66E-13 |  |
|  | THC | Vehicle | 1.46371E-012^*^ | 6.74E-13 | 0.038 | 8.87E-14 | 2.84E-12 |  |
|  |  | CBD | 1.35084E-012^*^ | 6.53E-13 | 0.047 | 1.90E-14 | 2.68E-12 |  |
|  |  | THC+CBD | 4.08E-13 | 6.74E-13 | 0.549 | -9.67E-13 | 1.78E-12 |  |
|  | THC+CBD | Vehicle | 1.06E-12 | 7.11E-13 | 0.148 | -3.94E-13 | 2.50E-12 |  |
|  |  | CBD | 9.42E-13 | 6.91E-13 | 0.182 | -4.66E-13 | 2.35E-12 |  |
|  |  | THC | -4.08E-13 | 6.74E-13 | 0.549 | -1.78E-12 | 9.67E-13 |  |
| *N-*oleoyl leucine | Vehicle | CBD | -2.05E-13 | 2.50E-13 | 0.420 | -7.14E-13 | 3.05E-13 |  |
|  |  | THC | -4.65E-13 | 2.39E-13 | 0.061 | -9.52E-13 | 2.19E-14 | 1.41 |
|  |  | THC+CBD | -4.62E-13 | 2.50E-13 | 0.074 | -9.72E-13 | 4.68E-14 | 1.40 |
|  | CBD | Vehicle | 2.05E-13 | 2.50E-13 | 0.420 | -3.05E-13 | 7.14E-13 |  |
|  |  | THC | -2.61E-13 | 2.32E-13 | 0.269 | -7.32E-13 | 2.11E-13 |  |
|  |  | THC+CBD | -2.58E-13 | 2.43E-13 | 0.296 | -7.52E-13 | 2.36E-13 |  |
|  | THC | Vehicle | 4.65E-13 | 2.39E-13 | 0.061 | -2.19E-14 | 9.52E-13 |  |
|  |  | CBD | 2.61E-13 | 2.32E-13 | 0.269 | -2.11E-13 | 7.32E-13 |  |
|  |  | THC+CBD | 2.67E-15 | 2.32E-13 | 0.991 | -4.68E-13 | 4.74E-13 |  |
|  | THC+CBD | Vehicle | 4.62E-13 | 2.50E-13 | 0.074 | -4.68E-14 | 9.72E-13 |  |
|  |  | CBD | 2.58E-13 | 2.43E-13 | 0.296 | -2.36E-13 | 7.52E-13 |  |
|  |  | THC | -2.67E-15 | 2.32E-13 | 0.991 | -4.74E-13 | 4.68E-13 |  |
| *N-*docosahexaenoyl leucine | Vehicle | CBD | 6.57957E-013^*^ | 2.27E-13 | 0.007 | 1.95E-13 | 1.12E-12 | 0.48 |
|  |  | THC | 4.86945E-013^*^ | 2.11E-13 | 0.028 | 5.68E-14 | 9.17E-13 | 0.62 |
|  |  | THC+CBD | 9.91831E-013^*^ | 2.27E-13 | 0.000 | 5.29E-13 | 1.45E-12 | 0.22 |
|  | CBD | Vehicle | -6.57957E-013^*^ | 2.27E-13 | 0.007 | -1.12E-12 | -1.95E-13 |  |
|  |  | THC | -1.71E-13 | 2.11E-13 | 0.424 | -6.01E-13 | 2.59E-13 |  |
|  |  | THC+CBD | 3.34E-13 | 2.27E-13 | 0.151 | -1.29E-13 | 7.97E-13 |  |
|  | THC | Vehicle | -4.86945E-013^*^ | 2.11E-13 | 0.028 | -9.17E-13 | -5.68E-14 |  |
|  |  | CBD | 1.71E-13 | 2.11E-13 | 0.424 | -2.59E-13 | 6.01E-13 |  |
|  |  | THC+CBD | 5.04886E-013^*^ | 2.11E-13 | 0.023 | 7.48E-14 | 9.35E-13 |  |
|  | THC+CBD | Vehicle | -9.91831E-013^*^ | 2.27E-13 | 0.000 | -1.45E-12 | -5.29E-13 |  |
|  |  | CBD | -3.34E-13 | 2.27E-13 | 0.151 | -7.97E-13 | 1.29E-13 |  |
|  |  | THC | -5.04886E-013^*^ | 2.11E-13 | 0.023 | -9.35E-13 | -7.48E-14 |  |

### Supplemental Figure 13. ANOVA for the effect of CBD, THC, or THC+CBD (3mg/kg) injections administered to a dam on *N-*acyl methionine levels measured in milk collected from nursing pups’ stomachs

|  | | Sum of Squares | df | Mean Square | F | Sig. |
| --- | --- | --- | --- | --- | --- | --- |
| *N-*palmitoyl methionine | Between Groups | 0.000 | 3 | 0.000 | 2.034 | 0.129 |
|  | Within Groups | 0.000 | 31 | 0.000 |  |  |
|  | Total | 0.000 | 34 |  |  |  |
| *N-*stearoyl methionine | Between Groups | 0.000 | 3 | 0.000 | 13.117 | 0.000 |
|  | Within Groups | 0.000 | 33 | 0.000 |  |  |
|  | Total | 0.000 | 36 |  |  |  |
| *N-*oleoyl methionine | Between Groups | 0.000 | 3 | 0.000 | 10.635 | 0.000 |
|  | Within Groups | 0.000 | 31 | 0.000 |  |  |
|  | Total | 0.000 | 34 |  |  |  |
| *N-*linoleoyl methionine | Between Groups | 0.000 | 3 | 0.000 | 41.032 | 0.000 |
|  | Within Groups | 0.000 | 32 | 0.000 |  |  |
|  | Total | 0.000 | 35 |  |  |  |
| *N-*arachidonoyl methionine | Between Groups | 0.000 | 3 | 0.000 | 33.997 | 0.000 |
|  | Within Groups | 0.000 | 32 | 0.000 |  |  |
|  | Total | 0.000 | 35 |  |  |  |

### Supplemental Figure 14. Post-hoc Fisher’s LSD results for *N-*acyl methionines

| Dependent Variable | | | Mean Difference (I-J) | Std. Error | Sig. | 95% Confidence Interval | |  |
| --- | --- | --- | --- | --- | --- | --- | --- | --- |
|  |  |  |  |  |  | Lower Bound | Upper Bound | Magnitude (drug/vehicle) |
| *N-*palmitoyl methionine | Vehicle | CBD | 6.73E-13 | 3.75E-13 | 0.083 | -9.26E-14 | 1.44E-12 | 0.74 |
|  |  | THC | 7.53659E-013^*^ | 3.49E-13 | 0.038 | 4.26E-14 | 1.46E-12 | 0.70 |
|  |  | THC+CBD | 7.88365E-013^*^ | 3.75E-13 | 0.044 | 2.32E-14 | 1.55E-12 | 0.69 |
|  | CBD | Vehicle | -6.73E-13 | 3.75E-13 | 0.083 | -1.44E-12 | 9.26E-14 |  |
|  |  | THC | 8.10E-14 | 3.49E-13 | 0.818 | -6.30E-13 | 7.92E-13 |  |
|  |  | THC+CBD | 1.16E-13 | 3.75E-13 | 0.760 | -6.49E-13 | 8.81E-13 |  |
|  | THC | Vehicle | -7.53659E-013^*^ | 3.49E-13 | 0.038 | -1.46E-12 | -4.26E-14 |  |
|  |  | CBD | -8.10E-14 | 3.49E-13 | 0.818 | -7.92E-13 | 6.30E-13 |  |
|  |  | THC+CBD | 3.47E-14 | 3.49E-13 | 0.921 | -6.76E-13 | 7.46E-13 |  |
|  | THC+CBD | Vehicle | -7.88365E-013^*^ | 3.75E-13 | 0.044 | -1.55E-12 | -2.32E-14 |  |
|  |  | CBD | -1.16E-13 | 3.75E-13 | 0.760 | -8.81E-13 | 6.49E-13 |  |
|  |  | THC | -3.47E-14 | 3.49E-13 | 0.921 | -7.46E-13 | 6.76E-13 |  |
| *N-*stearoyl methionine | Vehicle | CBD | 1.52956E-012^*^ | 2.77E-13 | 0.000 | 9.66E-13 | 2.09E-12 | 0.44 |
|  |  | THC | 1.36474E-012^*^ | 2.65E-13 | 0.000 | 8.26E-13 | 1.90E-12 | 0.50 |
|  |  | THC+CBD | 6.83600E-013^*^ | 2.77E-13 | 0.019 | 1.20E-13 | 1.25E-12 | 0.75 |
|  | CBD | Vehicle | -1.52956E-012^*^ | 2.77E-13 | 0.000 | -2.09E-12 | -9.66E-13 |  |
|  |  | THC | -1.65E-13 | 2.56E-13 | 0.524 | -6.86E-13 | 3.56E-13 |  |
|  |  | THC+CBD | -8.45961E-013^*^ | 2.69E-13 | 0.003 | -1.39E-12 | -3.00E-13 |  |
|  | THC | Vehicle | -1.36474E-012^*^ | 2.65E-13 | 0.000 | -1.90E-12 | -8.26E-13 |  |
|  |  | CBD | 1.65E-13 | 2.56E-13 | 0.524 | -3.56E-13 | 6.86E-13 |  |
|  |  | THC+CBD | -6.81145E-013^*^ | 2.56E-13 | 0.012 | -1.20E-12 | -1.60E-13 |  |
|  | THC+CBD | Vehicle | -6.83600E-013^*^ | 2.77E-13 | 0.019 | -1.25E-12 | -1.20E-13 |  |
|  |  | CBD | 8.45961E-013^*^ | 2.69E-13 | 0.003 | 3.00E-13 | 1.39E-12 |  |
|  |  | THC | 6.81145E-013^*^ | 2.56E-13 | 0.012 | 1.60E-13 | 1.20E-12 |  |
| *N-*oleoyl methionine | Vehicle | CBD | 1.30648E-012^*^ | 2.89E-13 | 0.000 | 7.18E-13 | 1.90E-12 | 0.59 |
|  |  | THC | 1.39435E-012^*^ | 2.70E-13 | 0.000 | 8.45E-13 | 1.94E-12 | 0.56 |
|  |  | THC+CBD | 7.30336E-013^*^ | 2.81E-13 | 0.014 | 1.57E-13 | 1.30E-12 | 0.77 |
|  | CBD | Vehicle | -1.30648E-012^*^ | 2.89E-13 | 0.000 | -1.90E-12 | -7.18E-13 |  |
|  |  | THC | 8.79E-14 | 2.59E-13 | 0.737 | -4.41E-13 | 6.16E-13 |  |
|  |  | THC+CBD | -5.76148E-013^*^ | 2.71E-13 | 0.042 | -1.13E-12 | -2.36E-14 |  |
|  | THC | Vehicle | -1.39435E-012^*^ | 2.70E-13 | 0.000 | -1.94E-12 | -8.45E-13 |  |
|  |  | CBD | -8.79E-14 | 2.59E-13 | 0.737 | -6.16E-13 | 4.41E-13 |  |
|  |  | THC+CBD | -6.64019E-013^*^ | 2.51E-13 | 0.013 | -1.18E-12 | -1.53E-13 |  |
|  | THC+CBD | Vehicle | -7.30336E-013^*^ | 2.81E-13 | 0.014 | -1.30E-12 | -1.57E-13 |  |
|  |  | CBD | 5.76148E-013^*^ | 2.71E-13 | 0.042 | 2.36E-14 | 1.13E-12 |  |
|  |  | THC | 6.64019E-013^*^ | 2.51E-13 | 0.013 | 1.53E-13 | 1.18E-12 |  |
| *N-*linoleoyl methionine | Vehicle | CBD | 2.17994E-012^*^ | 5.49E-13 | 0.000 | 1.06E-12 | 3.30E-12 | 0.64 |
|  |  | THC | 8.45E-13 | 5.10E-13 | 0.107 | -1.94E-13 | 1.88E-12 |  |
|  |  | THC+CBD | -3.37998E-012^*^ | 5.33E-13 | 0.000 | -4.47E-12 | -2.29E-12 | 1.56 |
|  | CBD | Vehicle | -2.17994E-012^*^ | 5.49E-13 | 0.000 | -3.30E-12 | -1.06E-12 |  |
|  |  | THC | -1.33498E-012^*^ | 5.10E-13 | 0.013 | -2.37E-12 | -2.96E-13 |  |
|  |  | THC+CBD | -5.55992E-012^*^ | 5.33E-13 | 0.000 | -6.65E-12 | -4.47E-12 |  |
|  | THC | Vehicle | -8.45E-13 | 5.10E-13 | 0.107 | -1.88E-12 | 1.94E-13 |  |
|  |  | CBD | 1.33498E-012^*^ | 5.10E-13 | 0.013 | 2.96E-13 | 2.37E-12 |  |
|  |  | THC+CBD | -4.22494E-012^*^ | 4.93E-13 | 0.000 | -5.23E-12 | -3.22E-12 |  |
|  | THC+CBD | Vehicle | 3.37998E-012^*^ | 5.33E-13 | 0.000 | 2.29E-12 | 4.47E-12 |  |
|  |  | CBD | 5.55992E-012^*^ | 5.33E-13 | 0.000 | 4.47E-12 | 6.65E-12 |  |
|  |  | THC | 4.22494E-012^*^ | 4.93E-13 | 0.000 | 3.22E-12 | 5.23E-12 |  |
| *N-*arachidonoyl methionine | Vehicle | CBD | 9.94023E-013^*^ | 2.53E-13 | 0.000 | 4.78E-13 | 1.51E-12 | 0.57 |
|  |  | THC | 1.01865E-012^*^ | 2.47E-13 | 0.000 | 5.15E-13 | 1.52E-12 | 0.55 |
|  |  | THC+CBD | -1.09556E-012^*^ | 2.53E-13 | 0.000 | -1.61E-12 | -5.80E-13 | 1.48 |
|  | CBD | Vehicle | -9.94023E-013^*^ | 2.53E-13 | 0.000 | -1.51E-12 | -4.78E-13 |  |
|  |  | THC | 2.46E-14 | 2.39E-13 | 0.919 | -4.63E-13 | 5.12E-13 |  |
|  |  | THC+CBD | -2.08959E-012^*^ | 2.46E-13 | 0.000 | -2.59E-12 | -1.59E-12 |  |
|  | THC | Vehicle | -1.01865E-012^*^ | 2.47E-13 | 0.000 | -1.52E-12 | -5.15E-13 |  |
|  |  | CBD | -2.46E-14 | 2.39E-13 | 0.919 | -5.12E-13 | 4.63E-13 |  |
|  |  | THC+CBD | -2.11421E-012^*^ | 2.39E-13 | 0.000 | -2.60E-12 | -1.63E-12 |  |
|  | THC+CBD | Vehicle | 1.09556E-012^*^ | 2.53E-13 | 0.000 | 5.80E-13 | 1.61E-12 |  |
|  |  | CBD | 2.08959E-012^*^ | 2.46E-13 | 0.000 | 1.59E-12 | 2.59E-12 |  |
|  |  | THC | 2.11421E-012^*^ | 2.39E-13 | 0.000 | 1.63E-12 | 2.60E-12 |  |

### Supplemental Figure 15. ANOVA for the effect of CBD, THC, or THC+CBD (3mg/kg) injections administered to a dam on *N-*acyl phenylalanines measured in milk collected from nursing pups’ stomachs

|  | | Sum of Squares | df | Mean Square | F | Sig. |
| --- | --- | --- | --- | --- | --- | --- |
| *N-*palmitoyl phenylalanine | Between Groups | 0.000 | 3 | 0.000 | 1.052 | 0.384 |
|  | Within Groups | 0.000 | 30 | 0.000 |  |  |
|  | Total | 0.000 | 33 |  |  |  |
| *N-*stearoyl phenylalanine | Between Groups | 0.000 | 3 | 0.000 | 8.939 | 0.000 |
|  | Within Groups | 0.000 | 32 | 0.000 |  |  |
|  | Total | 0.000 | 35 |  |  |  |
| *N-*oleoyl phenylalanine | Between Groups | 0.000 | 3 | 0.000 | 1.386 | 0.265 |
|  | Within Groups | 0.000 | 32 | 0.000 |  |  |
|  | Total | 0.000 | 35 |  |  |  |
| *N-*linoleoyl phenylalanine | Between Groups | 0.000 | 3 | 0.000 | 2.597 | 0.069 |
|  | Within Groups | 0.000 | 32 | 0.000 |  |  |
|  | Total | 0.000 | 35 |  |  |  |
| *N-*arachidonoyl phenylalanine | Between Groups | 0.000 | 3 | 0.000 | 0.468 | 0.709 |
|  | Within Groups | 0.000 | 14 | 0.000 |  |  |
|  | Total | 0.000 | 17 |  |  |  |

### Supplemental Figure 16. Post-hoc Fisher’s LSD results for *N-*acyl phenylalanines

| Dependent Variable | | | Mean Difference (I-J) | Std. Error | Sig. | 95% Confidence Interval | |  |
| --- | --- | --- | --- | --- | --- | --- | --- | --- |
|  |  |  |  |  |  | Lower Bound | Upper Bound | Magnitude (drug/vehicle) |
| *N-*palmitoyl phenylalanine | Vehicle | CBD | -3.56E-14 | 1.17E-13 | 0.763 | -2.75E-13 | 2.04E-13 |  |
|  |  | THC | -1.35E-13 | 1.12E-13 | 0.236 | -3.63E-13 | 9.30E-14 |  |
|  |  | THC+CBD | 4.21E-14 | 1.14E-13 | 0.715 | -1.91E-13 | 2.75E-13 |  |
|  | CBD | Vehicle | 3.56E-14 | 1.17E-13 | 0.763 | -2.04E-13 | 2.75E-13 |  |
|  |  | THC | -9.94E-14 | 1.07E-13 | 0.363 | -3.19E-13 | 1.20E-13 |  |
|  |  | THC+CBD | 7.78E-14 | 1.10E-13 | 0.485 | -1.47E-13 | 3.03E-13 |  |
|  | THC | Vehicle | 1.35E-13 | 1.12E-13 | 0.236 | -9.30E-14 | 3.63E-13 |  |
|  |  | CBD | 9.94E-14 | 1.07E-13 | 0.363 | -1.20E-13 | 3.19E-13 |  |
|  |  | THC+CBD | 1.77E-13 | 1.04E-13 | 0.099 | -3.55E-14 | 3.90E-13 |  |
|  | THC+CBD | Vehicle | -4.21E-14 | 1.14E-13 | 0.715 | -2.75E-13 | 1.91E-13 |  |
|  |  | CBD | -7.78E-14 | 1.10E-13 | 0.485 | -3.03E-13 | 1.47E-13 |  |
|  |  | THC | -1.77E-13 | 1.04E-13 | 0.099 | -3.90E-13 | 3.55E-14 |  |
| *N-*stearoyl phenylalanine | Vehicle | CBD | 2.43E-14 | 3.44E-14 | 0.485 | -4.58E-14 | 9.45E-14 |  |
|  |  | THC | -5.41E-14 | 3.29E-14 | 0.110 | -1.21E-13 | 1.29E-14 |  |
|  |  | THC+CBD | 1.14626E-013^*^ | 3.54E-14 | 0.003 | 4.25E-14 | 1.87E-13 | 0.61 |
|  | CBD | Vehicle | -2.43E-14 | 3.44E-14 | 0.485 | -9.45E-14 | 4.58E-14 |  |
|  |  | THC | -7.84865E-014^*^ | 3.18E-14 | 0.019 | -1.43E-13 | -1.36E-14 |  |
|  |  | THC+CBD | 9.02838E-014^*^ | 3.44E-14 | 0.013 | 2.02E-14 | 1.60E-13 |  |
|  | THC | Vehicle | 5.41E-14 | 3.29E-14 | 0.110 | -1.29E-14 | 1.21E-13 |  |
|  |  | CBD | 7.84865E-014^*^ | 3.18E-14 | 0.019 | 1.36E-14 | 1.43E-13 |  |
|  |  | THC+CBD | 1.68770E-013^*^ | 3.29E-14 | 0.000 | 1.02E-13 | 2.36E-13 |  |
|  | THC+CBD | Vehicle | -1.14626E-013^*^ | 3.54E-14 | 0.003 | -1.87E-13 | -4.25E-14 |  |
|  |  | CBD | -9.02838E-014^*^ | 3.44E-14 | 0.013 | -1.60E-13 | -2.02E-14 |  |
|  |  | THC | -1.68770E-013^*^ | 3.29E-14 | 0.000 | -2.36E-13 | -1.02E-13 |  |
| *N-*oleoyl phenylalanine | Vehicle | CBD | -8.42E-14 | 1.02E-13 | 0.417 | -2.93E-13 | 1.24E-13 |  |
|  |  | THC | -1.76E-13 | 9.51E-14 | 0.074 | -3.69E-13 | 1.82E-14 | 1.29 |
|  |  | THC+CBD | -1.63E-13 | 9.95E-14 | 0.111 | -3.65E-13 | 3.97E-14 |  |
|  | CBD | Vehicle | 8.42E-14 | 1.02E-13 | 0.417 | -1.24E-13 | 2.93E-13 |  |
|  |  | THC | -9.14E-14 | 9.51E-14 | 0.344 | -2.85E-13 | 1.02E-13 |  |
|  |  | THC+CBD | -7.87E-14 | 9.95E-14 | 0.435 | -2.81E-13 | 1.24E-13 |  |
|  | THC | Vehicle | 1.76E-13 | 9.51E-14 | 0.074 | -1.82E-14 | 3.69E-13 |  |
|  |  | CBD | 9.14E-14 | 9.51E-14 | 0.344 | -1.02E-13 | 2.85E-13 |  |
|  |  | THC+CBD | 1.27E-14 | 9.20E-14 | 0.891 | -1.75E-13 | 2.00E-13 |  |
|  | THC+CBD | Vehicle | 1.63E-13 | 9.95E-14 | 0.111 | -3.97E-14 | 3.65E-13 |  |
|  |  | CBD | 7.87E-14 | 9.95E-14 | 0.435 | -1.24E-13 | 2.81E-13 |  |
|  |  | THC | -1.27E-14 | 9.20E-14 | 0.891 | -2.00E-13 | 1.75E-13 |  |
| *N-*linoleoyl phenylalanine | Vehicle | CBD | -2.78E-13 | 2.27E-13 | 0.229 | -7.40E-13 | 1.84E-13 |  |
|  |  | THC | -3.89E-13 | 2.21E-13 | 0.089 | -8.40E-13 | 6.23E-14 | 1.42 |
|  |  | THC+CBD | -6.22053E-013^*^ | 2.27E-13 | 0.010 | -1.08E-12 | -1.60E-13 | 1.68 |
|  | CBD | Vehicle | 2.78E-13 | 2.27E-13 | 0.229 | -1.84E-13 | 7.40E-13 |  |
|  |  | THC | -1.11E-13 | 2.14E-13 | 0.610 | -5.47E-13 | 3.26E-13 |  |
|  |  | THC+CBD | -3.44E-13 | 2.20E-13 | 0.128 | -7.92E-13 | 1.04E-13 |  |
|  | THC | Vehicle | 3.89E-13 | 2.21E-13 | 0.089 | -6.23E-14 | 8.40E-13 |  |
|  |  | CBD | 1.11E-13 | 2.14E-13 | 0.610 | -3.26E-13 | 5.47E-13 |  |
|  |  | THC+CBD | -2.33E-13 | 2.14E-13 | 0.284 | -6.70E-13 | 2.03E-13 |  |
|  | THC+CBD | Vehicle | 6.22053E-013^*^ | 2.27E-13 | 0.010 | 1.60E-13 | 1.08E-12 |  |
|  |  | CBD | 3.44E-13 | 2.20E-13 | 0.128 | -1.04E-13 | 7.92E-13 |  |
|  |  | THC | 2.33E-13 | 2.14E-13 | 0.284 | -2.03E-13 | 6.70E-13 |  |
| *N-*arachidonoyl phenylalanine | Vehicle | CBD | 2.85E-14 | 3.52E-14 | 0.432 | -4.71E-14 | 1.04E-13 |  |
|  |  | THC | -7.20E-15 | 4.25E-14 | 0.868 | -9.84E-14 | 8.40E-14 |  |
|  |  | THC+CBD | 3.13E-14 | 3.90E-14 | 0.436 | -5.24E-14 | 1.15E-13 |  |
|  | CBD | Vehicle | -2.85E-14 | 3.52E-14 | 0.432 | -1.04E-13 | 4.71E-14 |  |
|  |  | THC | -3.57E-14 | 4.12E-14 | 0.400 | -1.24E-13 | 5.26E-14 |  |
|  |  | THC+CBD | 2.79E-15 | 3.76E-14 | 0.942 | -7.78E-14 | 8.34E-14 |  |
|  | THC | Vehicle | 7.20E-15 | 4.25E-14 | 0.868 | -8.40E-14 | 9.84E-14 |  |
|  |  | CBD | 3.57E-14 | 4.12E-14 | 0.400 | -5.26E-14 | 1.24E-13 |  |
|  |  | THC+CBD | 3.85E-14 | 4.45E-14 | 0.401 | -5.68E-14 | 1.34E-13 |  |
|  | THC+CBD | Vehicle | -3.13E-14 | 3.90E-14 | 0.436 | -1.15E-13 | 5.24E-14 |  |
|  |  | CBD | -2.79E-15 | 3.76E-14 | 0.942 | -8.34E-14 | 7.78E-14 |  |
|  |  | THC | -3.85E-14 | 4.45E-14 | 0.401 | -1.34E-13 | 5.68E-14 |  |

### Supplemental Figure 17. ANOVA for the effect of CBD, THC, or THC+CBD (3mg/kg) injections administered to a dam on *N-*acyl prolines measured in milk collected from nursing pups’ stomachs

|  | | Sum of Squares | df | Mean Square | F | Sig. |
| --- | --- | --- | --- | --- | --- | --- |
| *N-*palmitoyl proline | Between Groups | 0.000 | 3 | 0.000 | 3.903 | 0.018 |
|  | Within Groups | 0.000 | 32 | 0.000 |  |  |
|  | Total | 0.000 | 35 |  |  |  |
|  | Total | 0.000 | 2 |  |  |  |
| *N-*linoleoyl proline | Between Groups | 0.000 | 3 | 0.000 | 7.441 | 0.001 |
|  | Within Groups | 0.000 | 27 | 0.000 |  |  |
|  | Total | 0.000 | 30 |  |  |  |

### Supplemental Figure 18. Post-hoc Fisher’s LSD results for *N-*acyl prolines

| Dependent Variable | | | Mean Difference (I-J) | Std. Error | Sig. | 95% Confidence Interval | |  |
| --- | --- | --- | --- | --- | --- | --- | --- | --- |
|  |  |  |  |  |  | Lower Bound | Upper Bound | Magnitude (drug/vehicle) |
| *N-*palmitoyl proline | Vehicle | CBD | -3.29E-14 | 1.81E-14 | 0.079 | -6.98E-14 | 4.03E-15 | 1.47 |
|  |  | THC | -4.06232E-014^*^ | 1.73E-14 | 0.025 | -7.59E-14 | -5.31E-15 | 1.58 |
|  |  | THC+CBD | -6.26544E-014^*^ | 1.87E-14 | 0.002 | -1.01E-13 | -2.47E-14 | 1.90 |
|  | CBD | Vehicle | 3.29E-14 | 1.81E-14 | 0.079 | -4.03E-15 | 6.98E-14 |  |
|  |  | THC | -7.72E-15 | 1.68E-14 | 0.648 | -4.19E-14 | 2.64E-14 |  |
|  |  | THC+CBD | -2.98E-14 | 1.81E-14 | 0.111 | -6.67E-14 | 7.18E-15 |  |
|  | THC | Vehicle | 4.06232E-014^*^ | 1.73E-14 | 0.025 | 5.31E-15 | 7.59E-14 |  |
|  |  | CBD | 7.72E-15 | 1.68E-14 | 0.648 | -2.64E-14 | 4.19E-14 |  |
|  |  | THC+CBD | -2.20E-14 | 1.73E-14 | 0.213 | -5.73E-14 | 1.33E-14 |  |
|  | THC+CBD | Vehicle | 6.26544E-014^*^ | 1.87E-14 | 0.002 | 2.47E-14 | 1.01E-13 |  |
|  |  | CBD | 2.98E-14 | 1.81E-14 | 0.111 | -7.18E-15 | 6.67E-14 |  |
|  |  | THC | 2.20E-14 | 1.73E-14 | 0.213 | -1.33E-14 | 5.73E-14 |  |
| *N-*linoleoyl proline | Vehicle | CBD | -6.80E-15 | 8.34E-15 | 0.422 | -2.39E-14 | 1.03E-14 |  |
|  |  | THC | -1.18E-14 | 7.77E-15 | 0.139 | -2.78E-14 | 4.08E-15 |  |
|  |  | THC+CBD | -3.15646E-014^*^ | 7.99E-15 | 0.001 | -4.80E-14 | -1.52E-14 | 2.52 |
|  | CBD | Vehicle | 6.80E-15 | 8.34E-15 | 0.422 | -1.03E-14 | 2.39E-14 |  |
|  |  | THC | -5.05E-15 | 6.43E-15 | 0.439 | -1.82E-14 | 8.15E-15 |  |
|  |  | THC+CBD | -2.47623E-014^*^ | 6.70E-15 | 0.001 | -3.85E-14 | -1.10E-14 |  |
|  | THC | Vehicle | 1.18E-14 | 7.77E-15 | 0.139 | -4.08E-15 | 2.78E-14 |  |
|  |  | CBD | 5.05E-15 | 6.43E-15 | 0.439 | -8.15E-15 | 1.82E-14 |  |
|  |  | THC+CBD | -1.97148E-014^*^ | 5.98E-15 | 0.003 | -3.20E-14 | -7.45E-15 |  |
|  | THC+CBD | Vehicle | 3.15646E-014^*^ | 7.99E-15 | 0.001 | 1.52E-14 | 4.80E-14 |  |
|  |  | CBD | 2.47623E-014^*^ | 6.70E-15 | 0.001 | 1.10E-14 | 3.85E-14 |  |
|  |  | THC | 1.97148E-014^*^ | 5.98E-15 | 0.003 | 7.45E-15 | 3.20E-14 |  |

### Supplemental Figure 19. ANOVA for the effect of CBD, THC, or THC+CBD (3mg/kg) injections administered to a dam on *N-*acyl serines measured in milk collected from nursing pups’ stomachs

|  | | Sum of Squares | df | Mean Square | F | Sig. |
| --- | --- | --- | --- | --- | --- | --- |
| *N-*palmitoyl serine | Between Groups | 0.000 | 3 | 0.000 | 7.053 | 0.001 |
|  | Within Groups | 0.000 | 33 | 0.000 |  |  |
|  | Total | 0.000 | 36 |  |  |  |
| *N-*steroyl serine | Between Groups | 0.000 | 3 | 0.000 | 5.653 | 0.003 |
|  | Within Groups | 0.000 | 30 | 0.000 |  |  |
|  | Total | 0.000 | 33 |  |  |  |
| *N-*oleoyl serine | Between Groups | 0.000 | 3 | 0.000 | 8.910 | 0.000 |
|  | Within Groups | 0.000 | 32 | 0.000 |  |  |
|  | Total | 0.000 | 35 |  |  |  |
| *N-*linoleoyl serine | Between Groups | 0.000 | 3 | 0.000 | 7.586 | 0.001 |
|  | Within Groups | 0.000 | 32 | 0.000 |  |  |
|  | Total | 0.000 | 35 |  |  |  |
| *N-*arachidonoyl serine | Between Groups | 0.000 | 3 | 0.000 | 3.990 | 0.042 |
|  | Within Groups | 0.000 | 10 | 0.000 |  |  |
|  | Total | 0.000 | 13 |  |  |  |

### Supplemental Figure 20. Post-hoc Fisher’s LSD results for *N-*acyl serines

| Dependent Variable | | | Mean Difference (I-J) | Std. Error | Sig. | 95% Confidence Interval | |  |
| --- | --- | --- | --- | --- | --- | --- | --- | --- |
|  |  |  |  |  |  | Lower Bound | Upper Bound | Magnitude (drug/vehicle) |
| *N-*palmitoyl serine | Vehicle | CBD | -3.61E-14 | 6.68E-13 | 0.957 | -1.40E-12 | 1.32E-12 |  |
|  |  | THC | -2.39482E-012^*^ | 6.39E-13 | 0.001 | -3.69E-12 | -1.09E-12 | 1.66 |
|  |  | THC+CBD | -3.70E-13 | 6.68E-13 | 0.583 | -1.73E-12 | 9.89E-13 |  |
|  | CBD | Vehicle | 3.61E-14 | 6.68E-13 | 0.957 | -1.32E-12 | 1.40E-12 |  |
|  |  | THC | -2.35875E-012^*^ | 6.18E-13 | 0.001 | -3.62E-12 | -1.10E-12 |  |
|  |  | THC+CBD | -3.34E-13 | 6.48E-13 | 0.610 | -1.65E-12 | 9.85E-13 |  |
|  | THC | Vehicle | 2.39482E-012^*^ | 6.39E-13 | 0.001 | 1.09E-12 | 3.69E-12 |  |
|  |  | CBD | 2.35875E-012^*^ | 6.18E-13 | 0.001 | 1.10E-12 | 3.62E-12 |  |
|  |  | THC+CBD | 2.02473E-012^*^ | 6.18E-13 | 0.002 | 7.67E-13 | 3.28E-12 |  |
|  | THC+CBD | Vehicle | 3.70E-13 | 6.68E-13 | 0.583 | -9.89E-13 | 1.73E-12 |  |
|  |  | CBD | 3.34E-13 | 6.48E-13 | 0.610 | -9.85E-13 | 1.65E-12 |  |
|  |  | THC | -2.02473E-012^*^ | 6.18E-13 | 0.002 | -3.28E-12 | -7.67E-13 |  |
| *N-*stearoyl serine | Vehicle | CBD | 1.43E-13 | 1.06E-13 | 0.187 | -7.31E-14 | 3.59E-13 |  |
|  |  | THC | 1.90E-13 | 1.04E-13 | 0.077 | -2.17E-14 | 4.01E-13 | 0.73 |
|  |  | THC+CBD | 4.34124E-013^*^ | 1.09E-13 | 0.000 | 2.12E-13 | 6.56E-13 | 0.38 |
|  | CBD | Vehicle | -1.43E-13 | 1.06E-13 | 0.187 | -3.59E-13 | 7.31E-14 |  |
|  |  | THC | 4.67E-14 | 9.65E-14 | 0.632 | -1.50E-13 | 2.44E-13 |  |
|  |  | THC+CBD | 2.91084E-013^*^ | 1.02E-13 | 0.008 | 8.26E-14 | 5.00E-13 |  |
|  | THC | Vehicle | -1.90E-13 | 1.04E-13 | 0.077 | -4.01E-13 | 2.17E-14 |  |
|  |  | CBD | -4.67E-14 | 9.65E-14 | 0.632 | -2.44E-13 | 1.50E-13 |  |
|  |  | THC+CBD | 2.44430E-013^*^ | 9.96E-14 | 0.020 | 4.10E-14 | 4.48E-13 |  |
|  | THC+CBD | Vehicle | -4.34124E-013^*^ | 1.09E-13 | 0.000 | -6.56E-13 | -2.12E-13 |  |
|  |  | CBD | -2.91084E-013^*^ | 1.02E-13 | 0.008 | -5.00E-13 | -8.26E-14 |  |
|  |  | THC | -2.44430E-013^*^ | 9.96E-14 | 0.020 | -4.48E-13 | -4.10E-14 |  |
| *N-*oleoyl serine | Vehicle | CBD | -1.53E-13 | 2.17E-13 | 0.488 | -5.95E-13 | 2.90E-13 |  |
|  |  | THC | -8.77331E-013^*^ | 2.02E-13 | 0.000 | -1.29E-12 | -4.66E-13 | 2.03 |
|  |  | THC+CBD | -7.41931E-013^*^ | 2.11E-13 | 0.001 | -1.17E-12 | -3.12E-13 | 1.87 |
|  | CBD | Vehicle | 1.53E-13 | 2.17E-13 | 0.488 | -2.90E-13 | 5.95E-13 |  |
|  |  | THC | -7.24670E-013^*^ | 2.02E-13 | 0.001 | -1.14E-12 | -3.13E-13 |  |
|  |  | THC+CBD | -5.89270E-013^*^ | 2.11E-13 | 0.009 | -1.02E-12 | -1.59E-13 |  |
|  | THC | Vehicle | 8.77331E-013^*^ | 2.02E-13 | 0.000 | 4.66E-13 | 1.29E-12 |  |
|  |  | CBD | 7.24670E-013^*^ | 2.02E-13 | 0.001 | 3.13E-13 | 1.14E-12 |  |
|  |  | THC+CBD | 1.35E-13 | 1.95E-13 | 0.493 | -2.63E-13 | 5.33E-13 |  |
|  | THC+CBD | Vehicle | 7.41931E-013^*^ | 2.11E-13 | 0.001 | 3.12E-13 | 1.17E-12 |  |
|  |  | CBD | 5.89270E-013^*^ | 2.11E-13 | 0.009 | 1.59E-13 | 1.02E-12 |  |
|  |  | THC | -1.35E-13 | 1.95E-13 | 0.493 | -5.33E-13 | 2.63E-13 |  |
| *N-*linoleoyl serine | Vehicle | CBD | -1.20E-13 | 5.24E-13 | 0.821 | -1.19E-12 | 9.48E-13 |  |
|  |  | THC | -1.83404E-012^*^ | 4.87E-13 | 0.001 | -2.83E-12 | -8.41E-13 | 1.64 |
|  |  | THC+CBD | -1.59546E-012^*^ | 5.10E-13 | 0.004 | -2.63E-12 | -5.57E-13 | 1.56 |
|  | CBD | Vehicle | 1.20E-13 | 5.24E-13 | 0.821 | -9.48E-13 | 1.19E-12 |  |
|  |  | THC | -1.71426E-012^*^ | 4.87E-13 | 0.001 | -2.71E-12 | -7.22E-13 |  |
|  |  | THC+CBD | -1.47569E-012^*^ | 5.10E-13 | 0.007 | -2.51E-12 | -4.38E-13 |  |
|  | THC | Vehicle | 1.83404E-012^*^ | 4.87E-13 | 0.001 | 8.41E-13 | 2.83E-12 |  |
|  |  | CBD | 1.71426E-012^*^ | 4.87E-13 | 0.001 | 7.22E-13 | 2.71E-12 |  |
|  |  | THC+CBD | 2.39E-13 | 4.71E-13 | 0.616 | -7.22E-13 | 1.20E-12 |  |
|  | THC+CBD | Vehicle | 1.59546E-012^*^ | 5.10E-13 | 0.004 | 5.57E-13 | 2.63E-12 |  |
|  |  | CBD | 1.47569E-012^*^ | 5.10E-13 | 0.007 | 4.38E-13 | 2.51E-12 |  |
|  |  | THC | -2.39E-13 | 4.71E-13 | 0.616 | -1.20E-12 | 7.22E-13 |  |
| *N-*arachidonoyl serine | Vehicle | CBD | 4.41E-14 | 2.80E-14 | 0.146 | -1.83E-14 | 1.07E-13 |  |
|  |  | THC | 8.36245E-014^*^ | 2.62E-14 | 0.010 | 2.52E-14 | 1.42E-13 | 0.38 |
|  |  | THC+CBD | 7.36229E-014^*^ | 2.62E-14 | 0.018 | 1.52E-14 | 1.32E-13 | 0.45 |
|  | CBD | Vehicle | -4.41E-14 | 2.80E-14 | 0.146 | -1.07E-13 | 1.83E-14 |  |
|  |  | THC | 3.95E-14 | 2.62E-14 | 0.162 | -1.89E-14 | 9.79E-14 |  |
|  |  | THC+CBD | 2.95E-14 | 2.62E-14 | 0.286 | -2.89E-14 | 8.79E-14 |  |
|  | THC | Vehicle | -8.36245E-014^*^ | 2.62E-14 | 0.010 | -1.42E-13 | -2.52E-14 |  |
|  |  | CBD | -3.95E-14 | 2.62E-14 | 0.162 | -9.79E-14 | 1.89E-14 |  |
|  |  | THC+CBD | -1.00E-14 | 2.43E-14 | 0.689 | -6.41E-14 | 4.41E-14 |  |
|  | THC+CBD | Vehicle | -7.36229E-014^*^ | 2.62E-14 | 0.018 | -1.32E-13 | -1.52E-14 |  |
|  |  | CBD | -2.95E-14 | 2.62E-14 | 0.286 | -8.79E-14 | 2.89E-14 |  |
|  |  | THC | 1.00E-14 | 2.43E-14 | 0.689 | -4.41E-14 | 6.41E-14 |  |

### Supplemental Figure 21. ANOVA for the effect of CBD, THC, or THC+CBD (3mg/kg) injections administered to a dam on *N-*acyl taurine levels measured in milk collected from nursing pups’ stomachs

|  | | Sum of Squares | df | Mean Square | F | Sig. |
| --- | --- | --- | --- | --- | --- | --- |
| *N-*palmitoyl taurine | Between Groups | 0.000 | 3 | 0.000 | 2.867 | 0.054 |
|  | Within Groups | 0.000 | 29 | 0.000 |  |  |
|  | Total | 0.000 | 32 |  |  |  |
| *N-*oleoyl taurine | Between Groups | 0.000 | 3 | 0.000 | 4.500 | 0.010 |
|  | Within Groups | 0.000 | 30 | 0.000 |  |  |
|  | Total | 0.000 | 33 |  |  |  |
| *N-*arachidonoyl taurine | Between Groups | 0.000 | 3 | 0.000 | 20.885 | 0.000 |
|  | Within Groups | 0.000 | 30 | 0.000 |  |  |
|  | Total | 0.000 | 33 |  |  |  |

### Supplemental Figure 22. Post-hoc Fisher’s LSD results for *N-*acyl taurines

| Dependent Variable | | | Mean Difference (I-J) | Std. Error | Sig. | 95% Confidence Interval | |  |
| --- | --- | --- | --- | --- | --- | --- | --- | --- |
|  |  |  |  |  |  | Lower Bound | Upper Bound | Magnitude (drug/vehicle) |
| *N-*palmitoyl taurine | Vehicle | CBD | 3.61663E-011^*^ | 1.37E-11 | 0.013 | 8.08E-12 | 6.43E-11 | 0.37 |
|  |  | THC | 2.44E-11 | 1.27E-11 | 0.063 | -1.45E-12 | 5.03E-11 | 0.58 |
|  |  | THC+CBD | 9.25E-12 | 1.29E-11 | 0.481 | -1.72E-11 | 3.57E-11 |  |
|  | CBD | Vehicle | -3.61663E-011^*^ | 1.37E-11 | 0.013 | -6.43E-11 | -8.08E-12 |  |
|  |  | THC | -1.17E-11 | 1.27E-11 | 0.362 | -3.76E-11 | 1.42E-11 |  |
|  |  | THC+CBD | -2.69198E-011^*^ | 1.29E-11 | 0.047 | -5.34E-11 | -4.41E-13 |  |
|  | THC | Vehicle | -2.44E-11 | 1.27E-11 | 0.063 | -5.03E-11 | 1.45E-12 |  |
|  |  | CBD | 1.17E-11 | 1.27E-11 | 0.362 | -1.42E-11 | 3.76E-11 |  |
|  |  | THC+CBD | -1.52E-11 | 1.18E-11 | 0.208 | -3.93E-11 | 8.95E-12 |  |
|  | THC+CBD | Vehicle | -9.25E-12 | 1.29E-11 | 0.481 | -3.57E-11 | 1.72E-11 |  |
|  |  | CBD | 2.69198E-011^*^ | 1.29E-11 | 0.047 | 4.41E-13 | 5.34E-11 |  |
|  |  | THC | 1.52E-11 | 1.18E-11 | 0.208 | -8.95E-12 | 3.93E-11 |  |
| *N-*oleoyl taurine | Vehicle | CBD | 1.94779E-011^*^ | 5.75E-12 | 0.002 | 7.74E-12 | 3.12E-11 | 3.31 |
|  |  | THC | 7.98E-12 | 5.45E-12 | 0.154 | -3.16E-12 | 1.91E-11 |  |
|  |  | THC+CBD | 2.72E-12 | 5.75E-12 | 0.639 | -9.02E-12 | 1.45E-11 |  |
|  | CBD | Vehicle | -1.94779E-011^*^ | 5.75E-12 | 0.002 | -3.12E-11 | -7.74E-12 |  |
|  |  | THC | -1.15005E-011^*^ | 5.45E-12 | 0.043 | -2.26E-11 | -3.68E-13 |  |
|  |  | THC+CBD | -1.67583E-011^*^ | 5.75E-12 | 0.007 | -2.85E-11 | -5.02E-12 |  |
|  | THC | Vehicle | -7.98E-12 | 5.45E-12 | 0.154 | -1.91E-11 | 3.16E-12 |  |
|  |  | CBD | 1.15005E-011^*^ | 5.45E-12 | 0.043 | 3.68E-13 | 2.26E-11 |  |
|  |  | THC+CBD | -5.26E-12 | 5.45E-12 | 0.342 | -1.64E-11 | 5.87E-12 |  |
|  | THC+CBD | Vehicle | -2.72E-12 | 5.75E-12 | 0.639 | -1.45E-11 | 9.02E-12 |  |
|  |  | CBD | 1.67583E-011^*^ | 5.75E-12 | 0.007 | 5.02E-12 | 2.85E-11 |  |
|  |  | THC | 5.26E-12 | 5.45E-12 | 0.342 | -5.87E-12 | 1.64E-11 |  |
| *N-*arachidonoyl taurine | Vehicle | CBD | 3.22349E-011^*^ | 1.18E-11 | 0.011 | 8.11E-12 | 5.64E-11 | 3.21 |
|  |  | THC | 1.08E-11 | 1.08E-11 | 0.328 | -1.14E-11 | 3.29E-11 |  |
|  |  | THC+CBD | -5.24623E-011^*^ | 1.11E-11 | 0.000 | -7.51E-11 | -2.98E-11 |  |
|  | CBD | Vehicle | -3.22349E-011^*^ | 1.18E-11 | 0.011 | -5.64E-11 | -8.11E-12 |  |
|  |  | THC | -2.15E-11 | 1.12E-11 | 0.066 | -4.44E-11 | 1.49E-12 |  |
|  |  | THC+CBD | -8.46972E-011^*^ | 1.15E-11 | 0.000 | -1.08E-10 | -6.12E-11 |  |
|  | THC | Vehicle | -1.08E-11 | 1.08E-11 | 0.328 | -3.29E-11 | 1.14E-11 |  |
|  |  | CBD | 2.15E-11 | 1.12E-11 | 0.066 | -1.49E-12 | 4.44E-11 |  |
|  |  | THC+CBD | -6.32177E-011^*^ | 1.05E-11 | 0.000 | -8.46E-11 | -4.18E-11 |  |
|  | THC+CBD | Vehicle | 5.24623E-011^*^ | 1.11E-11 | 0.000 | 2.98E-11 | 7.51E-11 |  |
|  |  | CBD | 8.46972E-011^*^ | 1.15E-11 | 0.000 | 6.12E-11 | 1.08E-10 |  |
|  |  | THC | 6.32177E-011^*^ | 1.05E-11 | 0.000 | 4.18E-11 | 8.46E-11 |  |

### Supplemental Figure 23. ANOVA for the effect of CBD, THC, or THC+CBD (3mg/kg) injections administered to a dam on *N-*acyl tryptophans measured in milk collected from nursing pups’ stomachs

|  | | Sum of Squares | df | Mean Square | F | Sig. |
| --- | --- | --- | --- | --- | --- | --- |
| *N-*oleoyl tryptophan | Between Groups | 0.000 | 3 | 0.000 | 1.178 | 0.337 |
|  | Within Groups | 0.000 | 26 | 0.000 |  |  |
|  | Total | 0.000 | 29 |  |  |  |
| *N-*palmitoyl tryptophan | Between Groups | 0.000 | 3 | 0.000 | 0.588 | 0.628 |
|  | Within Groups | 0.000 | 26 | 0.000 |  |  |
|  | Total | 0.000 | 29 |  |  |  |
| *N-*stearoyl tryptophan | Between Groups | 0.000 | 3 | 0.000 | 4.670 | 0.009 |
|  | Within Groups | 0.000 | 28 | 0.000 |  |  |
|  | Total | 0.000 | 31 |  |  |  |
| *N-*linoleoyl tryptophan | Between Groups | 0.000 | 3 | 0.000 | 6.734 | 0.001 |
|  | Within Groups | 0.000 | 32 | 0.000 |  |  |
|  | Total | 0.000 | 35 |  |  |  |

### Supplemental Figure 24. Post-hoc Fisher’s LSD results for *N-*acyl tryptophans

| Dependent Variable | | | Mean Difference (I-J) | Std. Error | Sig. | 95% Confidence Interval | |  |
| --- | --- | --- | --- | --- | --- | --- | --- | --- |
|  |  |  |  |  |  | Lower Bound | Upper Bound | Magnitude (drug/vehicle) |
| *N-*oleoyl tryptophan | Vehicle | CBD | 1.38E-13 | 4.21E-13 | 0.745 | -7.27E-13 | 1.00E-12 |  |
|  |  | THC | 6.23E-13 | 4.01E-13 | 0.132 | -2.01E-13 | 1.45E-12 |  |
|  |  | THC+CBD | 5.15E-13 | 4.01E-13 | 0.210 | -3.09E-13 | 1.34E-12 |  |
|  | CBD | Vehicle | -1.38E-13 | 4.21E-13 | 0.745 | -1.00E-12 | 7.27E-13 |  |
|  |  | THC | 4.85E-13 | 3.62E-13 | 0.192 | -2.60E-13 | 1.23E-12 |  |
|  |  | THC+CBD | 3.76E-13 | 3.62E-13 | 0.308 | -3.68E-13 | 1.12E-12 |  |
|  | THC | Vehicle | -6.23E-13 | 4.01E-13 | 0.132 | -1.45E-12 | 2.01E-13 |  |
|  |  | CBD | -4.85E-13 | 3.62E-13 | 0.192 | -1.23E-12 | 2.60E-13 |  |
|  |  | THC+CBD | -1.08E-13 | 3.39E-13 | 0.752 | -8.05E-13 | 5.88E-13 |  |
|  | THC+CBD | Vehicle | -5.15E-13 | 4.01E-13 | 0.210 | -1.34E-12 | 3.09E-13 |  |
|  |  | CBD | -3.76E-13 | 3.62E-13 | 0.308 | -1.12E-12 | 3.68E-13 |  |
|  |  | THC | 1.08E-13 | 3.39E-13 | 0.752 | -5.88E-13 | 8.05E-13 |  |
| *N-*palmitoyl tryptophan | Vehicle | CBD | -1.99E-13 | 2.96E-13 | 0.506 | -8.08E-13 | 4.09E-13 |  |
|  |  | THC | -7.20E-16 | 2.86E-13 | 0.998 | -5.89E-13 | 5.87E-13 |  |
|  |  | THC+CBD | 1.17E-13 | 2.96E-13 | 0.696 | -4.92E-13 | 7.25E-13 |  |
|  | CBD | Vehicle | 1.99E-13 | 2.96E-13 | 0.506 | -4.09E-13 | 8.08E-13 |  |
|  |  | THC | 1.99E-13 | 2.29E-13 | 0.394 | -2.73E-13 | 6.70E-13 |  |
|  |  | THC+CBD | 3.16E-13 | 2.42E-13 | 0.202 | -1.81E-13 | 8.13E-13 |  |
|  | THC | Vehicle | 7.20E-16 | 2.86E-13 | 0.998 | -5.87E-13 | 5.89E-13 |  |
|  |  | CBD | -1.99E-13 | 2.29E-13 | 0.394 | -6.70E-13 | 2.73E-13 |  |
|  |  | THC+CBD | 1.18E-13 | 2.29E-13 | 0.613 | -3.54E-13 | 5.89E-13 |  |
|  | THC+CBD | Vehicle | -1.17E-13 | 2.96E-13 | 0.696 | -7.25E-13 | 4.92E-13 |  |
|  |  | CBD | -3.16E-13 | 2.42E-13 | 0.202 | -8.13E-13 | 1.81E-13 |  |
|  |  | THC | -1.18E-13 | 2.29E-13 | 0.613 | -5.89E-13 | 3.54E-13 |  |
| *N-*stearoyl tryptophan | Vehicle | CBD | 5.29E-13 | 3.11E-13 | 0.100 | -1.08E-13 | 1.17E-12 |  |
|  |  | THC | 6.07E-13 | 3.20E-13 | 0.068 | -4.74E-14 | 1.26E-12 | 1.39 |
|  |  | THC+CBD | 1.18884E-012^*^ | 3.20E-13 | 0.001 | 5.34E-13 | 1.84E-12 | 2.23 |
|  | CBD | Vehicle | -5.29E-13 | 3.11E-13 | 0.100 | -1.17E-12 | 1.08E-13 |  |
|  |  | THC | 7.78E-14 | 3.00E-13 | 0.797 | -5.37E-13 | 6.92E-13 |  |
|  |  | THC+CBD | 6.59589E-013^*^ | 3.00E-13 | 0.036 | 4.51E-14 | 1.27E-12 |  |
|  | THC | Vehicle | -6.07E-13 | 3.20E-13 | 0.068 | -1.26E-12 | 4.74E-14 |  |
|  |  | CBD | -7.78E-14 | 3.00E-13 | 0.797 | -6.92E-13 | 5.37E-13 |  |
|  |  | THC+CBD | 5.82E-13 | 3.09E-13 | 0.070 | -5.05E-14 | 1.21E-12 |  |
|  | THC+CBD | Vehicle | -1.18884E-012^*^ | 3.20E-13 | 0.001 | -1.84E-12 | -5.34E-13 |  |
|  |  | CBD | -6.59589E-013^*^ | 3.00E-13 | 0.036 | -1.27E-12 | -4.51E-14 |  |
|  |  | THC | -5.82E-13 | 3.09E-13 | 0.070 | -1.21E-12 | 5.05E-14 |  |
| *N-*linoleoyl tryptophan | Vehicle | CBD | -1.14E-12 | 6.75E-13 | 0.101 | -2.51E-12 | 2.35E-13 |  |
|  |  | THC | -1.78166E-012^*^ | 6.58E-13 | 0.011 | -3.12E-12 | -4.40E-13 | 1.58 |
|  |  | THC+CBD | -2.95218E-012^*^ | 6.75E-13 | 0.000 | -4.33E-12 | -1.58E-12 | 1.97 |
|  | CBD | Vehicle | 1.14E-12 | 6.75E-13 | 0.101 | -2.35E-13 | 2.51E-12 |  |
|  |  | THC | -6.42E-13 | 6.38E-13 | 0.322 | -1.94E-12 | 6.57E-13 |  |
|  |  | THC+CBD | -1.81269E-012^*^ | 6.54E-13 | 0.009 | -3.15E-12 | -4.80E-13 |  |
|  | THC | Vehicle | 1.78166E-012^*^ | 6.58E-13 | 0.011 | 4.40E-13 | 3.12E-12 |  |
|  |  | CBD | 6.42E-13 | 6.38E-13 | 0.322 | -6.57E-13 | 1.94E-12 |  |
|  |  | THC+CBD | -1.17E-12 | 6.38E-13 | 0.076 | -2.47E-12 | 1.29E-13 |  |
|  | THC+CBD | Vehicle | 2.95218E-012^*^ | 6.75E-13 | 0.000 | 1.58E-12 | 4.33E-12 |  |
|  |  | CBD | 1.81269E-012^*^ | 6.54E-13 | 0.009 | 4.80E-13 | 3.15E-12 |  |
|  |  | THC | 1.17E-12 | 6.38E-13 | 0.076 | -1.29E-13 | 2.47E-12 |  |

### Supplemental Figure 25. ANOVA for the effect of CBD, THC, or THC+CBD (3mg/kg) injections administered to a dam on *N-*acyl tyrosines measured in milk collected from nursing pups’ stomachs

|  | | Sum of Squares | df | Mean Square | F | Sig. |
| --- | --- | --- | --- | --- | --- | --- |
| *N-*palmitoyl tyrosine | Between Groups | 0.000 | 3 | 0.000 | 1.008 | 0.403 |
|  | Within Groups | 0.000 | 31 | 0.000 |  |  |
|  | Total | 0.000 | 34 |  |  |  |
| *N-*stearoyl tyrosine | Between Groups | 0.000 | 3 | 0.000 | 2.683 | 0.067 |
|  | Within Groups | 0.000 | 27 | 0.000 |  |  |
|  | Total | 0.000 | 30 |  |  |  |
| *N-*oleoyl tyrosine | Between Groups | 0.000 | 3 | 0.000 | 1.760 | 0.175 |
|  | Within Groups | 0.000 | 32 | 0.000 |  |  |
|  | Total | 0.000 | 35 |  |  |  |
| *N-linoleoyl tyrosine* | Between Groups | 0.000 | 3 | 0.000 | 5.639 | 0.003 |
|  | Within Groups | 0.000 | 32 | 0.000 |  |  |
|  | Total | 0.000 | 35 |  |  |  |
| *N-*docosahexaenoyl tyrosine | Between Groups | 0.000 | 3 | 0.000 | 2.366 | 0.157 |
|  | Within Groups | 0.000 | 7 | 0.000 |  |  |
|  | Total | 0.000 | 10 |  |  |  |

### Supplemental Figure 26. Post-hoc Fisher’s LSD results for *N-*acyl tyrosines

| Dependent Variable | | | Mean Difference (I-J) | Std. Error | Sig. | 95% Confidence Interval | |  |
| --- | --- | --- | --- | --- | --- | --- | --- | --- |
|  |  |  |  |  |  | Lower Bound | Upper Bound | Magnitude (drug/vehicle) |
| *N-*palmitoyl tyrosine | Vehicle | CBD | 1.46E-14 | 1.30E-13 | 0.911 | -2.51E-13 | 2.80E-13 |  |
|  |  | THC | -7.95E-14 | 1.22E-13 | 0.518 | -3.27E-13 | 1.68E-13 |  |
|  |  | THC+CBD | -1.75E-13 | 1.27E-13 | 0.177 | -4.33E-13 | 8.35E-14 |  |
|  | CBD | Vehicle | -1.46E-14 | 1.30E-13 | 0.911 | -2.80E-13 | 2.51E-13 |  |
|  |  | THC | -9.41E-14 | 1.17E-13 | 0.426 | -3.32E-13 | 1.44E-13 |  |
|  |  | THC+CBD | -1.89E-13 | 1.22E-13 | 0.131 | -4.38E-13 | 5.97E-14 |  |
|  | THC | Vehicle | 7.95E-14 | 1.22E-13 | 0.518 | -1.68E-13 | 3.27E-13 |  |
|  |  | CBD | 9.41E-14 | 1.17E-13 | 0.426 | -1.44E-13 | 3.32E-13 |  |
|  |  | THC+CBD | -9.52E-14 | 1.13E-13 | 0.406 | -3.26E-13 | 1.35E-13 |  |
|  | THC+CBD | Vehicle | 1.75E-13 | 1.27E-13 | 0.177 | -8.35E-14 | 4.33E-13 |  |
|  |  | CBD | 1.89E-13 | 1.22E-13 | 0.131 | -5.97E-14 | 4.38E-13 |  |
|  |  | THC | 9.52E-14 | 1.13E-13 | 0.406 | -1.35E-13 | 3.26E-13 |  |
| *N-*stearoyl tyrosine | Vehicle | CBD | 6.08893E-014^*^ | 2.77E-14 | 0.037 | 4.08E-15 | 1.18E-13 | 0.67 |
|  |  | THC | 5.57369E-014^*^ | 2.65E-14 | 0.045 | 1.42E-15 | 1.10E-13 | 0.69 |
|  |  | THC+CBD | 7.64842E-014^*^ | 2.85E-14 | 0.012 | 1.80E-14 | 1.35E-13 | 0.58 |
|  | CBD | Vehicle | -6.08893E-014^*^ | 2.77E-14 | 0.037 | -1.18E-13 | -4.08E-15 |  |
|  |  | THC | -5.15E-15 | 2.43E-14 | 0.834 | -5.50E-14 | 4.47E-14 |  |
|  |  | THC+CBD | 1.56E-14 | 2.65E-14 | 0.562 | -3.88E-14 | 7.00E-14 |  |
|  | THC | Vehicle | -5.57369E-014^*^ | 2.65E-14 | 0.045 | -1.10E-13 | -1.42E-15 |  |
|  |  | CBD | 5.15E-15 | 2.43E-14 | 0.834 | -4.47E-14 | 5.50E-14 |  |
|  |  | THC+CBD | 2.07E-14 | 2.53E-14 | 0.419 | -3.11E-14 | 7.26E-14 |  |
|  | THC+CBD | Vehicle | -7.64842E-014^*^ | 2.85E-14 | 0.012 | -1.35E-13 | -1.80E-14 |  |
|  |  | CBD | -1.56E-14 | 2.65E-14 | 0.562 | -7.00E-14 | 3.88E-14 |  |
|  |  | THC | -2.07E-14 | 2.53E-14 | 0.419 | -7.26E-14 | 3.11E-14 |  |
| *N-*oleoyl tyrosine | Vehicle | CBD | -4.62E-14 | 2.05E-13 | 0.823 | -4.63E-13 | 3.71E-13 |  |
|  |  | THC | -2.66E-13 | 1.90E-13 | 0.171 | -6.54E-13 | 1.21E-13 |  |
|  |  | THC+CBD | -3.92E-13 | 1.99E-13 | 0.058 | -7.97E-13 | 1.34E-14 | 1.47 |
|  | CBD | Vehicle | 4.62E-14 | 2.05E-13 | 0.823 | -3.71E-13 | 4.63E-13 |  |
|  |  | THC | -2.20E-13 | 1.90E-13 | 0.256 | -6.07E-13 | 1.67E-13 |  |
|  |  | THC+CBD | -3.45E-13 | 1.99E-13 | 0.092 | -7.51E-13 | 5.96E-14 |  |
|  | THC | Vehicle | 2.66E-13 | 1.90E-13 | 0.171 | -1.21E-13 | 6.54E-13 |  |
|  |  | CBD | 2.20E-13 | 1.90E-13 | 0.256 | -1.67E-13 | 6.07E-13 |  |
|  |  | THC+CBD | -1.25E-13 | 1.84E-13 | 0.500 | -5.00E-13 | 2.49E-13 |  |
|  | THC+CBD | Vehicle | 3.92E-13 | 1.99E-13 | 0.058 | -1.34E-14 | 7.97E-13 |  |
|  |  | CBD | 3.45E-13 | 1.99E-13 | 0.092 | -5.96E-14 | 7.51E-13 |  |
|  |  | THC | 1.25E-13 | 1.84E-13 | 0.500 | -2.49E-13 | 5.00E-13 |  |
| *N-*linoleoyl tyrosine | Vehicle | CBD | 1.14E-13 | 1.90E-13 | 0.552 | -2.73E-13 | 5.02E-13 |  |
|  |  | THC | -2.02E-13 | 1.77E-13 | 0.261 | -5.62E-13 | 1.58E-13 |  |
|  |  | THC+CBD | -5.86952E-013^*^ | 1.85E-13 | 0.003 | -9.64E-13 | -2.10E-13 | 1.48 |
|  | CBD | Vehicle | -1.14E-13 | 1.90E-13 | 0.552 | -5.02E-13 | 2.73E-13 |  |
|  |  | THC | -3.17E-13 | 1.77E-13 | 0.083 | -6.77E-13 | 4.32E-14 |  |
|  |  | THC+CBD | -7.01407E-013^*^ | 1.85E-13 | 0.001 | -1.08E-12 | -3.25E-13 |  |
|  | THC | Vehicle | 2.02E-13 | 1.77E-13 | 0.261 | -1.58E-13 | 5.62E-13 |  |
|  |  | CBD | 3.17E-13 | 1.77E-13 | 0.083 | -4.32E-14 | 6.77E-13 |  |
|  |  | THC+CBD | -3.84548E-013^*^ | 1.71E-13 | 0.032 | -7.33E-13 | -3.62E-14 |  |
|  | THC+CBD | Vehicle | 5.86952E-013^*^ | 1.85E-13 | 0.003 | 2.10E-13 | 9.64E-13 |  |
|  |  | CBD | 7.01407E-013^*^ | 1.85E-13 | 0.001 | 3.25E-13 | 1.08E-12 |  |
|  |  | THC | 3.84548E-013^*^ | 1.71E-13 | 0.032 | 3.62E-14 | 7.33E-13 |  |
| *N-*docosahexaenoyl tyrosine | Vehicle | CBD | 1.05E-12 | 4.65E-13 | 0.059 | -4.96E-14 | 2.15E-12 | 0.27 |
|  |  | THC | 7.55E-13 | 3.89E-13 | 0.094 | -1.65E-13 | 1.67E-12 | 0.47 |
|  |  | THC+CBD | 9.75E-13 | 4.65E-13 | 0.074 | -1.25E-13 | 2.07E-12 | 0.32 |
|  | CBD | Vehicle | -1.05E-12 | 4.65E-13 | 0.059 | -2.15E-12 | 4.96E-14 |  |
|  |  | THC | -2.95E-13 | 4.41E-13 | 0.525 | -1.34E-12 | 7.48E-13 |  |
|  |  | THC+CBD | -7.50E-14 | 5.09E-13 | 0.887 | -1.28E-12 | 1.13E-12 |  |
|  | THC | Vehicle | -7.55E-13 | 3.89E-13 | 0.094 | -1.67E-12 | 1.65E-13 |  |
|  |  | CBD | 2.95E-13 | 4.41E-13 | 0.525 | -7.48E-13 | 1.34E-12 |  |
|  |  | THC+CBD | 2.20E-13 | 4.41E-13 | 0.633 | -8.23E-13 | 1.26E-12 |  |
|  | THC+CBD | Vehicle | -9.75E-13 | 4.65E-13 | 0.074 | -2.07E-12 | 1.25E-13 |  |
|  |  | CBD | 7.50E-14 | 5.09E-13 | 0.887 | -1.13E-12 | 1.28E-12 |  |
|  |  | THC | -2.20E-13 | 4.41E-13 | 0.633 | -1.26E-12 | 8.23E-13 |  |

#### Supplemental Figure 27. ANOVA for the effect of CBD, THC, or THC+CBD (3mg/kg) injections administered to a dam on *N-*acyl valines measured in milk collected from nursing pups’ stomachs

|  | | Sum of Squares | df | Mean Square | F | Sig. |
| --- | --- | --- | --- | --- | --- | --- |
| *N-*palmitoyl valine | Between Groups | 0.000 | 3 | 0.000 | 1.882 | 0.153 |
|  | Within Groups | 0.000 | 32 | 0.000 |  |  |
|  | Total | 0.000 | 35 |  |  |  |
| *N-*linoleoyl valine | Between Groups | 0.000 | 3 | 0.000 | 7.891 | 0.000 |
|  | Within Groups | 0.000 | 32 | 0.000 |  |  |
|  | Total | 0.000 | 35 |  |  |  |
| *N-*stearoyl valine | Between Groups | 0.000 | 3 | 0.000 | 0.618 | 0.608 |
|  | Within Groups | 0.000 | 33 | 0.000 |  |  |
|  | Total | 0.000 | 36 |  |  |  |
| *N-*oleoyl valine | Between Groups | 0.000 | 3 | 0.000 | 5.519 | 0.004 |
|  | Within Groups | 0.000 | 31 | 0.000 |  |  |
|  | Total | 0.000 | 34 |  |  |  |

### Supplemental Figure 28. Post-hoc Fisher’s LSD results for *N-*acyl valines

| Dependent Variable | | | Mean Difference (I-J) | Std. Error | Sig. | 95% Confidence Interval | |  |
| --- | --- | --- | --- | --- | --- | --- | --- | --- |
|  |  |  |  |  |  | Lower Bound | Upper Bound | Magnitude (drug/vehicle) |
| *N-*palmitoyl valine | Vehicle | CBD | -4.06E-13 | 4.46E-13 | 0.369 | -1.31E-12 | 5.02E-13 |  |
|  |  | THC | -9.23844E-013^*^ | 4.14E-13 | 0.033 | -1.77E-12 | -7.97E-14 | 1.36 |
|  |  | THC+CBD | -7.53E-13 | 4.33E-13 | 0.092 | -1.64E-12 | 1.30E-13 | 1.30 |
|  | CBD | Vehicle | 4.06E-13 | 4.46E-13 | 0.369 | -5.02E-13 | 1.31E-12 |  |
|  |  | THC | -5.18E-13 | 4.14E-13 | 0.221 | -1.36E-12 | 3.26E-13 |  |
|  |  | THC+CBD | -3.47E-13 | 4.33E-13 | 0.429 | -1.23E-12 | 5.36E-13 |  |
|  | THC | Vehicle | 9.23844E-013^*^ | 4.14E-13 | 0.033 | 7.97E-14 | 1.77E-12 |  |
|  |  | CBD | 5.18E-13 | 4.14E-13 | 0.221 | -3.26E-13 | 1.36E-12 |  |
|  |  | THC+CBD | 1.71E-13 | 4.01E-13 | 0.673 | -6.46E-13 | 9.87E-13 |  |
|  | THC+CBD | Vehicle | 7.53E-13 | 4.33E-13 | 0.092 | -1.30E-13 | 1.64E-12 |  |
|  |  | CBD | 3.47E-13 | 4.33E-13 | 0.429 | -5.36E-13 | 1.23E-12 |  |
|  |  | THC | -1.71E-13 | 4.01E-13 | 0.673 | -9.87E-13 | 6.46E-13 |  |
| *N-*linoleoyl valine | Vehicle | CBD | -1.17E-12 | 9.50E-13 | 0.225 | -3.11E-12 | 7.59E-13 |  |
|  |  | THC | -2.34874E-012^*^ | 9.27E-13 | 0.016 | -4.24E-12 | -4.61E-13 | 1.59 |
|  |  | THC+CBD | -4.38340E-012^*^ | 9.50E-13 | 0.000 | -6.32E-12 | -2.45E-12 | 2.10 |
|  | CBD | Vehicle | 1.17E-12 | 9.50E-13 | 0.225 | -7.59E-13 | 3.11E-12 |  |
|  |  | THC | -1.17E-12 | 8.98E-13 | 0.200 | -3.00E-12 | 6.55E-13 |  |
|  |  | THC+CBD | -3.20849E-012^*^ | 9.21E-13 | 0.001 | -5.08E-12 | -1.33E-12 |  |
|  | THC | Vehicle | 2.34874E-012^*^ | 9.27E-13 | 0.016 | 4.61E-13 | 4.24E-12 |  |
|  |  | CBD | 1.17E-12 | 8.98E-13 | 0.200 | -6.55E-13 | 3.00E-12 |  |
|  |  | THC+CBD | -2.03466E-012^*^ | 8.98E-13 | 0.030 | -3.86E-12 | -2.06E-13 |  |
|  | THC+CBD | Vehicle | 4.38340E-012^*^ | 9.50E-13 | 0.000 | 2.45E-12 | 6.32E-12 |  |
|  |  | CBD | 3.20849E-012^*^ | 9.21E-13 | 0.001 | 1.33E-12 | 5.08E-12 |  |
|  |  | THC | 2.03466E-012^*^ | 8.98E-13 | 0.030 | 2.06E-13 | 3.86E-12 |  |
| *N-*stearoyl valine | Vehicle | CBD | -7.00E-14 | 1.23E-13 | 0.574 | -3.21E-13 | 1.81E-13 |  |
|  |  | THC | -1.49E-13 | 1.18E-13 | 0.215 | -3.89E-13 | 9.08E-14 |  |
|  |  | THC+CBD | -3.45E-14 | 1.23E-13 | 0.781 | -2.85E-13 | 2.16E-13 |  |
|  | CBD | Vehicle | 7.00E-14 | 1.23E-13 | 0.574 | -1.81E-13 | 3.21E-13 |  |
|  |  | THC | -7.91E-14 | 1.14E-13 | 0.493 | -3.11E-13 | 1.53E-13 |  |
|  |  | THC+CBD | 3.55E-14 | 1.20E-13 | 0.769 | -2.08E-13 | 2.79E-13 |  |
|  | THC | Vehicle | 1.49E-13 | 1.18E-13 | 0.215 | -9.08E-14 | 3.89E-13 |  |
|  |  | CBD | 7.91E-14 | 1.14E-13 | 0.493 | -1.53E-13 | 3.11E-13 |  |
|  |  | THC+CBD | 1.15E-13 | 1.14E-13 | 0.322 | -1.17E-13 | 3.47E-13 |  |
|  | THC+CBD | Vehicle | 3.45E-14 | 1.23E-13 | 0.781 | -2.16E-13 | 2.85E-13 |  |
|  |  | CBD | -3.55E-14 | 1.20E-13 | 0.769 | -2.79E-13 | 2.08E-13 |  |
|  |  | THC | -1.15E-13 | 1.14E-13 | 0.322 | -3.47E-13 | 1.17E-13 |  |
| *N-*oleoyl valine | Vehicle | CBD | -5.52158E-013^*^ | 2.16E-13 | 0.016 | -9.93E-13 | -1.11E-13 | 1.58 |
|  |  | THC | -6.51849E-013^*^ | 2.11E-13 | 0.004 | -1.08E-12 | -2.21E-13 | 1.68 |
|  |  | THC+CBD | -8.62367E-013^*^ | 2.23E-13 | 0.001 | -1.32E-12 | -4.08E-13 | 1.90 |
|  | CBD | Vehicle | 5.52158E-013^*^ | 2.16E-13 | 0.016 | 1.11E-13 | 9.93E-13 |  |
|  |  | THC | -9.97E-14 | 2.05E-13 | 0.629 | -5.17E-13 | 3.18E-13 |  |
|  |  | THC+CBD | -3.10E-13 | 2.16E-13 | 0.162 | -7.51E-13 | 1.31E-13 |  |
|  | THC | Vehicle | 6.51849E-013^*^ | 2.11E-13 | 0.004 | 2.21E-13 | 1.08E-12 |  |
|  |  | CBD | 9.97E-14 | 2.05E-13 | 0.629 | -3.18E-13 | 5.17E-13 |  |
|  |  | THC+CBD | -2.11E-13 | 2.11E-13 | 0.327 | -6.41E-13 | 2.20E-13 |  |
|  | THC+CBD | Vehicle | 8.62367E-013^*^ | 2.23E-13 | 0.001 | 4.08E-13 | 1.32E-12 |  |
|  |  | CBD | 3.10E-13 | 2.16E-13 | 0.162 | -1.31E-13 | 7.51E-13 |  |
|  |  | THC | 2.11E-13 | 2.11E-13 | 0.327 | -2.20E-13 | 6.41E-13 |  |

### Supplemental Figure 29. ANOVA for the effect of CBD, THC, or THC+CBD (3mg/kg) injections administered to a dam on 2-acyl-glycerols measured in milk collected from nursing pups’ stomachs

|  | | Sum of Squares | df | Mean Square | F | Sig. |
| --- | --- | --- | --- | --- | --- | --- |
| 2-arachidonoyl glycerol | Between Groups | 0.000 | 3 | 0.000 | 5.058 | 0.005 |
|  | Within Groups | 0.000 | 33 | 0.000 |  |  |
|  | Total | 0.000 | 36 |  |  |  |
| 2-palmitoyl glycerol | Between Groups | 0.000 | 3 | 0.000 | 3.997 | 0.016 |
|  | Within Groups | 0.000 | 33 | 0.000 |  |  |
|  | Total | 0.000 | 36 |  |  |  |
| 2-oleoyl glycerol | Between Groups | 0.000 | 3 | 0.000 | 3.192 | 0.036 |
|  | Within Groups | 0.000 | 33 | 0.000 |  |  |
|  | Total | 0.000 | 36 |  |  |  |
| 2-linoleoyl glycerol | Between Groups | 0.000 | 3 | 0.000 | 0.452 | 0.718 |
|  | Within Groups | 0.000 | 33 | 0.000 |  |  |
|  | Total | 0.000 | 36 |  |  |  |

### Supplemental Figure 30. Post-hoc Fisher’s LSD results for 2-acyl-glycerols

| Dependent Variable | | | Mean Difference (I-J) | Std. Error | Sig. | 95% Confidence Interval | |  |
| --- | --- | --- | --- | --- | --- | --- | --- | --- |
|  |  |  |  |  |  | Lower Bound | Upper Bound | Magnitude (drug/vehicle) |
| 2-arachidonoyl glycerol | Vehicle | CBD | -3.65436E-011^*^ | 1.46517E-11 | 0.018 | -6.6353E-11 | -6.7345E-12 | 1.17 |
|  |  | THC | 7.47793E-12 | 1.40109E-11 | 0.597 | -2.1027E-11 | 3.5983E-11 |  |
|  |  | THC+CBD | 1.33194E-11 | 1.46517E-11 | 0.370 | -1.6490E-11 | 4.3128E-11 |  |
|  | CBD | Vehicle | 3.65436E-011^*^ | 1.46517E-11 | 0.018 | 6.7345E-12 | 6.6353E-11 |  |
|  |  | THC | 4.40215E-011^*^ | 1.35527E-11 | 0.003 | 1.6448E-11 | 7.1595E-11 |  |
|  |  | THC+CBD | 4.98630E-011^*^ | 1.42142E-11 | 0.001 | 2.0944E-11 | 7.8782E-11 |  |
|  | THC | Vehicle | -7.47793E-12 | 1.40109E-11 | 0.597 | -3.5983E-11 | 2.1027E-11 |  |
|  |  | CBD | -4.40215E-011^*^ | 1.35527E-11 | 0.003 | -7.1595E-11 | -1.6448E-11 |  |
|  |  | THC+CBD | 5.84148E-12 | 1.35527E-11 | 0.669 | -2.1732E-11 | 3.3415E-11 |  |
|  | THC+CBD | Vehicle | -1.33194E-11 | 1.46517E-11 | 0.370 | -4.3128E-11 | 1.6490E-11 |  |
|  |  | CBD | -4.98630E-011^*^ | 1.42142E-11 | 0.001 | -7.8782E-11 | -2.0944E-11 |  |
|  |  | THC | -5.84148E-12 | 1.35527E-11 | 0.669 | -3.3415E-11 | 2.1732E-11 |  |
| 2-palmitoyl glycerol | Vehicle | CBD | -1.87192E-009^*^ | 7.22398E-10 | 0.014 | -3.3417E-09 | -4.0219E-10 | 1.19 |
|  |  | THC | -4.46032E-10 | 6.90802E-10 | 0.523 | -1.8515E-09 | 9.5942E-10 |  |
|  |  | THC+CBD | 4.15267E-10 | 7.22398E-10 | 0.569 | -1.0545E-09 | 1.8850E-09 |  |
|  | CBD | Vehicle | 1.87192E-009^*^ | 7.22398E-10 | 0.014 | 4.0219E-10 | 3.3417E-09 |  |
|  |  | THC | 1.42589E-009^*^ | 6.68214E-10 | 0.040 | 6.6399E-11 | 2.7854E-09 |  |
|  |  | THC+CBD | 2.28719E-009^*^ | 7.00829E-10 | 0.003 | 8.6134E-10 | 3.7130E-09 |  |
|  | THC | Vehicle | 4.46032E-10 | 6.90802E-10 | 0.523 | -9.5942E-10 | 1.8515E-09 |  |
|  |  | CBD | -1.42589E-009^*^ | 6.68214E-10 | 0.040 | -2.7854E-09 | -6.6399E-11 |  |
|  |  | THC+CBD | 8.61299E-10 | 6.68214E-10 | 0.206 | -4.9819E-10 | 2.2208E-09 |  |
|  | THC+CBD | Vehicle | -4.15267E-10 | 7.22398E-10 | 0.569 | -1.8850E-09 | 1.0545E-09 |  |
|  |  | CBD | -2.28719E-009^*^ | 7.00829E-10 | 0.003 | -3.7130E-09 | -8.6134E-10 |  |
|  |  | THC | -8.61299E-10 | 6.68214E-10 | 0.206 | -2.2208E-09 | 4.9819E-10 |  |
| 2-oleoyl glycerol | Vehicle | CBD | -1.84707E-010^*^ | 6.65780E-11 | 0.009 | -3.2016E-10 | -4.9253E-11 | 1.23 |
|  |  | THC | -1.55243E-010^*^ | 6.36661E-11 | 0.020 | -2.8477E-10 | -2.5714E-11 | 1.20 |
|  |  | THC+CBD | -1.64478E-010^*^ | 6.65780E-11 | 0.019 | -2.9993E-10 | -2.9024E-11 | 1.21 |
|  | CBD | Vehicle | 1.84707E-010^*^ | 6.65780E-11 | 0.009 | 4.9253E-11 | 3.2016E-10 |  |
|  |  | THC | 2.94641E-11 | 6.15843E-11 | 0.635 | -9.5830E-11 | 1.5476E-10 |  |
|  |  | THC+CBD | 2.02296E-11 | 6.45902E-11 | 0.756 | -1.1118E-10 | 1.5164E-10 |  |
|  | THC | Vehicle | 1.55243E-010^*^ | 6.36661E-11 | 0.020 | 2.5714E-11 | 2.8477E-10 |  |
|  |  | CBD | -2.94641E-11 | 6.15843E-11 | 0.635 | -1.5476E-10 | 9.5830E-11 |  |
|  |  | THC+CBD | -9.23457E-12 | 6.15843E-11 | 0.882 | -1.3453E-10 | 1.1606E-10 |  |
|  | THC+CBD | Vehicle | 1.64478E-010^*^ | 6.65780E-11 | 0.019 | 2.9024E-11 | 2.9993E-10 |  |
|  |  | CBD | -2.02296E-11 | 6.45902E-11 | 0.756 | -1.5164E-10 | 1.1118E-10 |  |
|  |  | THC | 9.23457E-12 | 6.15843E-11 | 0.882 | -1.1606E-10 | 1.3453E-10 |  |
| 2-linoleoyl glycerol | Vehicle | CBD | 1.01684E-10 | 1.20099E-10 | 0.403 | -1.4266E-10 | 3.4603E-10 |  |
|  |  | THC | 1.26987E-10 | 1.14846E-10 | 0.277 | -1.0667E-10 | 3.6064E-10 |  |
|  |  | THC+CBD | 6.07551E-11 | 1.20099E-10 | 0.616 | -1.8359E-10 | 3.0510E-10 |  |
|  | CBD | Vehicle | -1.01684E-10 | 1.20099E-10 | 0.403 | -3.4603E-10 | 1.4266E-10 |  |
|  |  | THC | 2.53031E-11 | 1.11090E-10 | 0.821 | -2.0071E-10 | 2.5132E-10 |  |
|  |  | THC+CBD | -4.09289E-11 | 1.16513E-10 | 0.728 | -2.7798E-10 | 1.9612E-10 |  |
|  | THC | Vehicle | -1.26987E-10 | 1.14846E-10 | 0.277 | -3.6064E-10 | 1.0667E-10 |  |
|  |  | CBD | -2.53031E-11 | 1.11090E-10 | 0.821 | -2.5132E-10 | 2.0071E-10 |  |
|  |  | THC+CBD | -6.62320E-11 | 1.11090E-10 | 0.555 | -2.9225E-10 | 1.5978E-10 |  |
|  | THC+CBD | Vehicle | -6.07551E-11 | 1.20099E-10 | 0.616 | -3.0510E-10 | 1.8359E-10 |  |
|  |  | CBD | 4.09289E-11 | 1.16513E-10 | 0.728 | -1.9612E-10 | 2.7798E-10 |  |
|  |  | THC | 6.62320E-11 | 1.11090E-10 | 0.555 | -1.5978E-10 | 2.9225E-10 |  |

### Supplemental 31. ANOVA for the effect of CBD, THC, or THC+CBD (3mg/kg) injections administered to a dam on free fatty acids measured in milk collected from nursing pups’ stomachs

|  | | Sum of Squares | df | Mean Square | F | Sig. |
| --- | --- | --- | --- | --- | --- | --- |
| Oleic acid | Between Groups | 0.000 | 3 | 0.000 | 6.410 | 0.002 |
|  | Within Groups | 0.000 | 32 | 0.000 |  |  |
|  | Total | 0.000 | 35 |  |  |  |
| Linoleoic acid | Between Groups | 0.000 | 3 | 0.000 | 7.325 | 0.001 |
|  | Within Groups | 0.000 | 33 | 0.000 |  |  |
|  | Total | 0.000 | 36 |  |  |  |
| Arachidonic acid | Between Groups | 0.000 | 3 | 0.000 | 8.265 | 0.000 |
|  | Within Groups | 0.000 | 33 | 0.000 |  |  |
|  | Total | 0.000 | 36 |  |  |  |
| Eicosapentaenoic acid | Between Groups | 0.000 | 3 | 0.000 | 6.255 | 0.002 |
|  | Within Groups | 0.000 | 31 | 0.000 |  |  |
|  | Total | 0.000 | 34 |  |  |  |
| Docosahexaenoic acid | Between Groups | 0.000 | 3 | 0.000 | 2.636 | 0.066 |
|  | Within Groups | 0.000 | 33 | 0.000 |  |  |
|  | Total | 0.000 | 36 |  |  |  |

### Supplemental 32. Post-hoc Fisher’s LSD results for free fatty acids

| Dependent Variable | | | Mean Difference (I-J) | Std. Error | Sig. | 95% Confidence Interval | |  |
| --- | --- | --- | --- | --- | --- | --- | --- | --- |
|  |  |  |  |  |  | Lower Bound | Upper Bound | Magnitude (drug/vehicle) |
| Oleic acid | Vehicle | CBD | 3.14635E-008^*^ | 9.17349E-09 | 0.002 | 1.2778E-08 | 5.0149E-08 | 0.54 |
|  |  | THC | 3.47345E-008^*^ | 8.77227E-09 | 0.000 | 1.6866E-08 | 5.2603E-08 | 0.49 |
|  |  | THC+CBD | 3.25336E-008^*^ | 9.43944E-09 | 0.002 | 1.3306E-08 | 5.1761E-08 | 0.52 |
|  | CBD | Vehicle | -3.14635E-008^*^ | 9.17349E-09 | 0.002 | -5.0149E-08 | -1.2778E-08 |  |
|  |  | THC | 3.27101E-09 | 8.48543E-09 | 0.702 | -1.4013E-08 | 2.0555E-08 |  |
|  |  | THC+CBD | 1.07011E-09 | 9.17349E-09 | 0.908 | -1.7616E-08 | 1.9756E-08 |  |
|  | THC | Vehicle | -3.47345E-008^*^ | 8.77227E-09 | 0.000 | -5.2603E-08 | -1.6866E-08 |  |
|  |  | CBD | -3.27101E-09 | 8.48543E-09 | 0.702 | -2.0555E-08 | 1.4013E-08 |  |
|  |  | THC+CBD | -2.20090E-09 | 8.77227E-09 | 0.804 | -2.0069E-08 | 1.5668E-08 |  |
|  | THC+CBD | Vehicle | -3.25336E-008^*^ | 9.43944E-09 | 0.002 | -5.1761E-08 | -1.3306E-08 |  |
|  |  | CBD | -1.07011E-09 | 9.17349E-09 | 0.908 | -1.9756E-08 | 1.7616E-08 |  |
|  |  | THC | 2.20090E-09 | 8.77227E-09 | 0.804 | -1.5668E-08 | 2.0069E-08 |  |
| Linoleoic acid | Vehicle | CBD | 6.40591E-008^*^ | 1.75386E-08 | 0.001 | 2.8377E-08 | 9.9742E-08 | 0.54 |
|  |  | THC | 7.41131E-008^*^ | 1.67715E-08 | 0.000 | 3.9991E-08 | 1.0823E-07 | 0.47 |
|  |  | THC+CBD | 5.78928E-008^*^ | 1.75386E-08 | 0.002 | 2.2210E-08 | 9.3575E-08 | 0.58 |
|  | CBD | Vehicle | -6.40591E-008^*^ | 1.75386E-08 | 0.001 | -9.9742E-08 | -2.8377E-08 |  |
|  |  | THC | 1.00539E-08 | 1.62231E-08 | 0.540 | -2.2952E-08 | 4.3060E-08 |  |
|  |  | THC+CBD | -6.16632E-09 | 1.70149E-08 | 0.719 | -4.0783E-08 | 2.8451E-08 |  |
|  | THC | Vehicle | -7.41131E-008^*^ | 1.67715E-08 | 0.000 | -1.0823E-07 | -3.9991E-08 |  |
|  |  | CBD | -1.00539E-08 | 1.62231E-08 | 0.540 | -4.3060E-08 | 2.2952E-08 |  |
|  |  | THC+CBD | -1.62203E-08 | 1.62231E-08 | 0.325 | -4.9226E-08 | 1.6786E-08 |  |
|  | THC+CBD | Vehicle | -5.78928E-008^*^ | 1.75386E-08 | 0.002 | -9.3575E-08 | -2.2210E-08 |  |
|  |  | CBD | 6.16632E-09 | 1.70149E-08 | 0.719 | -2.8451E-08 | 4.0783E-08 |  |
|  |  | THC | 1.62203E-08 | 1.62231E-08 | 0.325 | -1.6786E-08 | 4.9226E-08 |  |
| Arachidonic acid | Vehicle | CBD | 1.67856E-008^*^ | 5.17323E-09 | 0.003 | 6.2605E-09 | 2.7311E-08 | 0.60 |
|  |  | THC | 2.32123E-008^*^ | 4.94696E-09 | 0.000 | 1.3148E-08 | 3.3277E-08 | 0.45 |
|  |  | THC+CBD | 2.05304E-008^*^ | 5.17323E-09 | 0.000 | 1.0005E-08 | 3.1055E-08 | 0.51 |
|  | CBD | Vehicle | -1.67856E-008^*^ | 5.17323E-09 | 0.003 | -2.7311E-08 | -6.2605E-09 |  |
|  |  | THC | 6.42676E-09 | 4.78521E-09 | 0.188 | -3.3088E-09 | 1.6162E-08 |  |
|  |  | THC+CBD | 3.74482E-09 | 5.01877E-09 | 0.461 | -6.4659E-09 | 1.3956E-08 |  |
|  | THC | Vehicle | -2.32123E-008^*^ | 4.94696E-09 | 0.000 | -3.3277E-08 | -1.3148E-08 |  |
|  |  | CBD | -6.42676E-09 | 4.78521E-09 | 0.188 | -1.6162E-08 | 3.3088E-09 |  |
|  |  | THC+CBD | -2.68194E-09 | 4.78521E-09 | 0.579 | -1.2418E-08 | 7.0536E-09 |  |
|  | THC+CBD | Vehicle | -2.05304E-008^*^ | 5.17323E-09 | 0.000 | -3.1055E-08 | -1.0005E-08 |  |
|  |  | CBD | -3.74482E-09 | 5.01877E-09 | 0.461 | -1.3956E-08 | 6.4659E-09 |  |
|  |  | THC | 2.68194E-09 | 4.78521E-09 | 0.579 | -7.0536E-09 | 1.2418E-08 |  |
| Eicosapentaenoic acid | Vehicle | CBD | 1.96030E-009^*^ | 7.95365E-10 | 0.019 | 3.3815E-10 | 3.5825E-09 | 0.75 |
|  |  | THC | 2.67484E-009^*^ | 7.76425E-10 | 0.002 | 1.0913E-09 | 4.2584E-09 | 0.66 |
|  |  | THC+CBD | 3.30993E-009^*^ | 8.18424E-10 | 0.000 | 1.6407E-09 | 4.9791E-09 | 0.58 |
|  | CBD | Vehicle | -1.96030E-009^*^ | 7.95365E-10 | 0.019 | -3.5825E-09 | -3.3815E-10 |  |
|  |  | THC | 7.14532E-10 | 7.52079E-10 | 0.349 | -8.1934E-10 | 2.2484E-09 |  |
|  |  | THC+CBD | 1.34963E-09 | 7.95365E-10 | 0.100 | -2.7253E-10 | 2.9718E-09 |  |
|  | THC | Vehicle | -2.67484E-009^*^ | 7.76425E-10 | 0.002 | -4.2584E-09 | -1.0913E-09 |  |
|  |  | CBD | -7.14532E-10 | 7.52079E-10 | 0.349 | -2.2484E-09 | 8.1934E-10 |  |
|  |  | THC+CBD | 6.35098E-10 | 7.76425E-10 | 0.420 | -9.4843E-10 | 2.2186E-09 |  |
|  | THC+CBD | Vehicle | -3.30993E-009^*^ | 8.18424E-10 | 0.000 | -4.9791E-09 | -1.6407E-09 |  |
|  |  | CBD | -1.34963E-09 | 7.95365E-10 | 0.100 | -2.9718E-09 | 2.7253E-10 |  |
|  |  | THC | -6.35098E-10 | 7.76425E-10 | 0.420 | -2.2186E-09 | 9.4843E-10 |  |
| Docosahexaenoic acid | Vehicle | CBD | 2.52111E-09 | 1.28420E-09 | 0.058 | -9.1620E-11 | 5.1338E-09 | 0.67 |
|  |  | THC | 3.05075E-009^*^ | 1.22804E-09 | 0.018 | 5.5230E-10 | 5.5492E-09 | 0.60 |
|  |  | THC+CBD | 3.13546E-009^*^ | 1.28420E-09 | 0.020 | 5.2273E-10 | 5.7482E-09 | 0.59 |
|  | CBD | Vehicle | -2.52111E-09 | 1.28420E-09 | 0.058 | -5.1338E-09 | 9.1620E-11 |  |
|  |  | THC | 5.29642E-10 | 1.18788E-09 | 0.659 | -1.8871E-09 | 2.9464E-09 |  |
|  |  | THC+CBD | 6.14350E-10 | 1.24586E-09 | 0.625 | -1.9204E-09 | 3.1491E-09 |  |
|  | THC | Vehicle | -3.05075E-009^*^ | 1.22804E-09 | 0.018 | -5.5492E-09 | -5.5230E-10 |  |
|  |  | CBD | -5.29642E-10 | 1.18788E-09 | 0.659 | -2.9464E-09 | 1.8871E-09 |  |
|  |  | THC+CBD | 8.47085E-11 | 1.18788E-09 | 0.944 | -2.3321E-09 | 2.5015E-09 |  |
|  | THC+CBD | Vehicle | -3.13546E-009^*^ | 1.28420E-09 | 0.020 | -5.7482E-09 | -5.2273E-10 |  |
|  |  | CBD | -6.14350E-10 | 1.24586E-09 | 0.625 | -3.1491E-09 | 1.9204E-09 |  |
|  |  | THC | -8.47085E-11 | 1.18788E-09 | 0.944 | -2.5015E-09 | 2.3321E-09 |  |

### Supplemental Figure 33. Descriptive statistics for CBD and THC measured in milk taken from stomachs of pups nursing dams injected with 3mg/kg CBD, THC, or THC+CBD

|  | N | Mean | Std. Deviation | Std. Error | 95% Confidence Interval for Mean | | Minimum | Maximum |
| --- | --- | --- | --- | --- | --- | --- | --- | --- |
|  |  |  |  |  | Lower Bound | Upper Bound |  |  |
| CBD | 9 | 1.7386E-10 | 7.13885E-11 | 2.37962E-11 | 1.1899E-10 | 2.2874E-10 | 8.74E-11 | 2.94E-10 |
| THC | 11 | 6.9996E-11 | 1.76246E-11 | 5.31402E-12 | 5.8156E-11 | 8.1837E-11 | 5.06E-11 | 1.10E-10 |
| CBD in THC+CBD | 9 | 2.6861E-10 | 8.99802E-11 | 2.99934E-11 | 1.9944E-10 | 3.3777E-10 | 1.54E-10 | 4.11E-10 |
| THC in THC+CBD | 9 | 1.1395E-10 | 3.14295E-11 | 1.04765E-11 | 8.9794E-11 | 1.3811E-10 | 7.35E-11 | 1.65E-10 |

### Supplemental Figure 34. ANOVA for the effect of CBD, THC, or THC+CBD (3mg/kg) injections administered to a dam on cannabinoids measured in milk collected from nursing pups’ stomachs

|  | Sum of Squares | df | Mean Square | F | Sig. |
| --- | --- | --- | --- | --- | --- |
| Between Groups | 0.000 | 3 | 0.000 | 20.778 | 0.000 |
| Within Groups | 0.000 | 34 | 0.000 |  |  |
| Total | 0.000 | 37 |  |  |  |

### Supplemental Figure 35. Post-hoc Fisher’s LSD results for cannabinoids

| (I) TxGrp | | Mean Difference (I-J) | Std. Error | Sig. | 95% Confidence Interval | |
| --- | --- | --- | --- | --- | --- | --- |
|  |  |  |  |  | Lower Bound | Upper Bound |
| CBD | THC | 1.03867E-010^*^ | 2.63157E-11 | 0.000 | 5.0387E-11 | 1.5735E-10 |
|  | CBD in THC+CBD | -9.47438E-011^*^ | 2.76002E-11 | 0.002 | -1.5083E-10 | -3.8654E-11 |
|  | THC in THC+CBD | 5.99107E-011^*^ | 2.76002E-11 | 0.037 | 3.8204E-12 | 1.1600E-10 |
| THC | CBD | -1.03867E-010^*^ | 2.63157E-11 | 0.000 | -1.5735E-10 | -5.0387E-11 |
|  | CBD in THC+CBD | -1.98611E-010^*^ | 2.63157E-11 | 0.000 | -2.5209E-10 | -1.4513E-10 |
|  | THC in THC+CBD | -4.39565E-11 | 2.63157E-11 | 0.104 | -9.7437E-11 | 9.5235E-12 |
| CBD in THC+CBD | CBD | 9.47438E-011^*^ | 2.76002E-11 | 0.002 | 3.8654E-11 | 1.5083E-10 |
|  | THC | 1.98611E-010^*^ | 2.63157E-11 | 0.000 | 1.4513E-10 | 2.5209E-10 |
|  | THC in THC+CBD | 1.54655E-010^*^ | 2.76002E-11 | 0.000 | 9.8564E-11 | 2.1074E-10 |
| THC in THC+CBD | CBD | -5.99107E-011^*^ | 2.76002E-11 | 0.037 | -1.1600E-10 | -3.8204E-12 |
|  | THC | 4.39565E-11 | 2.63157E-11 | 0.104 | -9.5235E-12 | 9.7437E-11 |
|  | CBD in THC+CBD | -1.54655E-010^*^ | 2.76002E-11 | 0.000 | -2.1074E-10 | -9.8564E-11 |

### Supplemental Figure 36: Descriptive statistics for 11-OH-THC and 7-OH-CBD measured in milk from the stomachs of pups nursing a dam injected with (3mg/kg) CBD, THC, or THC+CBD

|  |  | N | Mean | Std. Deviation | Std. Error | 95% Confidence Interval for Mean | | Minimum | Maximum |
| --- | --- | --- | --- | --- | --- | --- | --- | --- | --- |
|  |  |  |  |  |  | Lower Bound | Upper Bound |  |  |
| 11-OH-THC | THC | 10 | 5.49E-12 | 1.90E-12 | 6.02E-13 | 4.13E-12 | 6.86E-12 | 3.50E-12 | 1.02E-11 |
|  | THC+CBD | 9 | 3.91E-11 | 1.18E-11 | 3.93E-12 | 3.00E-11 | 4.81E-11 | 1.73E-11 | 5.50E-11 |
| 7-OH-CBD | CBD | 9 | 2.11E-11 | 8.75E-12 | 2.92E-12 | 1.44E-11 | 2.78E-11 | 6.48E-12 | 3.27E-11 |
|  | THC+CBD | 9 | 7.21E-11 | 1.60E-11 | 5.35E-12 | 5.98E-11 | 8.45E-11 | 5.28E-11 | 1.03E-10 |

### Supplemental Figure 37: ANOVA for the effect of CBD, THC, or THC+CBD (3mg/kg) injections administered to a dam on 11-OH-THC measured in milk collected from nursing pups’ stomachs

|  | Sum of Squares | df | Mean Square | F | Sig. |
| --- | --- | --- | --- | --- | --- |
| Between Groups | 0.000 | 1 | 0.000 | 79.373 | 0.000 |
| Within Groups | 0.000 | 17 | 0.000 |  |  |
| Total | 0.000 | 18 |  |  |  |

### Supplemental Figure 38: ANOVA for the effect of CBD, THC, or THC+CBD (3mg/kg) injections administered to a nursing dam on 7-OH-CBD measured in milk collected from pups’ stomachs

|  | Sum of Squares | df | Mean Square | F | Sig. |
| --- | --- | --- | --- | --- | --- |
| Between Groups | 0.000 | 1 | 0.000 | 70.103 | 0.000 |
| Within Groups | 0.000 | 16 | 0.000 |  |  |
| Total | 0.000 | 17 |  |  |  |

### Supplemental Figure 39: Example chromatograms identifying 7-OH-CBD in standard and in milk samples from each treatment group.

(A) A standard consisting of 10nM 7-OH-CBD was scanned via MS/MS in negative ion mode for a parent mass of [329.5]^-^ paired with a daughter mass of [173]^-^. The peak retention time was 4.99 min. (B) 7-OH-CBD was identified in a sample from a pup in the CBD treatment group based on the peak present in negative ion mode [329.5/173]^-^ that had a retention time at 4.99 min. (C) 7-OH-CBD was identified in the THC+CBD sample by the matching peak in the negative ion [329.5/173]^-^ with a retention time of 4.99 min. (D) 7-OH-CBD was not identified in the THC alone sample. There was no peak in negative ion mode for [329.5/173]^-^ at the retention time 4.99 min. (E) 7-OH-CBD was not identified in the vehicle sample. There was no peak in negative ion mode for [329.5/173]^-^ at the retention time 4.99 min.

392.5/173

4.99

A

Intensity (cps)

| Time (min) | 7-OH-CBD |
| --- | --- |
| 0 | 0 |
| 0.018 | 0 |
| 0.036 | 6.7 |
| 0.054 | 0 |
| 0.072 | 0 |
| 0.09 | 0 |
| 0.109 | 0 |
| 0.127 | 0 |
| 0.145 | 0 |
| 0.163 | 6.7 |
| 0.181 | 6.7 |
| 0.199 | 0 |
| 0.217 | 0 |
| 0.235 | 6.7 |
| 0.253 | 0 |
| 0.271 | 0 |
| 0.289 | 6.7 |
| 0.307 | 0 |
| 0.326 | 0 |
| 0.344 | 0 |
| 0.362 | 0 |
| 0.38 | 0 |
| 0.398 | 0 |
| 0.416 | 0 |
| 0.434 | 6.7 |
| 0.452 | 0 |
| 0.47 | 0 |
| 0.488 | 0 |
| 0.506 | 0 |
| 0.525 | 6.7 |
| 0.543 | 0 |
| 0.561 | 6.7 |
| 0.579 | 0 |
| 0.597 | 6.7 |
| 0.615 | 0 |
| 0.633 | 0 |
| 0.651 | 6.7 |
| 0.669 | 0 |
| 0.687 | 0 |
| 0.705 | 0 |
| 0.723 | 6.7 |
| 0.742 | 0 |
| 0.76 | 6.7 |
| 0.778 | 0 |
| 0.796 | 0 |
| 0.814 | 0 |
| 0.832 | 0 |
| 0.85 | 0 |
| 0.868 | 0 |
| 0.886 | 6.7 |
| 0.904 | 0 |
| 0.922 | 0 |
| 0.94 | 0 |
| 0.959 | 6.7 |
| 0.977 | 0 |
| 0.995 | 0 |
| 1.013 | 20 |
| 1.031 | 0 |
| 1.049 | 13.3 |
| 1.067 | 6.7 |
| 1.085 | 0 |
| 1.103 | 0 |
| 1.121 | 0 |
| 1.139 | 0 |
| 1.157 | 26.7 |
| 1.176 | 6.7 |
| 1.194 | 0 |
| 1.212 | 6.7 |
| 1.23 | 0 |
| 1.248 | 0 |
| 1.266 | 0 |
| 1.284 | 0 |
| 1.302 | 0 |
| 1.32 | 0 |
| 1.338 | 0 |
| 1.356 | 13.3 |
| 1.375 | 0 |
| 1.393 | 0 |
| 1.411 | 0 |
| 1.429 | 6.7 |
| 1.447 | 0 |
| 1.465 | 13.3 |
| 1.483 | 0 |
| 1.501 | 0 |
| 1.519 | 0 |
| 1.537 | 0 |
| 1.555 | 6.7 |
| 1.573 | 0 |
| 1.592 | 0 |
| 1.61 | 20 |
| 1.628 | 0 |
| 1.646 | 20 |
| 1.664 | 0 |
| 1.682 | 0 |
| 1.7 | 20 |
| 1.718 | 0 |
| 1.736 | 0 |
| 1.754 | 6.7 |
| 1.772 | 6.7 |
| 1.79 | 0 |
| 1.809 | 6.7 |
| 1.827 | 0 |
| 1.845 | 0 |
| 1.863 | 0 |
| 1.881 | 0 |
| 1.899 | 0 |
| 1.917 | 0 |
| 1.935 | 0 |
| 1.953 | 0 |
| 1.971 | 0 |
| 1.989 | 0 |
| 2.008 | 26.7 |
| 2.026 | 0 |
| 2.044 | 0 |
| 2.062 | 6.7 |
| 2.08 | 0 |
| 2.098 | 0 |
| 2.116 | 6.7 |
| 2.134 | 6.7 |
| 2.152 | 6.7 |
| 2.17 | 0 |
| 2.188 | 0 |
| 2.206 | 0 |
| 2.225 | 0 |
| 2.243 | 0 |
| 2.261 | 0 |
| 2.279 | 0 |
| 2.297 | 0 |
| 2.315 | 0 |
| 2.333 | 0 |
| 2.351 | 0 |
| 2.369 | 6.7 |
| 2.387 | 0 |
| 2.405 | 0 |
| 2.423 | 0 |
| 2.442 | 6.7 |
| 2.46 | 6.7 |
| 2.478 | 0 |
| 2.496 | 0 |
| 2.514 | 6.7 |
| 2.532 | 0 |
| 2.55 | 6.7 |
| 2.568 | 0 |
| 2.586 | 0 |
| 2.604 | 0 |
| 2.622 | 6.7 |
| 2.64 | 6.7 |
| 2.659 | 0 |
| 2.677 | 13.3 |
| 2.695 | 0 |
| 2.713 | 0 |
| 2.731 | 0 |
| 2.749 | 6.7 |
| 2.767 | 0 |
| 2.785 | 0 |
| 2.803 | 6.7 |
| 2.821 | 6.7 |
| 2.839 | 0 |
| 2.857 | 6.7 |
| 2.876 | 0 |
| 2.894 | 6.7 |
| 2.912 | 0 |
| 2.93 | 0 |
| 2.948 | 6.7 |
| 2.966 | 0 |
| 2.984 | 0 |
| 3.002 | 0 |
| 3.02 | 0 |
| 3.038 | 6.7 |
| 3.056 | 0 |
| 3.075 | 0 |
| 3.093 | 0 |
| 3.111 | 6.7 |
| 3.129 | 0 |
| 3.147 | 0 |
| 3.165 | 0 |
| 3.183 | 6.7 |
| 3.201 | 0 |
| 3.219 | 6.7 |
| 3.237 | 0 |
| 3.255 | 0 |
| 3.273 | 0 |
| 3.292 | 6.7 |
| 3.31 | 0 |
| 3.328 | 0 |
| 3.346 | 0 |
| 3.364 | 0 |
| 3.382 | 0 |
| 3.4 | 0 |
| 3.418 | 0 |
| 3.436 | 0 |
| 3.454 | 0 |
| 3.472 | 0 |
| 3.49 | 0 |
| 3.509 | 0 |
| 3.527 | 6.7 |
| 3.545 | 0 |
| 3.563 | 0 |
| 3.581 | 6.7 |
| 3.599 | 0 |

Intensity (cps)

| Time (min) | CBD |
| --- | --- |
| 0 | 0 |
| 0.018 | 13.3 |
| 0.036 | 0 |
| 0.054 | 0 |
| 0.072 | 20 |
| 0.09 | 0 |
| 0.109 | 0 |
| 0.127 | 6.7 |
| 0.145 | 0 |
| 0.163 | 0 |
| 0.181 | 0 |
| 0.199 | 0 |
| 0.217 | 0 |
| 0.235 | 0 |
| 0.253 | 0 |
| 0.271 | 0 |
| 0.289 | 6.7 |
| 0.307 | 0 |
| 0.326 | 0 |
| 0.344 | 0 |
| 0.362 | 0 |
| 0.38 | 6.7 |
| 0.398 | 0 |
| 0.416 | 0 |
| 0.434 | 0 |
| 0.452 | 6.7 |
| 0.47 | 0 |
| 0.488 | 13.3 |
| 0.506 | 13.3 |
| 0.525 | 6.7 |
| 0.543 | 0 |
| 0.561 | 0 |
| 0.579 | 0 |
| 0.597 | 0 |
| 0.615 | 0 |
| 0.633 | 0 |
| 0.651 | 0 |
| 0.669 | 0 |
| 0.687 | 0 |
| 0.705 | 0 |
| 0.723 | 6.7 |
| 0.742 | 0 |
| 0.76 | 6.7 |
| 0.778 | 6.7 |
| 0.796 | 0 |
| 0.814 | 6.7 |
| 0.832 | 6.7 |
| 0.85 | 0 |
| 0.868 | 0 |
| 0.886 | 6.7 |
| 0.904 | 6.7 |
| 0.922 | 6.7 |
| 0.94 | 0 |
| 0.959 | 0 |
| 0.977 | 0 |
| 0.995 | 0 |
| 1.013 | 6.7 |
| 1.031 | 0 |
| 1.049 | 0 |
| 1.067 | 6.7 |
| 1.085 | 6.7 |
| 1.103 | 6.7 |
| 1.121 | 0 |
| 1.139 | 0 |
| 1.157 | 0 |
| 1.176 | 0 |
| 1.194 | 0 |
| 1.212 | 6.7 |
| 1.23 | 13.3 |
| 1.248 | 0 |
| 1.266 | 0 |
| 1.284 | 0 |
| 1.302 | 13.3 |
| 1.32 | 0 |
| 1.338 | 13.3 |
| 1.356 | 6.7 |
| 1.375 | 0 |
| 1.393 | 0 |
| 1.411 | 6.7 |
| 1.429 | 0 |
| 1.447 | 0 |
| 1.465 | 0 |
| 1.483 | 6.7 |
| 1.501 | 0 |
| 1.519 | 26.7 |
| 1.537 | 13.3 |
| 1.555 | 6.7 |
| 1.573 | 13.3 |
| 1.592 | 20 |
| 1.61 | 13.3 |
| 1.628 | 0 |
| 1.646 | 0 |
| 1.664 | 6.7 |
| 1.682 | 0 |
| 1.7 | 0 |
| 1.718 | 6.7 |
| 1.736 | 0 |
| 1.754 | 0 |
| 1.772 | 20 |
| 1.79 | 0 |
| 1.809 | 0 |
| 1.827 | 13.3 |
| 1.845 | 0 |
| 1.863 | 0 |
| 1.881 | 0 |
| 1.899 | 0 |
| 1.917 | 0 |
| 1.935 | 0 |
| 1.953 | 6.7 |
| 1.971 | 0 |
| 1.989 | 0 |
| 2.008 | 0 |
| 2.026 | 0 |
| 2.044 | 0 |
| 2.062 | 6.7 |
| 2.08 | 0 |
| 2.098 | 0 |
| 2.116 | 6.7 |
| 2.134 | 6.7 |
| 2.152 | 0 |
| 2.17 | 20 |
| 2.188 | 0 |
| 2.206 | 0 |
| 2.225 | 0 |
| 2.243 | 6.7 |
| 2.261 | 0 |
| 2.279 | 6.7 |
| 2.297 | 0 |
| 2.315 | 0 |
| 2.333 | 20 |
| 2.351 | 0 |
| 2.369 | 6.7 |
| 2.387 | 6.7 |
| 2.405 | 13.3 |
| 2.423 | 0 |
| 2.442 | 0 |
| 2.46 | 0 |
| 2.478 | 0 |
| 2.496 | 0 |
| 2.514 | 0 |
| 2.532 | 0 |
| 2.55 | 6.7 |
| 2.568 | 0 |
| 2.586 | 0 |
| 2.604 | 6.7 |
| 2.622 | 20 |
| 2.64 | 0 |
| 2.659 | 6.7 |
| 2.677 | 0 |
| 2.695 | 0 |
| 2.713 | 0 |
| 2.731 | 6.7 |
| 2.749 | 0 |
| 2.767 | 6.7 |
| 2.785 | 0 |
| 2.803 | 0 |
| 2.821 | 0 |
| 2.839 | 0 |
| 2.857 | 0 |
| 2.876 | 0 |
| 2.894 | 0 |
| 2.912 | 6.7 |
| 2.93 | 0 |
| 2.948 | 6.7 |
| 2.966 | 13.3 |
| 2.984 | 0 |
| 3.002 | 0 |
| 3.02 | 6.7 |
| 3.038 | 0 |
| 3.056 | 6.7 |
| 3.075 | 6.7 |
| 3.093 | 0 |
| 3.111 | 0 |
| 3.129 | 0 |
| 3.147 | 0 |
| 3.165 | 0 |
| 3.183 | 0 |
| 3.201 | 0 |
| 3.219 | 0 |
| 3.237 | 20 |
| 3.255 | 0 |
| 3.273 | 0 |
| 3.292 | 13.3 |
| 3.31 | 6.7 |
| 3.328 | 0 |
| 3.346 | 0 |
| 3.364 | 6.7 |
| 3.382 | 0 |
| 3.4 | 6.7 |
| 3.418 | 0 |
| 3.436 | 6.7 |
| 3.454 | 0 |
| 3.472 | 0 |
| 3.49 | 0 |
| 3.509 | 0 |
| 3.527 | 6.7 |
| 3.545 | 0 |
| 3.563 | 0 |
| 3.581 | 0 |
| 3.599 | 0 |

C

THC+CBD

Intensity (cps)

| Time (min) | T+C |
| --- | --- |
| 0 | 0 |
| 0.018 | 0 |
| 0.036 | 0 |
| 0.054 | 0 |
| 0.072 | 0 |
| 0.09 | 0 |
| 0.109 | 0 |
| 0.127 | 0 |
| 0.145 | 0 |
| 0.163 | 6.7 |
| 0.181 | 0 |
| 0.199 | 0 |
| 0.217 | 6.7 |
| 0.235 | 6.7 |
| 0.253 | 0 |
| 0.271 | 0 |
| 0.289 | 0 |
| 0.307 | 0 |
| 0.326 | 0 |
| 0.344 | 0 |
| 0.362 | 6.7 |
| 0.38 | 6.7 |
| 0.398 | 6.7 |
| 0.416 | 0 |
| 0.434 | 0 |
| 0.452 | 0 |
| 0.47 | 0 |
| 0.488 | 0 |
| 0.506 | 0 |
| 0.525 | 0 |
| 0.543 | 0 |
| 0.561 | 0 |
| 0.579 | 13.3 |
| 0.597 | 0 |
| 0.615 | 0 |
| 0.633 | 0 |
| 0.651 | 0 |
| 0.669 | 0 |
| 0.687 | 0 |
| 0.705 | 0 |
| 0.723 | 0 |
| 0.742 | 0 |
| 0.76 | 0 |
| 0.778 | 13.3 |
| 0.796 | 0 |
| 0.814 | 0 |
| 0.832 | 0 |
| 0.85 | 0 |
| 0.868 | 0 |
| 0.886 | 0 |
| 0.904 | 6.7 |
| 0.922 | 0 |
| 0.94 | 6.7 |
| 0.959 | 0 |
| 0.977 | 0 |
| 0.995 | 0 |
| 1.013 | 0 |
| 1.031 | 0 |
| 1.049 | 0 |
| 1.067 | 0 |
| 1.085 | 0 |
| 1.103 | 0 |
| 1.121 | 0 |
| 1.139 | 6.7 |
| 1.157 | 0 |
| 1.176 | 0 |
| 1.194 | 0 |
| 1.212 | 0 |
| 1.23 | 0 |
| 1.248 | 0 |
| 1.266 | 0 |
| 1.284 | 0 |
| 1.302 | 0 |
| 1.32 | 0 |
| 1.338 | 6.7 |
| 1.356 | 0 |
| 1.375 | 0 |
| 1.393 | 0 |
| 1.411 | 0 |
| 1.429 | 6.7 |
| 1.447 | 6.7 |
| 1.465 | 0 |
| 1.483 | 0 |
| 1.501 | 0 |
| 1.519 | 6.7 |
| 1.537 | 6.7 |
| 1.555 | 0 |
| 1.573 | 0 |
| 1.592 | 0 |
| 1.61 | 0 |
| 1.628 | 13.3 |
| 1.646 | 6.7 |
| 1.664 | 0 |
| 1.682 | 13.3 |
| 1.7 | 0 |
| 1.718 | 20 |
| 1.736 | 13.3 |
| 1.754 | 13.3 |
| 1.772 | 0 |
| 1.79 | 0 |
| 1.809 | 6.7 |
| 1.827 | 0 |
| 1.845 | 0 |
| 1.863 | 0 |
| 1.881 | 0 |
| 1.899 | 0 |
| 1.917 | 0 |
| 1.935 | 0 |
| 1.953 | 6.7 |
| 1.971 | 0 |
| 1.989 | 0 |
| 2.008 | 6.7 |
| 2.026 | 0 |
| 2.044 | 0 |
| 2.062 | 0 |
| 2.08 | 0 |
| 2.098 | 0 |
| 2.116 | 0 |
| 2.134 | 0 |
| 2.152 | 0 |
| 2.17 | 0 |
| 2.188 | 0 |
| 2.206 | 0 |
| 2.225 | 0 |
| 2.243 | 0 |
| 2.261 | 6.7 |
| 2.279 | 0 |
| 2.297 | 0 |
| 2.315 | 0 |
| 2.333 | 0 |
| 2.351 | 6.7 |
| 2.369 | 0 |
| 2.387 | 0 |
| 2.405 | 13.3 |
| 2.423 | 0 |
| 2.442 | 0 |
| 2.46 | 0 |
| 2.478 | 6.7 |
| 2.496 | 0 |
| 2.514 | 0 |
| 2.532 | 0 |
| 2.55 | 0 |
| 2.568 | 0 |
| 2.586 | 6.7 |
| 2.604 | 0 |
| 2.622 | 0 |
| 2.64 | 0 |
| 2.659 | 0 |
| 2.677 | 0 |
| 2.695 | 0 |
| 2.713 | 0 |
| 2.731 | 0 |
| 2.749 | 6.7 |
| 2.767 | 0 |
| 2.785 | 0 |
| 2.803 | 6.7 |
| 2.821 | 0 |
| 2.839 | 0 |
| 2.858 | 0 |
| 2.876 | 6.7 |
| 2.894 | 0 |
| 2.912 | 0 |
| 2.93 | 0 |
| 2.948 | 0 |
| 2.966 | 0 |
| 2.984 | 0 |
| 3.002 | 6.7 |
| 3.02 | 0 |
| 3.038 | 0 |
| 3.056 | 13.3 |
| 3.075 | 0 |
| 3.093 | 0 |
| 3.111 | 6.7 |
| 3.129 | 0 |
| 3.147 | 0 |
| 3.165 | 0 |
| 3.183 | 0 |
| 3.201 | 6.7 |
| 3.219 | 0 |
| 3.237 | 0 |
| 3.255 | 0 |
| 3.273 | 0 |
| 3.292 | 0 |
| 3.31 | 0 |
| 3.328 | 6.7 |
| 3.346 | 0 |
| 3.364 | 0 |
| 3.382 | 0 |
| 3.4 | 0 |
| 3.418 | 0 |
| 3.436 | 6.7 |
| 3.454 | 0 |
| 3.472 | 0 |
| 3.49 | 0 |
| 3.509 | 0 |
| 3.527 | 0 |
| 3.545 | 0 |
| 3.563 | 0 |
| 3.581 | 0 |
| 3.599 | 0 |

Intensity (cps)

| Time (min) | THC |
| --- | --- |
| 0 | 6.7 |
| 0.018 | 0 |
| 0.036 | 6.7 |
| 0.054 | 0 |
| 0.072 | 13.3 |
| 0.09 | 6.7 |
| 0.109 | 0 |
| 0.127 | 0 |
| 0.145 | 0 |
| 0.163 | 0 |
| 0.181 | 6.7 |
| 0.199 | 13.3 |
| 0.217 | 0 |
| 0.235 | 6.7 |
| 0.253 | 0 |
| 0.271 | 0 |
| 0.289 | 0 |
| 0.307 | 0 |
| 0.326 | 0 |
| 0.344 | 0 |
| 0.362 | 13.3 |
| 0.38 | 0 |
| 0.398 | 0 |
| 0.416 | 0 |
| 0.434 | 6.7 |
| 0.452 | 0 |
| 0.47 | 6.7 |
| 0.488 | 6.7 |
| 0.506 | 0 |
| 0.525 | 6.7 |
| 0.543 | 0 |
| 0.561 | 0 |
| 0.579 | 0 |
| 0.597 | 0 |
| 0.615 | 13.3 |
| 0.633 | 0 |
| 0.651 | 0 |
| 0.669 | 13.3 |
| 0.687 | 0 |
| 0.705 | 0 |
| 0.723 | 0 |
| 0.742 | 0 |
| 0.76 | 0 |
| 0.778 | 0 |
| 0.796 | 6.7 |
| 0.814 | 0 |
| 0.832 | 0 |
| 0.85 | 0 |
| 0.868 | 0 |
| 0.886 | 0 |
| 0.904 | 6.7 |
| 0.922 | 6.7 |
| 0.94 | 0 |
| 0.959 | 6.7 |
| 0.977 | 0 |
| 0.995 | 0 |
| 1.013 | 0 |
| 1.031 | 6.7 |
| 1.049 | 0 |
| 1.067 | 0 |
| 1.085 | 6.7 |
| 1.103 | 0 |
| 1.121 | 6.7 |
| 1.139 | 0 |
| 1.157 | 0 |
| 1.176 | 6.7 |
| 1.194 | 0 |
| 1.212 | 0 |
| 1.23 | 0 |
| 1.248 | 0 |
| 1.266 | 6.7 |
| 1.284 | 0 |
| 1.302 | 0 |
| 1.32 | 0 |
| 1.338 | 0 |
| 1.356 | 0 |
| 1.375 | 6.7 |
| 1.393 | 0 |
| 1.411 | 6.7 |
| 1.429 | 0 |
| 1.447 | 13.3 |
| 1.465 | 13.3 |
| 1.483 | 6.7 |
| 1.501 | 0 |
| 1.519 | 13.3 |
| 1.537 | 0 |
| 1.555 | 0 |
| 1.573 | 0 |
| 1.592 | 6.7 |
| 1.61 | 0 |
| 1.628 | 20 |
| 1.646 | 0 |
| 1.664 | 0 |
| 1.682 | 6.7 |
| 1.7 | 0 |
| 1.718 | 0 |
| 1.736 | 6.7 |
| 1.754 | 0 |
| 1.772 | 0 |
| 1.79 | 0 |
| 1.809 | 0 |
| 1.827 | 0 |
| 1.845 | 6.7 |
| 1.863 | 6.7 |
| 1.881 | 6.7 |
| 1.899 | 0 |
| 1.917 | 0 |
| 1.935 | 0 |
| 1.953 | 0 |
| 1.971 | 0 |
| 1.989 | 0 |
| 2.008 | 0 |
| 2.026 | 0 |
| 2.044 | 0 |
| 2.062 | 0 |
| 2.08 | 0 |
| 2.098 | 0 |
| 2.116 | 0 |
| 2.134 | 6.7 |
| 2.152 | 0 |
| 2.17 | 0 |
| 2.188 | 0 |
| 2.206 | 0 |
| 2.225 | 0 |
| 2.243 | 0 |
| 2.261 | 0 |
| 2.279 | 0 |
| 2.297 | 0 |
| 2.315 | 0 |
| 2.333 | 0 |
| 2.351 | 0 |
| 2.369 | 0 |
| 2.387 | 0 |
| 2.405 | 6.7 |
| 2.423 | 0 |
| 2.442 | 0 |
| 2.46 | 0 |
| 2.478 | 0 |
| 2.496 | 6.7 |
| 2.514 | 6.7 |
| 2.532 | 0 |
| 2.55 | 6.7 |
| 2.568 | 0 |
| 2.586 | 0 |
| 2.604 | 0 |
| 2.622 | 0 |
| 2.64 | 0 |
| 2.659 | 6.7 |
| 2.677 | 0 |
| 2.695 | 0 |
| 2.713 | 0 |
| 2.731 | 0 |
| 2.749 | 0 |
| 2.767 | 0 |
| 2.785 | 0 |
| 2.803 | 0 |
| 2.821 | 0 |
| 2.839 | 0 |
| 2.858 | 0 |
| 2.876 | 0 |
| 2.894 | 0 |
| 2.912 | 0 |
| 2.93 | 0 |
| 2.948 | 0 |
| 2.966 | 0 |
| 2.984 | 6.7 |
| 3.002 | 6.7 |
| 3.02 | 6.7 |
| 3.038 | 0 |
| 3.056 | 0 |
| 3.075 | 0 |
| 3.093 | 0 |
| 3.111 | 0 |
| 3.129 | 0 |
| 3.147 | 0 |
| 3.165 | 0 |
| 3.183 | 0 |
| 3.201 | 0 |
| 3.219 | 0 |
| 3.237 | 13.3 |
| 3.255 | 6.7 |
| 3.273 | 0 |
| 3.292 | 0 |
| 3.31 | 0 |
| 3.328 | 0 |
| 3.346 | 0 |
| 3.364 | 20 |
| 3.382 | 13.3 |
| 3.4 | 0 |
| 3.418 | 0 |
| 3.436 | 0 |
| 3.454 | 0 |
| 3.472 | 0 |
| 3.491 | 6.7 |
| 3.509 | 0 |
| 3.527 | 6.7 |
| 3.545 | 0 |
| 3.563 | 0 |
| 3.581 | 0 |
| 3.599 | 0 |

E

Intensity (cps)

| Time (min) | Vehicle |
| --- | --- |
| 0 | 6.7 |
| 0.018 | 6.7 |
| 0.036 | 13.3 |
| 0.054 | 6.7 |
| 0.072 | 6.7 |
| 0.09 | 6.7 |
| 0.109 | 6.7 |
| 0.127 | 13.3 |
| 0.145 | 0 |
| 0.163 | 0 |
| 0.181 | 6.7 |
| 0.199 | 0 |
| 0.217 | 13.3 |
| 0.235 | 0 |
| 0.253 | 0 |
| 0.271 | 6.7 |
| 0.289 | 13.3 |
| 0.307 | 0 |
| 0.326 | 0 |
| 0.344 | 6.7 |
| 0.362 | 0 |
| 0.38 | 20 |
| 0.398 | 0 |
| 0.416 | 13.3 |
| 0.434 | 6.7 |
| 0.452 | 0 |
| 0.47 | 0 |
| 0.488 | 6.7 |
| 0.506 | 0 |
| 0.525 | 0 |
| 0.543 | 0 |
| 0.561 | 13.3 |
| 0.579 | 26.7 |
| 0.597 | 6.7 |
| 0.615 | 0 |
| 0.633 | 0 |
| 0.651 | 6.7 |
| 0.669 | 0 |
| 0.687 | 6.7 |
| 0.705 | 6.7 |
| 0.723 | 6.7 |
| 0.742 | 0 |
| 0.76 | 0 |
| 0.778 | 0 |
| 0.796 | 0 |
| 0.814 | 0 |
| 0.832 | 0 |
| 0.85 | 6.7 |
| 0.868 | 6.7 |
| 0.886 | 6.7 |
| 0.904 | 0 |
| 0.922 | 0 |
| 0.94 | 0 |
| 0.959 | 0 |
| 0.977 | 0 |
| 0.995 | 0 |
| 1.013 | 0 |
| 1.031 | 6.7 |
| 1.049 | 0 |
| 1.067 | 0 |
| 1.085 | 0 |
| 1.103 | 0 |
| 1.121 | 0 |
| 1.139 | 6.7 |
| 1.157 | 0 |
| 1.176 | 6.7 |
| 1.194 | 0 |
| 1.212 | 0 |
| 1.23 | 0 |
| 1.248 | 0 |
| 1.266 | 13.3 |
| 1.284 | 6.7 |
| 1.302 | 6.7 |
| 1.32 | 0 |
| 1.338 | 0 |
| 1.356 | 0 |
| 1.375 | 0 |
| 1.393 | 13.3 |
| 1.411 | 0 |
| 1.429 | 0 |
| 1.447 | 0 |
| 1.465 | 0 |
| 1.483 | 6.7 |
| 1.501 | 0 |
| 1.519 | 6.7 |
| 1.537 | 13.3 |
| 1.555 | 6.7 |
| 1.573 | 0 |
| 1.592 | 53.3 |
| 1.61 | 26.7 |
| 1.628 | 0 |
| 1.646 | 26.7 |
| 1.664 | 0 |
| 1.682 | 6.7 |
| 1.7 | 26.7 |
| 1.718 | 13.3 |
| 1.736 | 0 |
| 1.754 | 0 |
| 1.772 | 6.7 |
| 1.79 | 13.3 |
| 1.809 | 13.3 |
| 1.827 | 6.7 |
| 1.845 | 0 |
| 1.863 | 6.7 |
| 1.881 | 0 |
| 1.899 | 6.7 |
| 1.917 | 13.3 |
| 1.935 | 6.7 |
| 1.953 | 0 |
| 1.971 | 0 |
| 1.989 | 0 |
| 2.008 | 0 |
| 2.026 | 0 |
| 2.044 | 0 |
| 2.062 | 6.7 |
| 2.08 | 0 |
| 2.098 | 6.7 |
| 2.116 | 0 |
| 2.134 | 0 |
| 2.152 | 6.7 |
| 2.17 | 0 |
| 2.188 | 6.7 |
| 2.206 | 6.7 |
| 2.225 | 6.7 |
| 2.243 | 6.7 |
| 2.261 | 0 |
| 2.279 | 0 |
| 2.297 | 0 |
| 2.315 | 6.7 |
| 2.333 | 0 |
| 2.351 | 6.7 |
| 2.369 | 0 |
| 2.387 | 0 |
| 2.405 | 0 |
| 2.423 | 6.7 |
| 2.442 | 0 |
| 2.46 | 0 |
| 2.478 | 0 |
| 2.496 | 0 |
| 2.514 | 0 |
| 2.532 | 0 |
| 2.55 | 0 |
| 2.568 | 13.3 |
| 2.586 | 0 |
| 2.604 | 0 |
| 2.622 | 0 |
| 2.64 | 0 |
| 2.659 | 6.7 |
| 2.677 | 0 |
| 2.695 | 0 |
| 2.713 | 6.7 |
| 2.731 | 6.7 |
| 2.749 | 0 |
| 2.767 | 0 |
| 2.785 | 0 |
| 2.803 | 0 |
| 2.821 | 0 |
| 2.839 | 0 |
| 2.858 | 13.3 |
| 2.876 | 0 |
| 2.894 | 0 |
| 2.912 | 6.7 |
| 2.93 | 0 |
| 2.948 | 6.7 |
| 2.966 | 0 |
| 2.984 | 0 |
| 3.002 | 0 |
| 3.02 | 20 |
| 3.038 | 0 |
| 3.056 | 0 |
| 3.075 | 6.7 |
| 3.093 | 0 |
| 3.111 | 0 |
| 3.129 | 0 |
| 3.147 | 0 |
| 3.165 | 13.3 |
| 3.183 | 0 |
| 3.201 | 0 |
| 3.219 | 0 |
| 3.237 | 0 |
| 3.255 | 0 |
| 3.273 | 0 |
| 3.292 | 0 |
| 3.31 | 13.3 |
| 3.328 | 6.7 |
| 3.346 | 0 |
| 3.364 | 0 |
| 3.382 | 13.3 |
| 3.4 | 0 |
| 3.418 | 6.7 |
| 3.436 | 0 |
| 3.454 | 0 |
| 3.472 | 0 |
| 3.49 | 6.7 |
| 3.509 | 0 |
| 3.527 | 0 |
| 3.545 | 0 |
| 3.563 | 0 |
| 3.581 | 0 |
| 3.599 | 6.7 |
